# Multi-omics analysis reveals the molecular response to heat stress in a “red tide” dinoflagellate

**DOI:** 10.1101/2022.07.25.501386

**Authors:** Katherine E. Dougan, Zhi-Luo Deng, Lars Wöhlbrand, Carsten Reuse, Boyke Bunk, Yibi Chen, Juliane Hartlich, Karsten Hiller, Uwe John, Jana Kalvelage, Johannes Mansky, Meina Neumann-Schaal, Jörg Overmann, Jörn Petersen, Selene Sanchez-Garcia, Kerstin Schmidt-Hohagen, Sarah Shah, Cathrin Spröer, Helena Sztajer, Hui Wang, Debashish Bhattacharya, Ralf Rabus, Dieter Jahn, Cheong Xin Chan, Irene Wagner-Döbler

**Affiliations:** The University of Queensland, School of Chemistry and Molecular Biosciences, Australian Centre for Ecogenomics, Brisbane, QLD 4072, Australia; Helmholtz-Center for Infection Research (HZI), Inhoffenstr. 7, 38124 Braunschweig, Germany; Institute for Chemistry and Biology of the Marine Environment (ICBM), Carl von Ossietzky University of Oldenburg, 26129 Oldenburg, Germany; Braunschweig Center for Systems Biology (BRICS), Technische Universität Braunschweig, Rebenring 56, 38106 Braunschweig, Germany; German Culture Collection for Microorganisms and Cell Cultures (DSMZ), Inhoffenstraße 7B, 38124 Braunschweig, Germany; Alfred Wegener Institute, Helmholtz Centre for Polar and Marine Research, Am Handelshafen 12, 27570 Bremerhaven, Germany; Helmholtz Institute for Functional Marine Biodiversity at the University of Oldenburg (HIFMB), Ammerländer Heersstraße 231, 26129 Oldenburg, Germany; Rutgers University, Department of Biochemistry and Microbiology, New Brunswick, NJ 08901, USA

**Author notes:** These authors contributed equally (data acquisition, analysis, and interpretation) (Katherine E. Dougan, Zhi-Luo Deng, Lars Wöhlbrand, Carsten Reuse). These authors contributed equally (design of the study, integration of all findings) (Cheong Xin Chan, Irene Wagner-Döbler). corresponding authors: Cheong Xin Chan, The University of Queensland, Brisbane, Australia, and Irene Wagner-Döbler, Technische Universität Braunschweig, Germany,.

## Abstract

“Red tides” are harmful algal blooms (HABs) caused by dinoflagellate microalgae that accumulate toxins lethal to other organisms, including humans *via* consumption of contaminated seafood. Increasingly frequent, HABs are driven by a combination of environmental factors including nutrient enrichment, particularly in warm waters. Here, we present the *de novo* assembled genome (~4.75 Gbp), transcriptome, proteome, and metabolome from *Prorocentrum cordatum*, a globally abundant, bloom-forming dinoflagellate. Using axenic algal cultures, we studied the molecular mechanisms that underpin response to temperature stress, which is relevant to current ocean warming trends. We discovered a complementary interplay between RNA editing and exon usage that regulates the expression and functional diversity of biomolecules, reflected by reduction in photosynthesis, central metabolism, and protein synthesis. Our multi-omics analyses uncover the molecular response to heat stress in an important HAB species, which is driven by complex gene structures in a large, high-G+C genome, combined with multi-level transcriptional regulation.

## Introduction

Harmful algal blooms (HABs) result from highly accelerated microalgal growth that is often triggered by increasing water temperature, light intensity, and/or available nutrients. HABs often lead to oxygen depletion and toxin accumulation, causing significant losses to fisheries and aquaculture industries (~USD 8B annual losses globally^1, 2^). Among HABs, the increasingly frequent “red tides” are caused by bloom-forming dinoflagellate microalgae, such as species of *Alexandrium*, *Amphidinium* and *Prorocentrum*^3, 4^. Habitat expansion of bloom-forming dinoflagellates has been linked to warming oceans and global climate change^5^.

Dinoflagellates are an ancient and highly diverse plankton group within the Alveolata, encompassing free-living, bloom-forming, parasitic, and symbiotic taxa^6, 7^, with most species being mixotrophs (*i.e*. they combine phototrophic and heterotrophic modes of energy generation) or heterotrophs^8^. The photosynthetic machinery of dinoflagellates powers the biological carbon pump of oceans, which is essential for lowering the global carbon budget. With increasing water temperature, aquatic microbes are exposed to several cellular challenges at their upper tolerance level, including with high metabolic rates and membrane fluidity, while maintaining photosynthetic efficiency. For dinoflagellates, the formation of large blooms and the maintenance of beneficial symbioses with corals and other organisms are all impacted by increasing water temperature^5, 9^. Transcriptome studies of toxic dinoflagellates, *e.g*.^10–12^, had revealed stress-related gene functions including metabolism and cell signaling. However, the molecular regulatory and evolutionary mechanisms that underlie the heat-stress response in HAB forming species remain little understood. This is primarily explained by the lack of high-quality genome data from these dinoflagellates, which may be up to ~200 Gbp in size^13, 14^ and exhibit features atypical of eukaryotes^15, 16^. Thus far, genome studies^17–21^ have targeted members of the family Symbiodiniaceae which form coral symbiosis and their free-living relatives in the genus *Polarella* (genome sizes ≤ 3Gbp). These analyses reveal extensive sequence divergence and lineage-specific innovations with respect to putative gene functions. A single draft genome assembly exists for HAB-forming dinoflagellates, from *Amphidinium gibbosum* (~6.4 Gbp)^22^, which was used to study metabolic and toxin biosynthesis functions vis-à-vis nutrient deprivation. Past studies lack proteome and metabolome data, which are necessary to elucidate the molecular mechanisms that underpin gene-expression regulation. Although some muti-omics data were recently generated from three Symbiodiniaceae species^23^, these results are not relevant to distantly related HAB species, given the high divergence that exists among dinoflagellate genomes^21, 24^.

Relevant to our study, transcriptional regulation is minimal in dinoflagellates^12, 16^ with only a handful of known transcriptional regulators^25^, and chromosomes existing in a permanently condensed, liquid crystalline state^26^. Initial studies^27, 28^ suggested that most dinoflagellate genes are constitutively expressed regardless of growth conditions, particularly of shock treatments, but more-recent research suggests a potentially important role for differential gene regulation in these species^29^. *Trans*-splicing of a conserved spliced-leader sequence in nuclear genes has been described^30, 31^. Editing of mRNAs occurs for both organelle- and nuclear-encoded genes^28, 32^, suggesting a role for this mechanism in generating physiological flexibility.

Among bloom-forming dinoflagellates, *Prorocentrum cordatum* (formerly *Prorocentrum minimum*)^33, 34^ is an invasive, potentially toxic species that is found globally^35^ and is regularly detected in the North Atlantic^36^. The tolerance of *P. cordatum* to a wide range of salinities and temperatures facilitates its increased bloom frequency^33, 34^. Here, we present the genome and multi-omics data from *P. cordatum*, targeting the algal heat stress response in axenic cultures. Our results provide an integrated view of how a HAB-forming species may respond to ocean warming induced by global climate change.

## Results

### *P. cordatum* genome reveals hallmarks of bloom-forming dinoflagellates

We generated a *de novo* haploid, repeat-rich genome assembly from *P. cordatum* CCMP1329 (4.15 Gb, scaffold N50 length = 349.2 Kb; Extended Data Table 1 and Fig. 1a). Compared to five representative genomes of dinoflagellates^21, 37–40^ from diverse ecological niches (Supplementary Note), *P. cordatum* has the highest G+C content in the genome sequences (mean 59.7%; Fig. 1c and Extended Data Table 1) and in protein-coding genes (mean 65.9%; Fig. 1d and Extended Data Table 2), compared to the moderate G+C content observed in the bloom-forming *A. gibbosum*^22^ and the free-living *Polarella glacialis*^37^ (Extended Data Table 1). The larger genome of *P. cordatum* encodes more protein-coding genes with longer introns than do the other species (Fig. 1e, Extended Data Table 2). These introns are enriched in introner elements, *i.e*., introns that contain simple inverted repeats at both ends (Supplementary Table 1), suggesting a prevalence of these elements in the genomes of free-living dinoflagellates (Supplementary Table 2 and Supplementary Note).

**Fig. 1.**
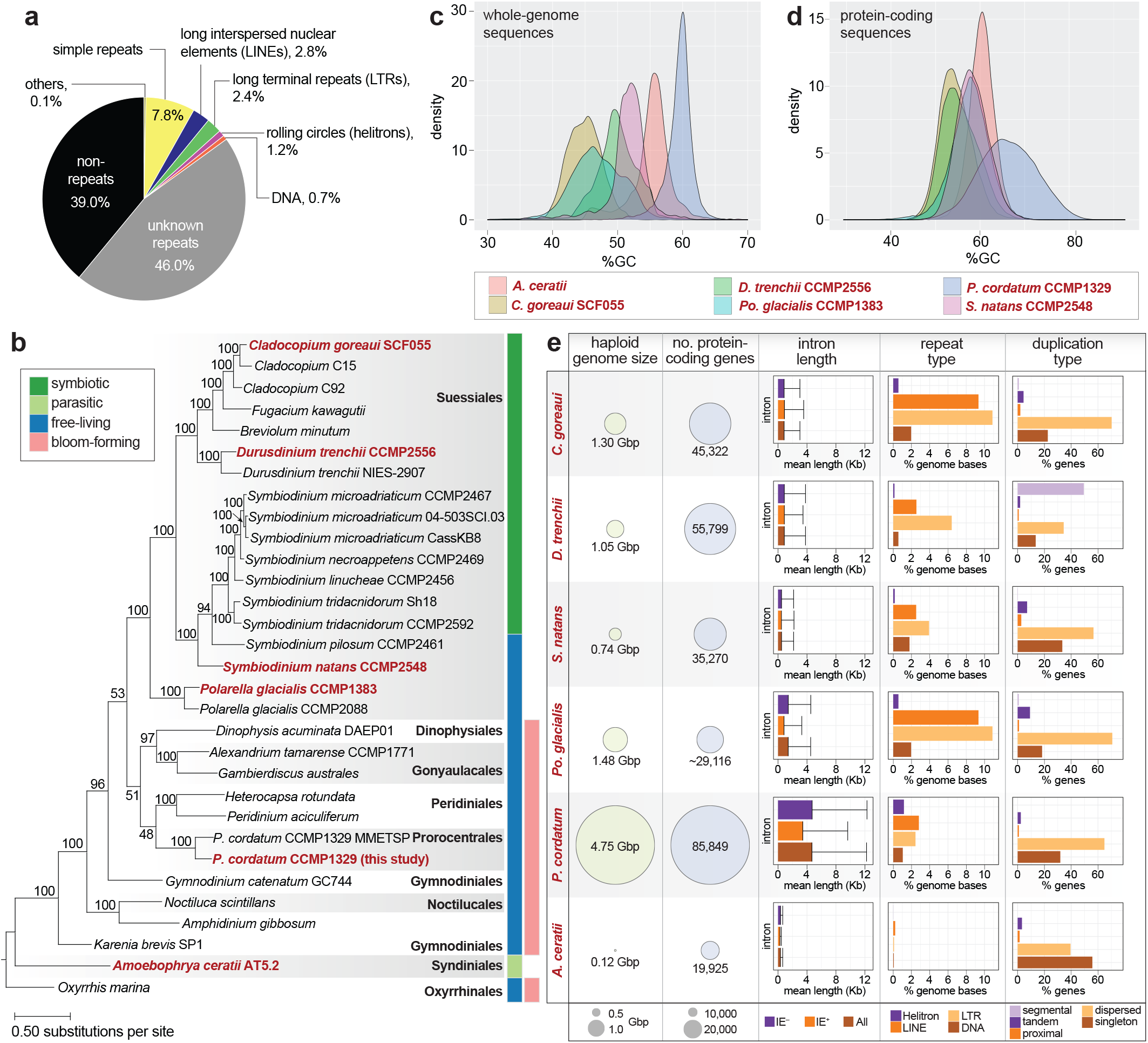
Genome features of *P. cordatum*. (a) Distribution of repeat types in the *P. cordatum* genome. (b) Maximum likelihood tree inferred using 3,507 strictly orthologous, single-copy protein sets among 31 dinoflagellate taxa, with ultrafast bootstrap support (based on 2000 replicate samples) shown at each internal node; unit of branch length is number of substitutions per site. The ecological niche for each taxon is shown on the right of the tree. The five representative taxa and *P. cordatum* from this study are highlighted on the tree in red text. Distribution of G+C content for (c) whole-genome sequences and (d) protein-coding sequences relative to the other five representative genomes. (e) Genome and gene features of *P. cordatum* relative to the other five taxa, showing haploid genome size estimated based on sequence data, number of protein-coding genes, intron lengths, and separately for introns that contain introner elements (IE^+^), and those that lack these elements (IE^−^), known repeat types, and types of duplicated genes.

We predicted 85,849 gene models in *P. cordatum*, 41,078 (47.8%) of which were annotated using a stringent approach (Supplementary Tables 3 and 4, and Supplementary Note); about half (52.2%) are assigned as “dark”, coding for functions yet to be discovered^41^. Based on the relative abundance of annotated Gene Ontology (GO) terms (Fig. 2a) in *P. cordatum* genes, we observed more-abundant functions related to metabolism, cell signaling, transmembrane transport, and stress response (Supplementary Note). Whereas most genes are unidirectionally encoded (Supplementary Fig. 3), a substantial proportion (64.8% of 85,849) are dispersed duplicates (Fig. 1e and Supplementary Table 5), suggesting that most duplication events occurred independently; alternatively, collinearity of duplicated blocks was disrupted by extensive rearrangements, due in part to the abundant transposable elements. We found significantly enriched (*p* ≤ 0.01) gene functions in the distinct types of gene duplicates (Fig. 2b), *e.g*. transmembrane transport and organelle assembly among the dispersed duplicates, compared to metabolic processes (*e.g*. tricarboxylic acid cycle [TCA]) and binding of biomolecules/ions among the tandem duplicates (Supplementary Table 6). These results demonstrate that distinct duplication modes have shaped the evolution of *P. cordatum* genes and their functions. Moreover, 47 genes were potentially acquired *via* horizontal transfer from uncultivated marine prokaryotes with functions related to structural conversion of amino acids and biosynthesis of metabolites (Supplementary Table 7, Supplementary Fig. 4, and Supplementary Note).

**Fig. 2.**
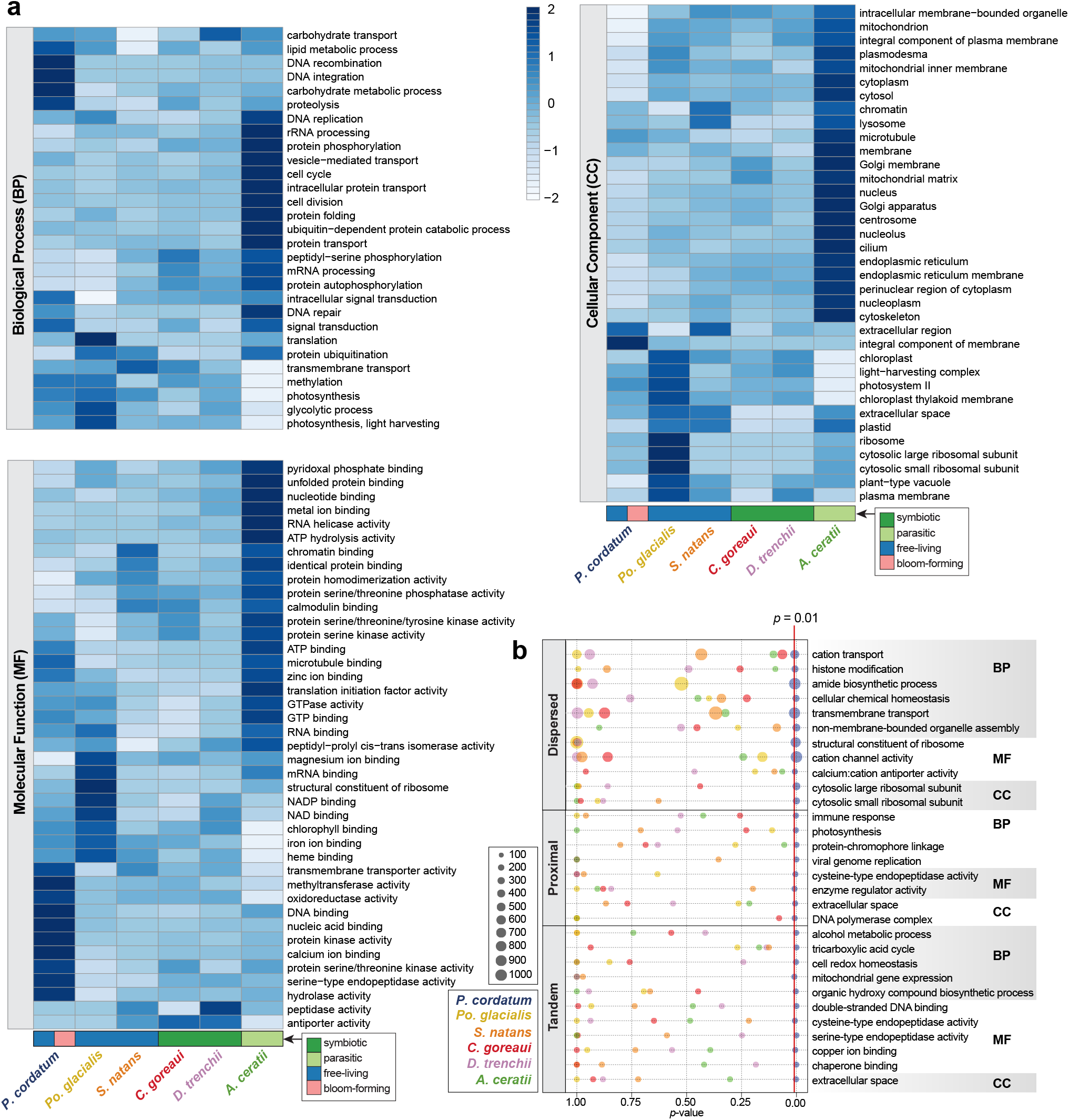
Gene functions encoded in the *P. cordatum* genome. (a) Gene functions encoded in the genome of *P. cordatum* and the other five representative taxa based on relative abundance of Gene Ontology (GO) terms per genome, shown for categories of Biological Process (BP), Molecular Function (MF) and Cellular Component (CC). The ecological niche for each taxon is shown at the bottom of the heatmaps. (b) GO terms that are significantly enriched (*p* < 0.01) among genes for each duplication type (*i.e*. dispersed, proximal and tandem) in *P. cordatum*, relative to *p*-values observed for the other taxa, and the associated number of genes for each GO term.

### Integrated multi-omics analysis of heat-stress responses specific to *P. cordatum*

To investigate the heat-stress response in *P. cordatum*, axenic cultures were grown in defined media at the optimal temperature (20°C) before they were exposed to either 26°C or 30°C (Fig. 3a). We observed similar growth rates (0.33-0.47 day^−1^ and 0.47-0.68 doubling rate day^−1^) under all three conditions, but relative to the final cell density observed at 20°C, algal biomass was reduced to 62% and 41% at 26°C and 30°C, respectively (Fig. 3b). Stable cell numbers over two weeks at stationary phase at both elevated temperatures indicate the tolerance of *P. cordatum* to heat stress. Transcriptome, proteome, and metabolome data (Supplementary Tables 8, 9, 10 and 11) were generated from cells harvested independently at exponential (Ex) and stationary (St) growth phases in the three temperature conditions (Fig. 3b; see Methods).

**Fig. 3.**
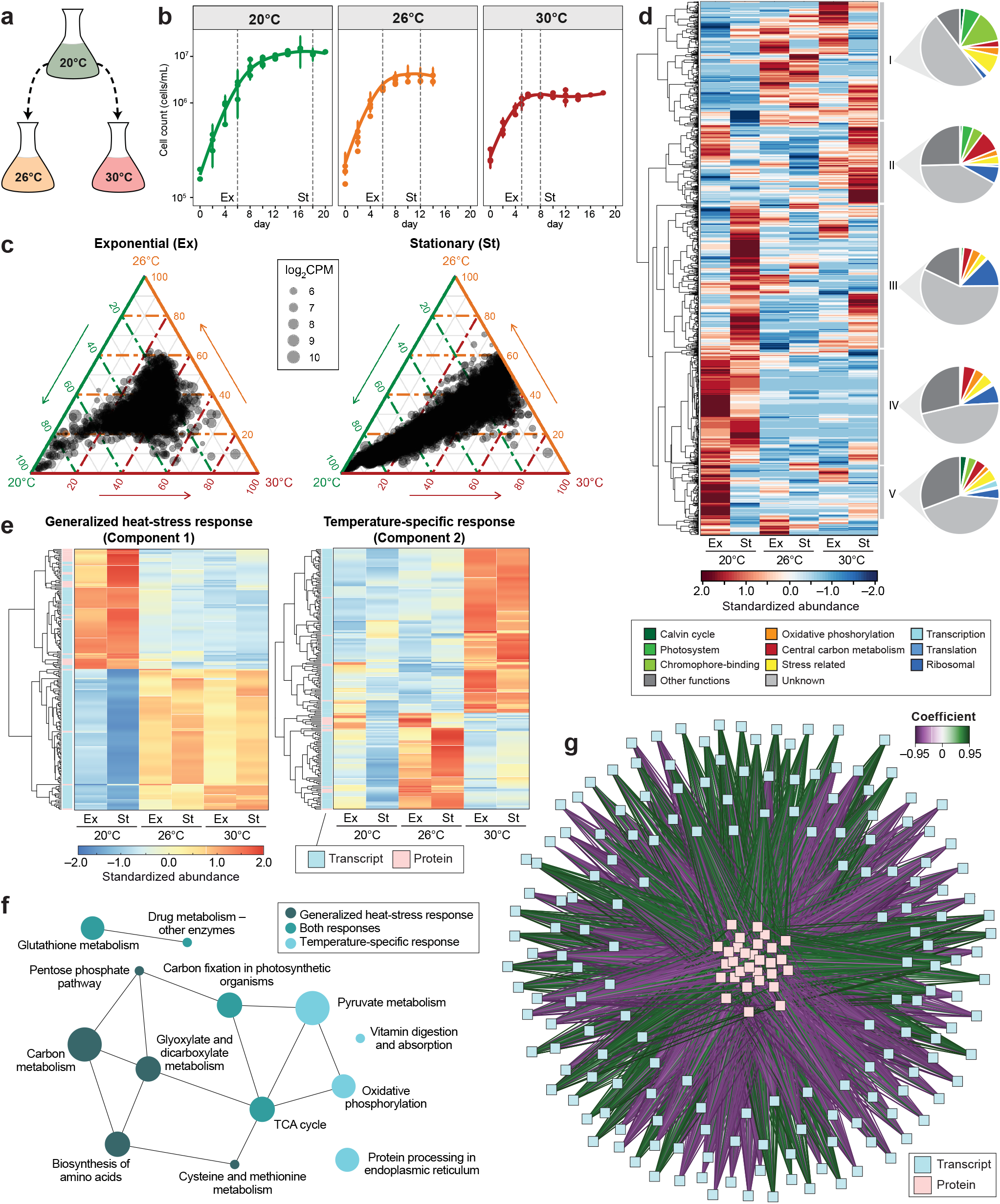
Integrated analysis of the transcriptome and proteome response of *P. cordatum* to heat stress. (a) Experimental design. (b) Growth of *P. cordatum* at 20°C, 26°C and 30°C. Collection of cells for multi-omics analysis is indicated by dashed vertical lines (Ex: exponential, St: stationary phase). (c) Ternary plots of highly expressed gene models with mean log_2_(count per million) >5 in response to temperature and growth phase (8,593 transcripts in each plot). (d) Clustering of 2,098 differentially abundant proteins in response to temperature and growth phase. Abundances of proteins were calculated from standardized peptide counts. (e) Heatmap of transcripts and proteins showing significant correlations for generalized heat stress response (component 1) and temperature-specific response (component 2). (f) Over-represented KEGG pathways in the networks of generalized and temperature specific heat-stress response. (g) DIABLO network of generalized heat stress response (component 1) revealing positive and negative correlations (coefficient ≥0.7) between transcripts and proteins.

Analysis of the transcriptome data (1.96 Gb, ~110 million reads per sample; Supplementary Table 8) from 18 samples (6 conditions × 3 replicates) using Principal Component Analysis revealed a clear separation between 20°C and the higher temperatures in both Ex and St phases (Supplementary Fig. 5 and Supplementary Table 9), although gene expression changes are less pronounced in the Ex phase (Fig. 3c). Analysis of soluble and membrane protein fractions yielded 2,098 proteins, of which 1,032 were of unknown function; 244 are unique to *P. cordatum*. The 68 chromophore-binding (antennae) proteins of the photosystem comprised the largest group, accounting for 70.6% of detected peptides. We found 779 proteins with significantly changed abundance at 26°C or 30°C compared to 20°C, with functions related to photosynthesis, energy generation, and central metabolism (Fig. 3d, Supplementary Fig. 6 and Supplementary Table 10). Metabolome analysis (see Methods) yielded 173 compounds, of which 73 could be identified (Extended Data Fig. 1 and Supplementary Table 11). Fifty-four metabolites displayed significantly changed abundances (*p* < 0.05) in response to growth phase and temperature, particularly those involved in central metabolism and amino acid biosynthesis (Supplementary Table 11).

To identify the molecular response in *P. cordatum* to heat stress, we integrated the transcriptome and proteome data using DIABLO^42^ to identify shared multi-omics signatures of the Ex and St phases. This analysis revealed two types of heat stress response: a generalized response (component 1) with abundance changes common to both elevated temperatures (26°C and 30°C), and a temperature-specific response (component 2) with abundance changes specific to 26°C or to 30°C (Fig. 3e); transcripts and proteins comprising these two components are shown in Supplementary Table 12. KEGG pathways for carbon metabolism such as the pentose phosphate pathway, glyoxylate and dicarboxylate metabolism, the biosynthesis of amino acids, and the metabolism of cysteine and methionine were enriched in the generalized response (component 1; Fig. 3f and Extended Data Fig. 2). In contrast, oxidative phosphorylation, protein processing in endoplasmic reticulum, vitamin digestion and absorption, and pyruvate metabolism pathways were enriched in the temperature-specific response (component 2; Extended Data Fig. 3). Our results also reveal both positive and negative correlations of expression between transcripts and proteins in component 1 (Fig. 3g) and component 2 (Supplementary Fig. 7). For instance, gene expression of glutamine synthetase was positively correlated to the expression of chlorophyll *a-b* binding protein and the light-harvesting complex I LH38 proteins, whereas it was negatively correlated to protein expression of pyruvate dehydrogenase, and the sulfate and formate transporters.

### Heat-induced multi-omics modulation of central metabolism in *P. cordatum*

Given the lower biomass observed at elevated temperatures (Fig. 3b), we studied the recovery of biomolecules specific to three metabolic modules that drive growth and primary production in *P. cordatum*: photosynthesis, central metabolism, and oxidative phosphorylation. A global visualization of the relevant expressed transcripts (753) and proteins (278) revealed differential expression at elevated temperatures, with a marked increase in photosynthesis proteins but a decreased abundance of proteins related to central metabolism and oxidative phosphorylation (Fig. 4a). This result is supported by reduction in the accumulation of 38 metabolites associated with central metabolism at elevated temperatures.

**Fig. 4.**
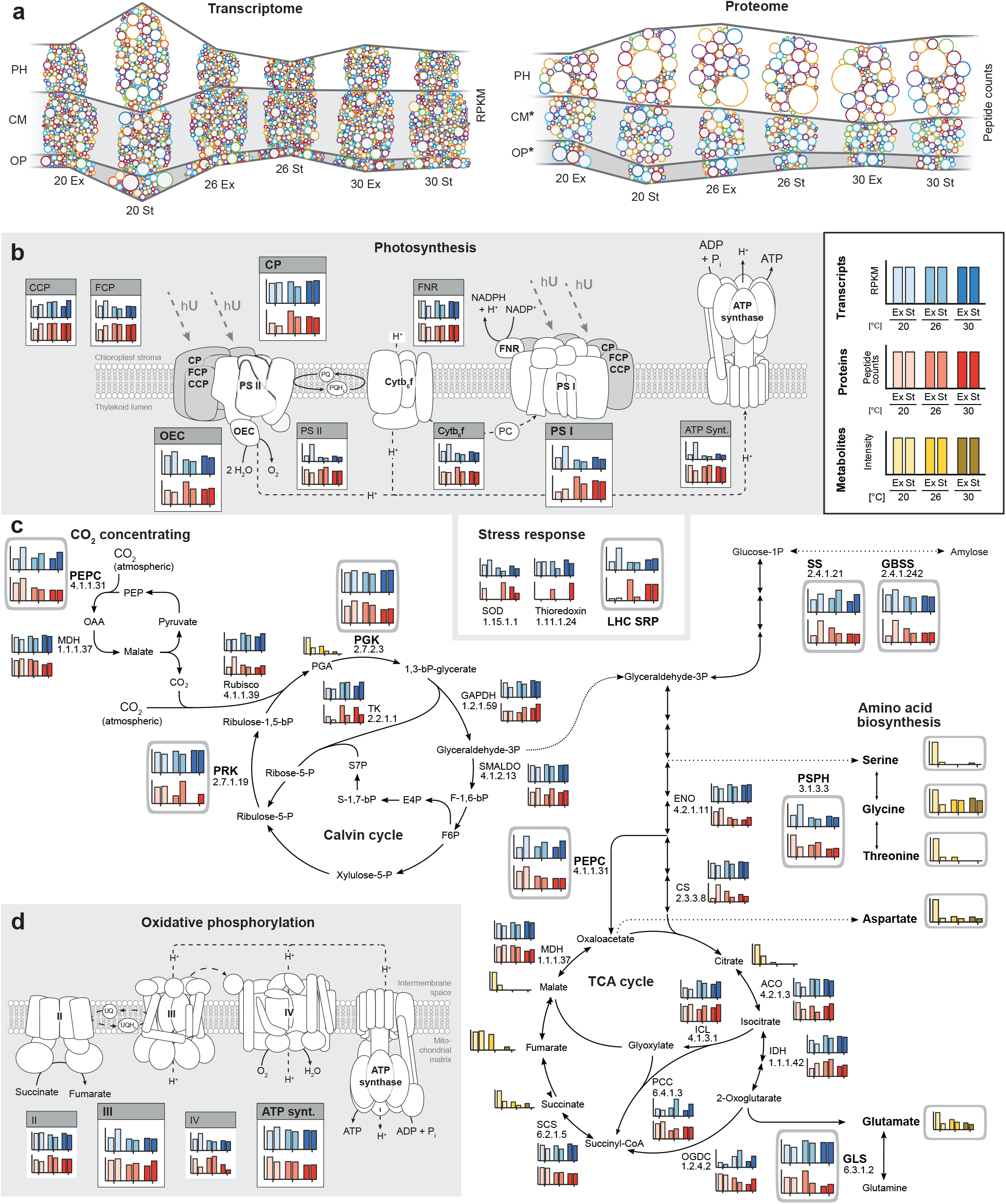
Heat stress response of central modules of energy and carbon metabolism in *P. cordatum*. (a) Temperature-dependent dynamics of sub-transcriptomes (left) and sub-proteomes (right) associated with photosynthesis (PH), central metabolism (CM), and oxidative phosphorylation (OP). Colored circles represent individual transcripts and proteins, respectively, with their areas proportional to the determined abundances. For the proteome, heights of the CM- and OP-bands (marked with an asterisk) were magnified 10-fold to allow easier comparison with the PH-band. Expression of transcripts, proteins, and metabolites is shown for functions specific to (b) light reaction of photosynthesis, showing CCP, carotenoid/chlorophyll-binding protein; CP, chlorophyll-binding protein; FCP, fucoxanthin/chlorophyll-binding protein; FNR, ferredoxin:NADP oxidoreductase; OEC, oxygen evolving complex; and PS, photosystem; (c) central metabolism including CO_2_-concentrating Calvin cycle, central carbon metabolism, and selected biosynthesis of amino acids, showing PEPC, phosphoenolpyruvate carboxylase; GBSS, granule-bound starch synthase (NDP-glucose-starch glycosyltransferase); GLS, glutamine synthetase; PGK, phosphoglycerate kinase; PRK, phosphoribulokinase; PSPH, phosphoserine phosphatase; and SS, starch synthase; and (d) oxidative phosphorylation. A detailed scheme is presented in Extended Data Fig. 4.

The detected components of the photosynthetic electron transport chain (PETC) showed varying abundance profiles (Fig. 4b). Whereas chlorophyll-binding proteins (CPs), light harvesting complex (LHC), stress related proteins and photosystem I (PS I) proteins increased at elevated temperature, subunits of the chloroplast ATP synthase did not change. Notably, PS II, and the associated oxygen evolving complex (OEC) and enzyme components of the response to reactive oxygen species (Supplementary Tables 13 and 14), appeared unchanged, but complex IV and mitochondrial ATP synthase of oxidative phosphorylation showed reduced abundance at higher temperature (Fig. 4d). Taken together, during heat stress *P. cordatum* faces energy deprivation arising from a less efficient PETC, which should directly impact protein synthesis and central metabolism. Accordingly, amino acid synthesis was reduced, demonstrated by decreased levels of phosphoserine phosphatase (PSPH) and glutamine synthetase (GLS), amino acids (*e.g*. serine and glutamate), and TCA-cycle intermediates (*e.g*. succinate) (Fig. 4c; detailed reconstruction of central metabolism in Extended Data Fig. 4). These results suggest lower levels of enzymes involved in concentrating CO_2_ (phosphoenolpyruvate carboxylase [PEPC]) and in ATP-consuming reactions of the Calvin cycle, *e.g*. phosphoribulokinase (PRK) and phosphoglycerate kinase (PGK) (Fig. 4c). The generally unchanged profile of CO_2_-fixing ribulose 1,5-bisphosphate carboxylase (Calvin cycle) may be misleading in this context, because activity of this enzyme is known to decrease at higher temperatures^43^. Enzymes involved in the synthesis of amylose, *e.g*. starch synthase (SS) and granule-bound starch synthase (GBSS), remained stable (Fig. 4c). In contrast to the declining levels of TCA metabolites and amino acids, some carbohydrates and fatty acids increased at elevated temperature, suggesting the recycling metabolites and re-organization of cellular processes and structural elements (*e.g*. lipids) to protect cells from heat stress, or as a countermeasure against increased membrane fluidity^44^. These results demonstrate the severe impact of heat stress on essential metabolic processes that attenuated *P. cordatum* growth.

### Transcriptional dynamics under heat stress

The transcriptome of *P. cordatum* showed extensive differentially expressed genes (DEGs) under heat stress: 2,142 in the Ex phase, and 22,924 in the St phase (Supplementary Table 15), and we observed no strong evidence of batch-effect biases (Supplementary Fig. S5a). The most extensive difference (14,322 DEGs) was observed between 20°C and 26°C in the St phase (*i.e*. a chronic heat-stress response), of which 11,155 were also observed between 20°C and 30°C (Supplementary Fig. S5b). Among 2,368 homologous gene sets that contain two dispersed gene copies, 368 contain both copies as DEGs, of which 341 shared a similar expression profile (176 + 165 in Fig. 5a). Of the 994 sets containing three dispersed gene copies, 982 (98.8%) consist of two or more copies showing similar expression patterns (Fig. 5b). Moreover, among 949 sets containing ≥ 3 dispersed copies as DEGs, 789 (83.1%) contain 80% of such copies that were expressed similarly (Fig. 5c). The conserved expression patterns for this subset of dispersed duplicates hint toward gene dosage as a driver of the transcriptome in *P. cordatum* despite the lack of evidence for whole-genome duplication^38^; see Supplementary Note and Supplementary Fig. 8 for more detail.

**Fig. 5.**
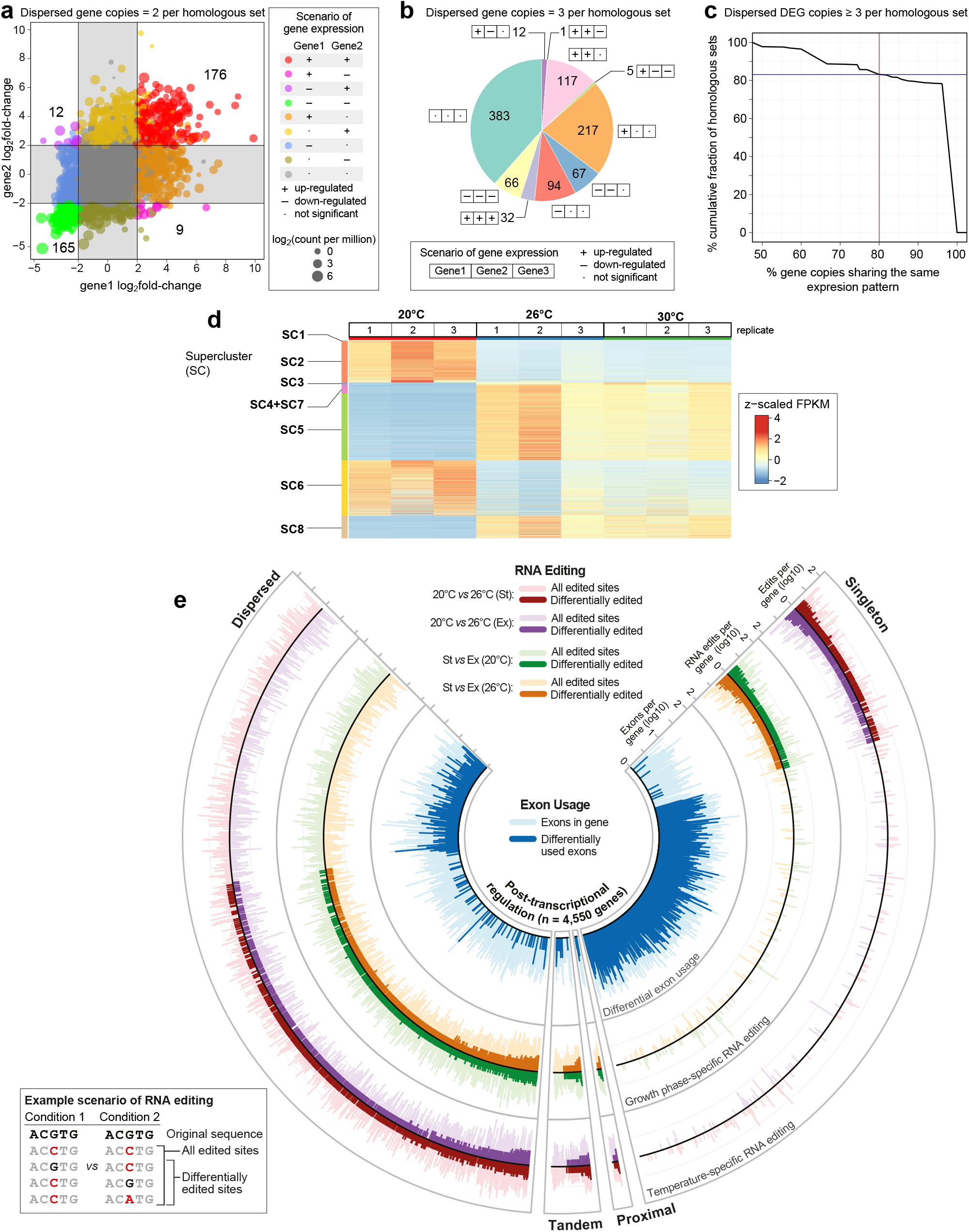
The transcriptome landscape under heat stress. The expression pattern of dispersed duplicated genes under treatments comparing 26°C against 20°C at St phase, shown for homologous sequence sets that contain (a) two and (b) three copies, in three possible outcomes: up-regulated (+), down-regulated (−), and not significant (·). (c) Proportion of dispersed DEGs that share similar expression pattern in homologous sets containing ≥3 of such copies. (d) Heatmap and clusters of gene-expression pattern across triplicate samples at 20, 26 and 30°C in the St phase, showing eight superclusters (SCs). Centered log_2_(FPKM+1) values are shown. (e) Post-transcriptional regulation in *P. cordatum*, shown for 4,550 genes in distinct duplication modes (counter-clockwise): dispersed, tandem, proximal, and singleton. Features shown from the inner-most to the outer-most circle: differential exon usage (blue), differential editing of mRNA per gene in response to growth phase (green/brown), and differential mRNA editing per gene in response to temperature (red/purple). In each circle, a bar in a light shade indicates the number of corresponding features identified in a gene, a bar in a dark shade indicates the number of statistically significant features in a gene. The bottom-left legend shows the distinct scenarios for identifying sites for mRNA editing in all transcripts *versus* differentially edited sites.

Based on the expression pattern across the three temperature conditions, DEGs were grouped into eight superclusters (SCs) independently at Ex (Extended Data Fig. 5a, Supplementary Table 16) and St phase (Fig. 5d, Extended Data Fig. 5b, Supplementary Table 17). Of the 22,924 DEGs in St phase, 11,647 (50.8%) showed an increase in expression from 20, 26 to 30°C (SC3–5, SC7 and SC8), 4,853 (21.2%; SC1 and SC2) showed a gradual decrease, and 6424 (28.0%; SC6) showed little changes (Fig. 5d, Extended Data Fig. 5b). Enrichment analysis of GO terms revealed that at 26°C relative to 20°C in St phase, functions related to inositol oxygenase activity, cellulase activity, ATP-binding, and metal ion binding were upregulated (Extended Data Fig. 6a, Supplementary Table 17), whereas those related to ribosome, rRNA binding, translation, transmembrane transporter, photosystem II, and photosynthesis light reaction were downregulated (Extended Data Fig. 6b). At Ex phase, functions related to structural constituents of cytoskeleton and microtubule were downregulated at 26°C (Supplementary Table 18). These results are consistent with the observation of enriched metabolic pathways (Extended Data Fig. 6, Supplementary Tables 19 and 20), and GO terms among the SCs (Extended Data Fig. 5c). Interestingly, among 13 polyketide synthase I genes, likely involved in biosynthesis of most dinoflagellate toxins^45^, five were upregulated and none were downregulated under heat stress (Supplementary Table 21). These results highlight the dynamic transcriptional response of *P. cordatum* to heat stress, which is greater than previously reported for any dinoflagellate; see Supplementary Note for more detail.

### Post-transcriptional regulation *via* the complementary interplay of RNA editing and exon usage

The integration of genome and transcriptome data revealed two modes of post-transcriptional regulation in *P. cordatum* that may generate protein diversity (and functions): (a) alteration of a single base in mRNA (*i.e*. differential RNA editing), and (b) alternative splicing that leads to preferential usage of exons (*i.e*. differential exon usage: DEU); see Methods. We found evidence of post-transcriptional regulation in 4,550 genes (Fig. 5e): 2,744 genes (involving 4,410 sites) with differential RNA editing, 1,755 genes (involving 5,837 exons) with DEU (Supplementary Table 22), and 51 genes (involving 93 sites and 72 exons) with both modes of regulation. The number of mRNA edits were similar under all conditions, but the type of edit and overlap between edited sites was condition-specific (Fig. 5e, Supplementary Table 23).

Interestingly, the relatively few (51) genes with both RNA editing and DEU in *P. cordatum* suggests that post-transcriptional regulation of a gene tends to involve only one of these two modes: the editing of RNA was identified predominantly in genes for which no DEU was identified, and *vice versa* (Fig. 5e); singleton genes showed a greater extent of DEU in contrast to the dispersed duplicates that exhibited a greater extent of RNA editing. The 51 genes displaying both modes of regulation encode key photosynthetic and stress-related functions, such as chloroplastic ATP synthase subunits (β and ε subunits), ferredoxin, photosystem I reaction center subunit XI, and heat shock 70 kDa protein. The complementary role of these two pathways for post-transcriptional regulation in central pathways (*e.g*. photosynthesis) may represent a key mechanism for generating functional diversity of the proteome in *P. cordatum* (and potentially more broadly in dinoflagellates) to enable quick acclimation to environmental changes.

### Regulation of multi-protein-coding gene variants as a concerted heat-stress response

To assess the impact of RNA editing and exon usage on the heat-stress response, we focused on genes that undergo both of these processes. An example is the gene coding for heat-shock protein 70 kDa (HSP70), a ubiquitous chaperone associated with stress responses. Of the 16 *P. cordatum* gene models that putatively code for HSP70 of varying lengths (574–2,575 amino acids), one (s12246_g74608) encodes multiple full-length copies of the functional protein. The protein-coding sequence is organized in multiple sub-regions, in which each sub-region encodes the same full-length protein; we follow Shi et al.^46^ and define such a sub-region within a gene model as a coding unit (CU), and the entire sequence as a multi-CU gene model. These CUs are separated by spacer sequences that are collectively transcribed as a single transcript (Extended Data Fig. 7). This is a form of polycistronic transcription in dinoflagellates^30^, whereby multiple (different) genes are co-transcribed, and each CU has a termination codon for translation (Supplementary Fig. 9). HSP70 genes in other unicellular eukaryotes such as *Leishmania* and trypanosomes are also polycistronically transcribed, with the 3’-UTR and regions downstream of the protein important for translational regulation^47–49^. Polycistronic transcripts in dinoflagellates are thought to be converted into monocistronic sequences *via trans*-splicing of a conserved 22-nucleotide dinoflagellate spliced leader (dinoSL) sequence^30^. We found 4.95% of *P. cordatum* transcripts to contain dinoSL (Supplementary Table 24) involving 17,214 (20.1% of 85,849) gene models, including those encoding HSP70, lending further support to this idea (Supplementary Note).

In *P. cordatum*, the multi-CU gene model is the main contributor to the heat stress response of HSP70 along with one single-CU gene model in cluster 1 (Fig. 6a). Exon 1 (CU-1) of the multi-CU gene displayed a unique pattern of RNA editing (Fig. 6b), with contrasting RNA edits specific to growth phase and temperature, including multiple sites upstream of the transcriptional start site (Supplementary Table 23). In this gene, exons 1, 3, 5 and 7 constitute four CUs, with exons 9 and 10 encoding a partial CU (Fig. 6b, Extended Data Fig. 7). The usage of exon 3 (CU-2) was biased towards a higher abundance at 20°C of St phase, in combination with multiple specific intronic RNA-edited sites downstream of this exon. Interestingly, the HSP70 encoded by this exon lacks a disordered C-terminus region involved in protein-protein interactions in many species.

**Fig. 6.**
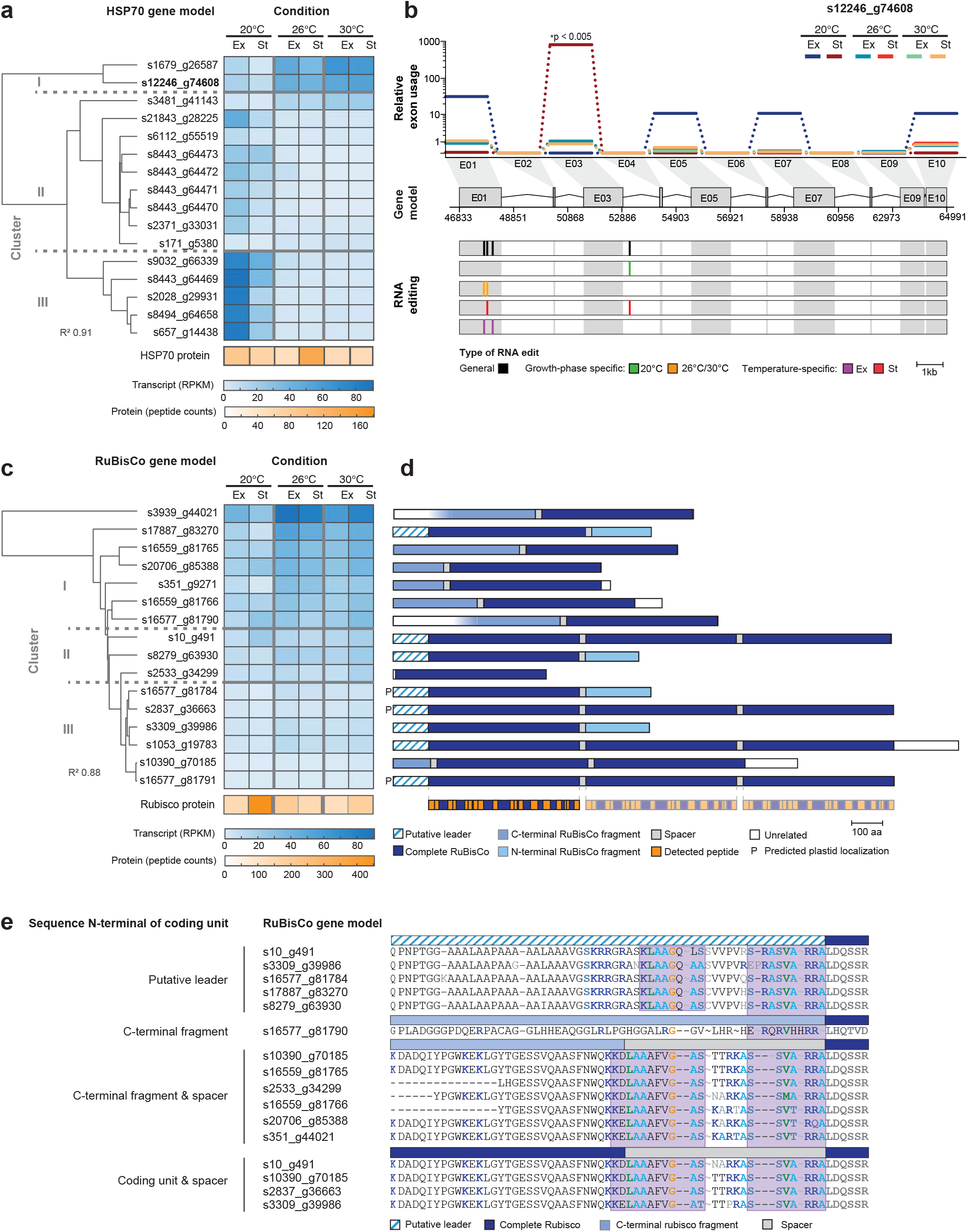
Expression and structure of multi-protein-coding genes in *P. cordatum*. The example of HSP70 is shown for (a) clustering of distinct gene models based on gene-expression pattern in a heatmap across growth conditions, with cumulative abundance of transcripts and proteins indicated at the bottom, and (b) the relative exon usage, CDS structure, and mRNA editing of gene model s12246_g74608 that harbors multiple CUs. Exon (E) is connected by introns (kinked line); each of E01, E03, E05 and E07 encodes a complete HSP70, E09 and E10 each encodes N-/C-terminal fragments, whereas the other exons encode spacers. The example of RuBisCo is shown for (c) clustering of gene models based on gene-expression pattern in a heatmap across growth conditions, with cumulative abundance of transcripts and proteins indicated at the bottom, and (d) protein structures corresponding to complete CUs, N- and C-terminal fragments, spacer, and putative leader sequences, with detected peptides indicated at the bottom. (e) Aligned upstream regions of 16 complete RuBisCo proteins of *P. cordatum*, including leader sequence and representative spacer sequences where available. Basic residues are marked in dark blue and conserved regions are highlighted in a purple background.

To assess expression variation of multi-CU *versus* single-CU genes in response to heat stress, we focused on the multi-CU gene model that encodes the CO_2_-fixing enzyme, ribulose-1,5-bisphosphate carboxylase (RuBisCo). Of the 32 *P. cordatum* gene models that putatively encode RuBisCo (of sizes 163 to 5,197 amino acids; Supplementary Fig. 10), one-half encode fragments of the protein, whereas the remainder are organized in single or multiple CUs (Fig. 6d). Distinct from polycistronic transcripts, here, each CU in a gene model does not have a termination codon for translation (Supplementary Fig. 9). Earlier studies of two *Prorocentrum* species^46, 50^ revealed a single transcript encoding three or four consecutive CUs, and the unprocessed protein product is thought to be transported into the plastid where it is split into separate, active proteins^46^. Here, we found gene models that encode three CUs (4), two CUs (1), and a single CU (11; Fig. 6d); dinoSL-containing transcripts were identified for some of these, *e.g*. in the three-CU-encoding s10_g491 (Supplementary Table 24).

Transport of RuBisCo to the plastid is facilitated by transit peptides including a leader sequence encoded upstream of the gene; these sequences are highly diverged among photoautotrophs, and no consensus sequence has been described for dinoflagellates^51, 52^. We found, in the upstream region of eight gene models (Fig. 6d), the predicted leader sequence from the closely related *Prorocentrum donghaiense*^46^. We identified two conserved sequence motifs in the leader sequence, the N-terminal region of proteins that lacked a leader sequence, and the spacer region between two CUs (Fig. 6e). The two sequence motifs are moderately hydrophobic and rich in basic amino acid residues (*i.e*. arginine or lysine) as expected for plastid transit peptides^52^. The conserved sequence regions indicate that proteins coded by these single- and multi-CU genes in *P. cordatum* can be transported into the plastid. Highly similar sequence motifs described in *P. donghaiense* and other dinoflagellates (Supplementary Fig. 11) support this hypothesis.

Associating expressed peptides to distinct CUs is not straightforward due to the high pairwise protein sequence identity (>95%). The cumulative abundance of proteins is highest at 20°C in St phase, whereas transcript expression increased with elevated temperature, for which the clustered expression profile follows the number of CUs in the transcript (Fig. 6c); this pattern was not observed in HSP70. In RuBisCo, multi-CU transcripts had a consistently low expression level, whereas single-CU derived transcripts were more strongly differentially expressed: *i.e*. upregulated under heat stress (Fig. 6c, Extended Data Fig. 8). Nine of the 16 complete RuBisCo gene models are in single exons, and no differential exon usage was identified in the multi-CU gene models. We found promoter motifs including a zinc finger motif in eight genes, but their presence does not correlate with the transcriptional profile. The modulation of expression responses in multi-*versus* single-CU transcripts reflects a regulatory mechanism in addition to alternative exon usage or RNA editing.

## Discussion

Our results provide an integrated multi-omics perspective on the molecular responses to heat stress in a pelagic HAB-forming dinoflagellate. These results are set against the backdrop of a large, complex genome structure and multi-level transcriptional regulation.

The high gene density, long introns^53, 54^, and extensive genetic duplication in *P. cordatum* likely reflect genomic hallmarks of bloom-forming dinoflagellates, consistent with data from *A. gibbosum*^22^. Habitats of *P. cordatum* are known to have rapidly fluctuating temperatures, such as diurnal vertical migration or in spring blooms and red tides. The elevated G+C content of the *P. cordatum* genome may be favored by selection to ensure high fidelity of transcription (*i.e*. G-C base pairs are more thermally stable). The long introns and presence of introner elements in the genome points to active transposition, *i.e*. non-autonomous DNA transposons, in contributing to the extensive rearrangement and duplication of genes^55–57^, and to the large genome size with many functionally redundant but slightly different transcript isoforms that provide rich adaptive resources in frequent changing environments in the aquatic realm.

Our data were generated from axenic cultures, thus the results reflect strict photoautotrophy, without the influence of cohabiting microbes, a host organism, and/or potential mixotrophy. *P. cordatum* is able to grow as a photoautotroph under heat stress^58^, supported by our observation of stable growth at 30°C. We report a dynamic transcriptional response to heat stress, with a more profound reprogramming during stationary growth phase compared to exponential phase. Our multiple lines of omics data demonstrate a strong coordinated response, particularly under chronic heat stress, that reduces energy production and consumption *via* the reduction of photosynthesis, carbon fixation, and amino acid biosynthesis, resulting in a suppressed central metabolism and protein synthesis. These observations lend support to the results of earlier transcriptome studies of pelagic dinoflagellates^59, 60^.

We present the first evidence of a complementary mechanism of post-transcriptional regulation in a pelagic dinoflagellate, involving RNA editing and exon usage for regulating molecular responses to elevated temperature. This mechanism likely provides the transcriptional flexibility that is essential for generating transcript and protein variants, and maximizes the functional diversity of gene products in dinoflagellates. In the absence of RNA editing and alternative exon usage, *e.g*. in single-exon genes that are common in dinoflagellates^37^, a third system we describe is through the adjustment of the number of coding units. All these mechanisms are set against the backdrop of polycistronic transcription followed by *trans*-splicing^30, 31, 61^.

HABs are occurring at an increasing frequency due in part to global climate change^5^. The multi-omics resources generated from this study provide a foundational reference for understanding the molecular regulatory and evolutionary processes of a HAB dinoflagellate species in response to environmental stress. The dynamics and interplay of molecular regulatory mechanisms may explain in part the complex and large genome in the context of vast diversification and evolutionary success of dinoflagellates over 696–1520 million years^62^.

## Methods

### Strain and culture conditions

The axenic culture of *Prorocentrum cordatum* strain CCMP1329 used in this work was obtained from the Provasoli-Guillard National Center for Marine Algae and Microbiota (NCMA). *P. cordatum* CCMP1329 was cultivated in L1 minus Si medium, which is a modified L1 medium^63^ in which synthetic ocean water was used instead of natural seawater, and Na_2_SiO_3_·9H_2_O omitted. Cultures were routinely maintained by transferring 15-day-old culture (10 mL) to fresh medium (90 mL) in 300-mL Erlenmeyer flasks. The flasks were kept in a climate chamber (RUMED type 3501; Rubarth Apparate GmbH, Laatzen, Germany) at 20°C in a 12:12 h light:dark cycle, with light intensity ~40 μmol photons m^−2^s^−1^ without agitation. Absence of contaminating bacteria was routinely checked by plating aliquots on LB and Difco^™^ marine agar 2216 (MB) plates. The number of cells mL^−1^ after transfer fluctuated between 2.0 × 10^4^ and 3.0 × 10^4^.

### Cell counting

Growth of *P. cordatum* was determined by cell counting using a BD LSR-Fortessa flow cytometer (BD Biosciences, San Jose, CA, USA). *P. cordatum* was identified according to its chlorophyll autofluorescence. Chlorophyll was excited with the 488-nm excitation laser and emission was detected at 695 nm. Samples (1 mL) were taken from three biological replicates during the light period and fixed with 25% glutaraldehyde (80 μL; final concentration 2% v/v) for 15 min at room temperature (RT). Samples were snap frozen in liquid nitrogen and stored at –70°C until they were analyzed. Each sample was analyzed in triplicates for 1 min.

Four independent growth curves per temperature were recorded. For each growth curve, counts were averaged from three biological replicates (each with three technical replicates). Counts were plotted against time and a generalized additive model (GAM) was fitted. The specific growth rate in the exponential growth phase (μexp per day) and the doubling rate per day (*k*) was calculated^64^.

### Extraction of genomic DNA

DNA of *P. cordatum* cells extracted with commercially available plant kits or by applying the common CTAB protocol was either too fragmented or contained too many contaminating compounds to be suitable for PacBio sequencing. Therefore, we performed an initial ultrasound treatment to break the cells and separate the nuclei from the debris, based on work aimed at isolating nuclei for electron microscopy^65^. This treatment was not performed for extracting genomic DNA for Illumina NovaSeq sequencing.

*P. cordatum* culture (100 mL) was transferred to two 50-mL Falcon tubes and centrifuged (685 *g*, 5 min, RT) using a Heraeus Multifuge^™^ X1R (Thermo Scientific, Schwerte, Germany). The supernatant was discarded, the pellets were dissolved in artificial seawater (15 mL) using an inoculation loop, and centrifuged (685 *g*, 5 min, RT). The supernatant was discarded, the pellets were dissolved in 30% ethanol (15 mL), and centrifuged (685 *g*, 5 min, RT); this step was repeated before the final pellet for each tube was dissolved in 30% ethanol (3 mL). For each tube, ultrasound using a Bandelin Probe Sonicator (Bandelin, Berlin, Germany) was applied (1 min, 5 cycles, amplitude 40%) followed by centrifugation (171 *g*, 3 min, RT), and the resulting pellet was suspended in 0.85% NaCl (1.5 mL) and centrifuged (10,000 *g*, 1 min, 4°C) using Heraeus Primo R Centrifuge (Thermo Scientific, Schwerte, Germany). Pellets from the two tubes were dissolved in high salt buffer (1 mL) in a 2-mL Eppendorf tube using inoculation loops. Proteinase K (8 μL) was added, and the mixture was incubated at 56°C (1 h), with the tubes inverted gently every 15 min. After cooling on ice (5 min), RNAse A (15 μL) was added, and the mixture was incubated at 37°C (30 min, no shaking). The sample was centrifuged (10,000 *g*, 5 min, 4°C) and the supernatant transferred to a new tube. NaCl (5M, 200 μL) were added, followed by thorough mixing by inverting the tube. Solution of CTAB/NaCl (200 μL) was then added, mixed well by inverting the tubes, and the mixture was incubated at 65°C (10 min). Chloroform extraction was performed using chloroform:isoamyl alcohol (24:1 v/v, 1 mL), and repeated three times until no interphase was visible. To the collected aqueous phases, an equal volume of isopropanol (pre-chilled at −20°C) was added, and the mixture was incubated overnight at −20°C. DNA was centrifuged (10,000 *g*, 10 min, 4°C) and the pellet was washed three times with 70% ethanol (pre-chilled at 4°C) with centrifugation (10,000 *g*, 10 min, 4°C). Following air drying under the clean bench, the DNA pellet was dissolved in the elution buffer (200 μL).

The DNA samples were sent to Helmholtz-Center for Infection Research (Braunschweig, Germany) for Illumina NovaSeq 6000 (pair-end 2 × 150bp) sequencing, and to the German Collection of Microorganisms and Cell Cultures (DSMZ, Braunschweig, Germany) PacBio Sequel and Sequel II sequencing.

### RNA isolation and sequencing

For RNA extraction, the sample was thawed at RT and transferred to a cryotube filled with 0.3 g acid-washed glass beads (100 μm). The cells were homogenized using the FastPrep-24 instrument (MP Biomedicals, Irvine CA, USA) at 6.0 m/s for 3 min (3 × 1 min, and 1 min on ice). Samples were centrifuged (12,000 *g*, 10 min, 4°C; Heraeus Primo R Centrifuge), and the supernatants were transferred to fresh tubes and incubated at RT (5 min). Next, 1-bromo-3-chlorophenol (100 μL; Sigma-Aldrich, Taufkirchen, Germany) was added, samples were shaken vigorously for 15 s and incubated at RT (10 min). Samples were centrifuged (12,000 *g*, 10 min, 4°C), the aqueous phase was transferred to a new tube, to which isopropanol (0.5 mL) was added, mixed, and incubated at RT (10 min). The sample was then centrifuged (12,000 *g*, 10 min, 4°C), after which the supernatant was removed. The RNA pellet was washed with 75% ethanol (1 mL) with centrifugation (7,500 *g*, 5 min, 4°C); this step was repeated. The final pellet was air-dried for 5 min, before being resuspended in RNase-free water (50 μL), and incubated at 55°C (10 min), prior to storage at −80°C.

Removal of genomic DNA was verified *via* PCR using total RNA as the template. The concentration of the RNA was quantified using a NanoDrop spectrophotometer (Peqlab, Erlangen, Germany) and the RNA integrity was assessed using a Bioanalyzer 2100 (Agilent, Santa Clara, USA). The average RNA concentration in the 18 samples was 297.4 + 109.5 ng μL^−1^ and the average RIN value was 5.5 + 0.78.

RNA sequencing was performed at the HZI Braunschweig with 300 cycles on a NovaSeq 6000 using pair-end 150bp chemistry with the library kit NEBNext Ultra II directional RNA. Ribosomal RNA was depleted prior to sequencing using polyA beads.

### *De novo* genome assembly

Genome data from *P. cordatum* were generated using Illumina NovaSeq 6000 and PacBio sequencing technologies, with a total data yield of 843.4 Gb (Supplementary Table 25). Combining these sequence reads, a hybrid genome assembly was generated using MaSuRCA v4.0.3^66^, independently with CABOG (option *FLYE_ASSEMBLY=0*) and FLYE (option *FLYE_ASSEMBLY=1*) as the assembly tool in the final step. Both assemblies are near identical, in which 99.94% of the scaffolds in each assembly share 99.04% identity on average. They exhibit the same level of data completeness (56.7%) based on recovery of BUSCO single-copy orthologs in alveolata_odb10 dataset (which is known to be poor in dinoflagellate data). Between these two preliminary assemblies, MaSuRCA-CABOG is more contiguous (N50 length of scaffolds = 194.50 Kb) and yields better recovery of the rRNA region (Supplementary Table 26); this assembly was used in the subsequent refinement steps.

To refine the assembled genome, we first incorporated RNA-Seq data to further scaffold the MaSuRCA-CABOG assembly using L_RNA_scaffolder^67^. Briefly, we mapped the *de novo* assembled transcriptome (for transcripts ≥ 500 bp) onto the assembled genome sequences using pblat v2.5^68^. The mapping results in psl format were used as input for L_RNA_scaffolder. This approach yielded a more-contiguous genome assembly (N50 length of scaffolds = 346.97 Kb) with a better recovery of BUSCO genes (57.9%; Supplementary Table 26).

Next, using BloBTools v1.1.1^69^, we assessed the assembled genome for potential outlier sequences based on sequence coverage, G+C content, and shared sequence similarity to known sequences in NCBI nt database (release 243; 15 April 2021). Genome scaffolds for which the sequencing coverage or G+C content is external to the range of median ± 1.5 × interquartile range are considered as potential outliers. Scaffolds that have bacterial, archaeal, or viral sequences as the top hits plus extreme sequencing coverage or extreme G+C content are considered sequences that are putatively external to the nuclear genome of *Prorocentrum cordatum*. In this analysis, majority (23,366; 98.2%) of the 24,295 genome scaffolds (implicating 3.89 Gb) do not have hits in the BLAST searches (20,914 scaffolds; 2.95Mbp) or have top hits in an undefined eukaryote sequence (2,452 scaffolds; 0.95Mbp); this observation is expected given the lack of dinoflagellate data in the existing databases. Using this approach, we identified 1,571 outlier sequences and removed them from the genome assembly. Most outlier sequences do not have shared similarity to bacterial sequences in the database. This is expected given the algal cultures from which the genomic DNA was extracted were axenic. The final genome assembly has a total size of 4.15 Gb (N50 length of scaffolds = 349.2 Kb; Supplementary Table 26).

### Transcriptome assembly

RNA-Seq reads from six conditions (20_ex, 20_st, 26_ex, 26_st, 30_ex and 30_st) were processed using fastp v0.21.0^70^ using parameter *–g* to remove adapter sequences and poly-G artifacts known in NovaSeq 6000 data. Transcriptomes were first assembled in “de novo” mode using Trinity v2.12.0^71^ independently for each condition. All *de novo* assembled transcripts were combined as a single assembly, from which redundant sequences were identified and removed using CD-HIT v4.8.1^72^ (98% identity; 0.9 length-difference cutoff), yielding the final representative reference transcriptome.

To generate the genome-guided transcriptome, processed RNA-Seq reads from each condition were first mapped onto the final genome assembly (above) using HISAT2 v2.2.1^73^. The mapping result (*i.e*. describing the alignments between RNA-Seq reads and genome scaffolds) was used as input for Trinity v2.12.0 in “genome-guided” mode. Using the same strategy above, individual genome-guided assemblies were combined, and redundant sequences removed, yielding the final representative genome-guided transcriptome.

### *Ab initio* prediction of protein-coding genes

To predict protein-coding genes, we follow the customized gene prediction workflow tailored for dinoflagellate genomes following Chen et al.^74^. The description of this workflow is available at https://github.com/TimothyStephens/Dinoflagellate_Annotation_Workflow. Briefly, this approach integrates evidence of transcript and protein sequences to guide predictions using multiple gene programs, after which the results were integrated to yield the final gene models.

First, we identified novel repetitive elements in the genome assembly using RepeatModeler v2.0.1 (http://www.repeatmasker.org/RepeatModeler/), combined these elements with existing repeat families in RepeatMasker database release 2018/10/26 as a reference, to predict and mask all repetitive sequence regions from the genome sequences using RepeatMasker v4.1.0 (http://www.repeatmasker.org/); this yields the repeat-masked genome assembly.

To predict protein-coding genes, we first mapped the representative *de novo* and genome-guided transcriptome assemblies to the genome assembly using Minimap2 v2.18^75^, for which the code was modified to recognize G-C and G-A donor splice sites. The mapping information was then used to predict transcript-based genes using PASA v2.3.3^76^ for which the code was modified to account for non-canonical splice sites. The proteins coded by the transcript-based genes were searched (BLASTP, *E* ≤ 10^−20^, >80% query cover) against a customized protein database combining RefSeq (release 98) proteins and predicted proteins from available Symbiodiniaceae genomes (Supplementary Table 27). The gene models were checked for transposable elements using HHblits v3.3.0^77^ and TransposonPSI (https://github.com/NBISweden/TransposonPSI), searching against the JAMg transposon database (https://github.com/genomecuration/JAMg); those genes containing transposable elements were removed from subsequent steps. Redundant sequences were removed based on similarity using CD-HIT v4.8.1^72^ (*-c 0.75 -n 5*). Among the remaining transcript-based gene sequences, we identified high-quality “golden genes” using the script *Prepare_golden_genes_for_predictors.pl* from the JAMg pipeline (https://github.com/genomecuration/JAMg), altered to recognize alternative splice sites. These “golden genes” represent high-quality training set for *ab initio* gene predictors. We used them as the training set for SNAP^78^ and AUGUSTUS v3.3.1^79^ for gene prediction from the repeat-masked genome assembly; the codes for AUGUSTUS was also modified to recognize alternative splice sites (https://github.com/chancx/dinoflag-alt-splice).

The repeat-masked genome was also used as the input for GeneMark-ES^80^. We also predicted genes using MAKER v2.31.10^81^, in which the code was modified to recognize GA donor splice sites. Protein-coding genes were predicted using MAKER (*protein2genome* mode) based on protein sequences from the Swiss-Prot database (retrieved 02 March 2020) and predicted protein sequences from other Symbiodiniaceae genomes. Finally, gene predictions from the five methods including the *ab initio* predictors (GeneMark-ES, AUGUSTUS, SNAP), MAKER protein-based predictions, and PASA transcript-based predictions were integrated using EvidenceModeler v1.1.1^82^ to yield the high-confident gene models; the weighting is PASA 10, MAKER protein 8, AUGUSTUS 6, SNAP 2 and GeneMark-ES 2. These gene models were subjected to three rounds of polishing during which the gene models were corrected based on transcriptome re-mapping using the annotation update utility in PASA^76^.

Introner elements were identified in the intronic regions using the program Pattern locator^83^. The patterns we searched for were inverted repeats of 8-20 nucleotides and direct repeats of 3-5 nucleotides within 30 bases of each end of the introns as described in Farhat et al.^84^.

### Functional annotation of protein-coding genes

Annotation of gene function for *P. cordatum* and other representative dinoflagellate genomes was conducted based on sequence similarity to known proteins in the UniProt database (release 2021_03). Predicted protein sequences from the gene models were first searched against the manually curated protein sequences of Swiss-Prot (UniProt release 2021_03) using BLASTp v2.3.0+ (*E* ≤ 10^−5^; subject-sequence cover ≥ 70%). Sequences that have no Swiss-Prot hits were then searched against TrEMBL (UniProt release 2021_03) using the same parameters. For predicted proteins of *P. cordatum*, we further assessed functions based on sequence-similarity search against known protein sequences in EnzymeDetector, InterProScan, eggNOG and Kofam.

### Prediction of transit peptides

For each predicted protein, transit peptides were first predicted independently using TargetP v2.0 (*-org pl*), SignalP v6 (*--organism eukarya --mode fast*), WoLF PSORT (*plant* mode), Predotar, and ChloroP v1.1. Subcellular localization is determined based on the consensus from these predictions confirmed in three or more programs.

### Analysis of homologous proteins

To identify homologous proteins of the predicted *P. cordatum* proteins, we compiled a comprehensive protein-sequence database (1,554,705 sequences from 31 dinoflagellate taxa; Supplementary Table 28) using available genome or transcriptome data. All data from the MMETSP database^85^ were downloaded from https://zenodo.org/record/1212585#.YlYczuhByUk. For species where there were multiple datasets for the same isolate, the protein sequences were clustered at 90% sequence identity using CD-HIT v4.8.1^72^ to reduce redundancy. Using these 31 sets of protein sequences, homologous sets were then identified based on clustering of protein sequences based on sequence identity using OrthoFinder v2.3.8^86^ at default settings.

### Identification of mRNA editing sites

Putative mRNA editing events were identified from single nucleotide variations observed in genome-sequence reads *versus* RNA-Seq reads. An observed nucleotide variation in the RNA-Seq reads but not in genome-sequence reads is considered a potential mRNA edited site. Briefly, 25% of all genome-sequence reads (randomly sampled) were mapped to the final genome assembly using bwa-mem v0.7.17-r1188 (https://github.com/lh3/bwa). Trimmed RNA reads from each sample (6 conditions × 3 replicates) were mapped to the genome assembly separately with HISAT2 v2.2.1^73^ using default parameters (--nodiscordant) and a HGFM index that was built using known exons and splice sites from the predicted gene models. PCR duplicates were marked by *MarkDuplicates* implemented in Picard v2.23.8 (https://broadinstitute.github.io/picard/). For each condition, mapping of RNA-Seq reads was compared with the mapped genome-sequence reads using JACUSA v2 (*call-2 -F 1024 -P2 RF-FIRSTSTRAND -s -a D,Y,H:condition=1*). We follow the authors’ recommendation to assess the statistical significance of an mRNA edited site. A site is considered statistically significant if it meets all the requirements: (a) a score >1.15; (b) coverage of genome reads >10; (c) coverage of RNA reads from each condition >5; (d) number of putative editing type is <2; (e) The editing site is present in all three replicates.

### Analysis of horizontal gene transfer

To identify putative horizontal gene transfer (HGT), *P. cordatum* proteins were searched (BLASTP, *E* ≤ 10^−5^) against a customized protein-sequence database that consist of 2,773,521 proteins from 82 other eukaryotes (Supplementary Table 29) and 688,212 proteins from 543 single-cell assembled genomes (SAGs) of prokaryotes^87^ (Supplementary Table 30). Excluding hits to other *Prorocentrum* proteins, *P. cordatum* proteins that have a bacterial top hit are considered results of HGT involving *P. cordatum* and bacteria. To support the inferences of putative HGT, we employed OrthoFinder v2.5.4^86^ to infer homologous protein sets from all the involved proteins (*i.e. P. cordatum* proteins, proteins from other eukaryotes and SAGs). Homologous protein-sequence sets that contain *P. cordatum* proteins implicated in HGT were multiply aligned using MAFFT v7.453^88^ at *--maxiterate 1000*. Trimmed with trimAl v1.4.1^89^ using parameter *-automated1*, the alignments were used to infer phylogenetic trees using IQ-TREE2^90^ at *-B 2000 -T AUTO*.

### Integrated mixOmics analysis

We conducted a multi-omics analysis in *P. cordatum* using the proteomic and transcriptomic data to identify a systems level heat stress response across the growth phases using the *mixOmics* package^91^ in R. Transcriptome FPKM values were first log_2_ transformed prior to quality filtering with normalized proteomic data, requiring features be present across 75% of samples. Selected features represented the top 25% and 50% most variable transcripts (15,097) and proteins (259), respectively, that were present across 75% of the samples. These features were then input to mixOmics for Data Integration Analysis for Biomarker discovery using a Latent cOmponents (DIABLO)^42^.

We conducted performance testing of the initial model to identify the number of latent components that contained a multi-omics signature using Mfold validation with 5 folds and 50 repeats. This suggested two latent components as the best fits for the model. We then performed final model tuning using the max distance to select diagnostic features for both component 1 (RNA-Seq: 170, proteins: 30) and component 2 (RNA-Seq: 190, proteins: 40). Ordination of all features indicate the separation of the three temperature levels for both transcriptome and proteome features. The transcriptome and proteome features selected for component 1 discriminate a common heat stress response at both 26°C and 30°C (Extended Data Fig. 2, Supplementary Table 12) and for component 2, a heat stress response specific to either 26°C or 30°C (Extended Data Fig. 3, Supplementary Table 12). The variates for the pathways from both components were then input to NetworkAnalyst^92^ for KEGG pathway over-representation analysis to identify functional categories that were enriched in each network.

A relevance associations network was created for each component using the *network* function within *mixOmics*, where values represent a robust approximation of the Pearson correlation. A heatmap displaying the features from each component was created using the *pheatmap* package in R with features clustered according to their Euclidean distances and scaled within rows. This revealed two main clusters within each component that were extracted using the R package *dendextend* according to the corresponding dendogram. A relevance associations network was then created for each subcluster as previously done for the two components.

### Experimental design of multi-omics analysis

Cultures of *P. cordatum* CCMP1329 were maintained as described above. After 15 days of cultivation at 20°C, 10 mL were transferred to 90 mL of fresh L1-Si medium in 300-mL Erlenmeyer flasks and placed in a climate chamber set to the desired temperature (26°C or 30°C) and exhibiting the same light intensity and light:dark cycle. These temperatures are already common in the Red Sea^93^ or near the equator^94^, and are well within the range predicted for the future oceans^95^. Replicate cultures were set up under identical conditions to allow sampling of 3 biological replicates each for transcriptome and proteome and 5 biological replicates for metabolome. For proteome analysis, up to 12 L of culture (120 flasks each with 100 mL culture) were cultivated in the same climate chamber to obtain ~2 g biomass (wet weight). For each growth stage (exponential, stationary) and temperature (20°C, 26°C, 30°C) a complete set of cultures were sacrificed. Cell counting and harvesting were started about 5 h after the onset of the light period. For cell counting, random samples were chosen from the climate chamber to account for slight differences in light intensity.

For transcriptome analysis, three 100 mL cultures (biological replicates) were sampled per temperature and growth phase. Each culture was centrifuged in two 50-mL Falcon tubes at (4,276 *g*, 4°C, 5 min) in a Heraeus Multifuge^™^ X1R. The supernatant was decanted, and the pellet was resuspended in the remaining medium. The two pellets were combined in an Eppendorf tube (2 mL) and centrifuged (17,000 *g*, 4°C, 3 min) in a Heraeus Primo R centrifuge. The remaining supernatant was removed by pipetting, and the weight of the wet pellet was determined. The pellet was resuspended in 1 mL TRIzol reagent (Thermo Fisher Scientific, Waltham MA, USA), snap frozen in liquid nitrogen and stored at −70°C until further analysis.

For proteome analysis, tubes and buffers used in the following steps were pre-chilled; all steps were conducted on ice. The culture (400 mL) was filled into pre-chilled 500-mL centrifuge bottles, centrifuged (4,248 *g*, 4°C, 30 min) using a Sorval Lynx 4000 (Thermo Fischer). The supernatant was decanted, the pellet was resuspended in a buffer (100 mL) containing Tris-HCl (100 mM, pH 7.5) and MgCl_2_· 6H_2_O (5 mM). The re-suspended pellets were centrifuged (4,248 *g*, 4°C, 30 min), and the supernatant was removed. The pellet was resuspended by gently pipetting in the same buffer (800 μL). The suspension was transferred into 2-mL Eppendorf tubes and centrifuged (17,000 *g*, 4°C, 5 min) using a Heraeus Primo R centrifuge. The supernatant was removed by pipetting and the weight of the wet pellet was determined. Samples were frozen in liquid nitrogen and stored at −70°C.

For metabolome analysis, 15 mL from five cultures (100 mL each; as biological replicates) from each temperature and growth phase were extracted immediately after cell counting with a filtration unit and 0.22 μm Millipore membrane filter. Samples were filtered at 500 mbar with a vacuum controller. The cells were washed three times with 4°C cold 3.5% NaCl.

The filters were immediately transferred to 5 mL Eppendorf tubes containing 100 mg of glass beads (0.7–100 μm; Kuhmichel, Ratingen, Germany), three stainless steel beads (two 5 mm^3^ and one 10 mm^3^; Kugel Winnie, Bamberg, Germany) for partially destroying the filter and to obtain cell lysis. Cold extraction fluid (2 mL, per-chilled at −20°C) was immediately added; this extraction fluid for metabolite extraction contained methanol, ethanol and chloroform^96^, and the internal standard ^13^C-ribitol. The pre-chilled 5-mL tubes with the filter and extraction fluid on ice were mixed for 20 s and placed back on ice until further treatment.

### Analysis of transcriptome and differentially expressed genes

The transcriptome reads were mapped to the assembled reference genome using HISAT2^73^. The reads mapped onto the exons were counted for the corresponding genes with featureCounts^97^. For differential expression analysis, only uniquely mapped reads were used to avoid ambiguity. The fragments per kilobase of transcript per million mapped fragments (FPKM) value of each gene was calculated by normalizing the fragments per million with the sum of the exon length of the corresponding gene. The ternary visualization of the gene expression pattern across three temperatures was produced with R package *ggtern*^98^.

For analysis of differentially expressed genes (DEGs), genes with low expression were filtered out by the *filterbyexpr* function in edgeR^99^ using default parameters. Then, the DEG analysis was performed with edgeR using the recommended *glmQLF* test on the raw read count per gene. Genes with a Benjamini-Hochberg corrected *p*-value ≤ 0.001 and an absolute log_2_-fold-change (log2FC) ≥ 2 were considered as significantly differentially expressed.

Hierarchical clustering with the complete-linkage algorithm was used to identify gene clusters based on their expression profile across temperature conditions. The input expression values were centered log_2_-transformed FPKM values, *i.e*. log_2_(FPKM + 1) centered by subtracting the mean. The tree was cut into eight clusters to represent different expression profiles. Superclusters 4 and 7 in the stationary phase (each with small number of genes) showed very similar expression pattern; these were merged for downstream functional enrichment analysis.

### Functional analysis of DEGs

The GO term enrichment analysis was carried out with the R package topGO (https://doi.org/10.18129/B9.bioc.topGO) for all three gene ontologies, *i.e*., biological process (BP), cellular component (CC) and molecular function (MF). REVIGO^100^ was used to summarize the GO terms according to the semantic similarity for a concise visualization (Supplementary Fig. 8). In order to perform a KEGG pathway enrichment analysis, we assigned KEGG ortholog (KO) number for each gene using KOfam^101^. When a gene had multiple KO assignments with an e-value ≤ 10^−10^, we chose the one with the lowest e-value. If a gene had several KO assignments with an e-value of zero, we kept all those KO assignments for this gene. Using this annotation, the genes with KO assigned held about 50% of mapped reads. The expression profile for each KO gene was then generated by summing up the read count for all genes belonging to the same KO gene. Differentially expressed KO genes were then identified using edgeR similar to the above DEG analysis. The KEGG pathway and module enrichment analyses were performed using clusterProfiler^102^ on the DEGs. For both GO term enrichment analysis and KEGG enrichment analysis, a false-discovery rate (FDR) ≤ 0.05 was considered as significantly enriched.

In the central carbon metabolism pathway analysis, a paired Wilcoxon test was performed to compare the expression change of the gene members belonging to the same EC number. The alteration of a metabolite’s concentration was analyzed using Wilcoxon test on the mean concentration values of biological replicates over the three technical replicates; FDR ≤ 0.05 was regarded as significantly changed.

### Proteome analysis

Cells of *P. cordatum* were resuspended in solubilization buffer and disrupted by bead beating (FastPrep-24 5G, MP Biomedicals) for 10 s at 6 m/s followed by 90 s on ice (in three repetitions) using 0.1 mm silica spheres. Cell debris and insoluble material were removed by ultracentrifugation (104,000 *g*, 1 h, 17°C) and the supernatant was again centrifuged (200,000 *g*, 1 h, 17°C). The protein content of the preparation was determined according to Bradford^103^; 50 μg was subjected to reduction, alkylation and tryptic digestion as reported previously^104^. Generated peptides were analyzed by (a) one-dimensional (1D) reversed phase nanoLC and (b) two dimensional (2D) SCX-/RP nanoLC separation coupled to MS-detection. For 1D separation, tryptic peptides (1 mg) were decomplexed by reversed phase nanoLC separation (Ultimate 3000 nanoRSLC; Thermo Fisher Scientific, Dreieich, Germany) using a trap column setup (2 cm length, 5 μm bead size, 75 μm inner diameter; Thermo Fisher Scientific) with a 25 cm separation column (2 μm bead size, 75 μm inner diameter; Thermo Fisher Scientific), and applying a 360 min linear gradient^104^. In case of 2D separation, 4 μg were subjected to SCX fractionation (eluent A: 5 mM H_3_PO_4_, 5% v/v acetonitrile, pH 3.0; eluent B: 5 mM H_3_PO_4_, 5% v/v acetonitrile, 1 M NaCl, pH 3.0) with a linear 20 min gradient and collection of 19 fractions per sample beginning 5 min after injection. Each fraction was subsequently applied for second dimension reversed phase separation (see above) using a 60 min gradient. For both methods, the eluent was continuously ionized (captive spray ion source; Bruker Daltonik GmbH, Bremen, Germany) and ions analyzed by an ion-trap mass spectrometer (amaZon speed ETD; Bruker Daltonik GmbH) as described in Wöhlbrand et al.^105^.

To prepare the membrane protein-enriched fraction, cell pellets were gently thawed, resuspended in 0.5 mL membrane lysis buffer (MLB; 100 mM Tris-HCl, 2 mM MgCl_2_, 10 % (w/v) glycerin, 0.5 mM DTT, pH 7.5), and disrupted by bead beating (as described above). DNA was digested using DNase I and the obtained raw fraction applied on top of a continuous sucrose gradient (30–80% w/v) prior to centrifugation (37,800 *g*, 12 h, 4°C).

Membrane containing fractions were collected and washed twice with MLB (centrifugation at 104,000 *g*, 1 h, 4°C). Obtained membrane pellets were resuspended in MLB (500 μL), pooled and centrifugation repeated. The final pellet was resuspended in sodium dodecyl sulfate (SDS, 300 μL, 1.0 % w/v) and incubated at 95°C (5 min) prior to centrifugation (20,000 *g*, 20 min, 20°C). The supernatant was snap-frozen in liquid nitrogen until further analysis. Protein content was determined using the RC-DC^™^ Protein Assay (BioRad GmbH, Munich, Germany). A total of 10 μg protein per fraction of each sample was separated by SDS-polyacrylamide gel electrophoresis (SDS-PAGE), and gels were stained with Coomassie brilliant blue^106^. Each sample lane was cut into 8 slices, and each slice into small pieces of ~1-2 mm^2^ for subsequent in-gel digest as previously described^105^. The generated peptide solutions were analyzed by reversed phase nanoLC-MS (as described above), *via* a 120 min gradient. Per sample, protein search results of each slice of fractions I and II were compiled.

Protein identification was performed using Mascot (version 2.3; Matrix Science, London, UK) *via* the ProteinScape platform (version 4.2; Bruker Daltonik GmbH) against the genomic database of *P. cordatum*. A target-decoy strategy with a false discovery rate <1.0% was applied together with following settings: enzyme = trypsin, missed cleavage allowed = 1, carbamidomethylation (*C*) = fixed, oxidation (*M*) = variable modification, peptide and MS/MS mass tolerance = 0.3 Da, monoisotopic mode, peptide charge = 2+ and 3+, instrument type = ESI-TRAP, significance threshold *p* < 0.05, ion score cutoff = 25.0, and minimum peptide length = 5. Search results of individual searches per sample were compiled using the ProteinExtractor function of ProteinScape.

### Metabolome analysis

The cell disruption was performed with MM400 oscillating mill (Retsch, Haan, Germany) at 30 Hz for 2 min, and repeated three times. The cooling during the treatment was ensured by using –80°C pre-chilled sample containers. For metabolite extraction, the tubes were centrifuged (14,489 *g*, 8 min, 4°C). Two volumes (100 μL and 200 μL) of the supernatant were transferred to gas chromatography (GC) glass *via*ls (Klaus Trott Chromatographiezubehör, Kriftel, Germany) and dried under vacuum (Labconco, Kansas City, Missouri, USA) at 4°C. After drying overnight, the *via*ls were capped with magnetic *via*l caps (Klaus Ziemer GmbH, Langerwehe, Germany) and stored at −80°C until measurements. All samples belonging to one experiment were analyzed in a batch.

To analyze polar metabolites with gas chromatography and mass spectrometry (GC-MS), a derivatization was performed to reduce polar interactions by introducing non-polar trimethylsilyl groups. This was necessary to evaporate the substances under low temperatures and avoid alteration. After extraction, the samples were analyzed with a gas chromatograph connected to a mass spectrometer (Agilent 7890A and 5975C inert XL Mass Selective Detector). The sample derivatization was automatically done with a multisampler (GERSTEL, Mühlheim an der Ruhr, Germany) and 2% methoxyamine hydrochloride (MeOX; 15 μL) in pyridine (Roth AG, Arlesheim, Switzerland) for 60 min at 40°C followed by the addition of 15 μL *N*-tert-butyldimethylsilyl-*N*-methyl and incubated at 40°C (30 min). For measurement, the sample (1 μL) was injected in splitless-mode into a split/splitless (SSL) injector at 270°C. The GC was equipped with a 30 m DB-35MS + 5 m DuraGuard capillary column (0.25 mm inner diameter, 0.25 μm film thickness; Agilent Technologies). Helium was used as carrier gas at a flow rate of 1.0 mL min^−1^ and the GC-oven was run at following temperatures and times per sample: 6 min at 80°C; 6 min from 80°C to 300°C; 10 min at 300°C; 2.5 min from 300°C to 325°C; and 4 min at 325°C. Each GC-MS run lasted 60 minutes. The transmission temperature from GC to MS was 280°C and the temperatures of the MS and the quadrupole were 230°C and 150°C, respectively. The ionization in the mass detector was performed at 70 eV. The detector was operated in SCAN-mode for full scan spectra from 70 *m/z* to 800 *m/z* with 2 scans s^−1^. For calibration of retention indices, the alkane standard mixture (C_10_-C_40_; Sigma Aldrich) was used.

Data analysis was performed using MetaboliteDetector^107^. After calibration with the alkane mixture, a normalization by ^13^C-ribitol was performed to eliminate variations in sample volumes. Batch quantifications were performed with MetaboliteDetector. Non-targeted analysis was performed with an in-house library and the following settings: peak threshold 5, minimum peak height 5, 10 bins per scan, deconvolution with 5, no baseline adjustment, compound reproducibility 1.00, maximum peak of discrimination index 100.00, and minimum number of ions = 20.

## Acknowledgements

This project was funded by the German Research Organisation (DFG) through the Transregio SFB TRR-52 Roseobacter. The following co-authors were funded by this agency: Z.L.D., L.W., C.R., B.B., J.H., K.H., J.K., J.M., J.O., J.P., S.S.G., K.S.H., C.S., H.S., H.W., R.R., D.J. and I.W.D. This project was also supported by Australian Research Council Discovery Project DP190102474 awarded to C.X.C. and D.B., the Australian National Computational Infrastructure (NCI) National Facility systems through the NCI Merit Allocation Scheme (Project d85) awarded to C.X.C., and other high-performance computing facilities at the Australian Centre for Ecogenomics and Research Computing Centre at the University of Queensland. We would like to thank Nicole Heyer and Simone Schrader for technical assistance in genome sequencing, and Sinem Bektas for her assistance in the wetlab and her enormous flexibility, unfailing reliability, and good spirits even in the worst of Corona times.

## Author contributions

C.X.C. and I.W.D. conceived the study; K.E.D., Z.L.D., L.W., C.R., B.B., R.R., C.X.C. and I.W.D. designed the analyses; K.E.D. led and performed the analyses of integrated multi-omics data and differential exon usage; Z.L.D. led and performed the analysis of transcriptome and differential gene expression; L.W. led and performed the analysis of proteome data; C.R. led and performed the analysis of metabolome data; K.E.D., Z.L.D., L.W., C.R., Y.C., J.H., U.J., S.S., and C.X.C. conducted all computational analyses; B.B. and C.S. performed the PacBio sequencing; Y.C. performed structural and functional annotation of the genome, and conducted analysis of genome sequence features and RNA editing; J.H. and U.W. conducted the preliminary genome analyses; K.H. and K.S.H. analyzed the metabolome data; J.K. conducted measurement and analysis of proteome samples; J.M. and S.S.G. maintained the cell cultures, determined growth rates, and extracted biological samples for multi-omics analysis; M.N.S. led the analysis extracted DNA samples; J.O. provided infrastructure necessary for this research; J.P. and H.W. optimized the extraction of RNA; S.S. conducted analysis of non-coding genomic elements; H.S. optimized the extraction genomic DNA; D.B., R.R. and D.J. contributed intellectual input to the structure and presentation of the manuscript; C.X.C. performed genome assembly and the comparative genome analysis; I.W.D. oversaw and designed the multi-omics experiment; K.E.D., Z.L.D., L.W., C.R., C.X.C, Y.C. and S.S. prepared all figures and tables; K.E.D., Z.L.D., L.W., C.R., R.R., C.X.C. and I.W.D. prepared corresponding draft sections for the manuscript; C.X.C. and I.W.D. prepared the first draft of the manuscript based on input from all authors; K.E.D., Z.L.D., L.W., U.J., D.B., R.R., D.J., C.X.C., and I.W.D. wrote, reviewed and commented on the manuscript; all authors approved the final manuscript.

## Data availability statement

The assembled genome and transcriptome, and the predicted gene and protein sequences from *P. cordatum* are available at https://cloudstor.aarnet.edu.au/plus/s/nxC7atJTpJuyr8h. The BioProject accession for the genome and transcriptome data is PRJEB54915.

Detailed information on identified proteins and peptides is deposited at fairdomhub.org under the project 293 (DOI will be available once the data are publicly available). For review, the data are available *via* the following links:

Metabolites (standard operating procedure):

https://fairdomhub.org/sops/551?code=RrSjntdGsUnt2zRDdFKIKnfmJviLFvw3O7BzG9Lv

Proteome (standard operating procedure):

https://fairdomhub.org/sops/549?code=TKCZcudIA05oY%2FSnAFW67R2IwE91d1x%2F9eQp6dnH

Metabolites (data):

https://fairdomhub.org/data_files/6075?code=kyxQ9ljLVTxjU54Wrk5HS62bQj6Z9niiJtT30GSc

Shotgun proteomics (data):

https://fairdomhub.org/data_files/6059?code=GhhJoFC9oBKYd7fWRlE9kZ%2BpMGnGRMD2Cvtf8sLE

Membrane fraction of proteins (data):

https://fairdomhub.org/data_files/6058?code=QwaeOVVnEvuCXftZqObFSFD82emv0qEqhs9oiHV1

2D LC-MS/MS of proteins (data):

https://fairdomhub.org/data_files/6057?code=4PPrRrP6qwTuD1iekFWzxzRijAyGkbeaySaybJjI

## Extended Data

**Extended Data Table 1.** Statistics of assembled genomes of *P. cordatum* and other dinoflagellates.

**Extended Data Table 2.** Statistics of predicted genes in *P. cordatum* and other dinoflagellates.

**Extended Data Fig. 1.** Heatmap of all identified metabolites (*p* < 0.001) at 20°C, 26°C and 30°C in *P. cordatum*, for Ex and St phase. Five biological replicates and three technical replicates were run per experiment. Relative intensity values are shown, and data were normalized to internal standard and cell count. Normalization for heatmap was done by using z-score, and significance was calculated using two-tailed T-test.

**Extended Data Fig. 2.** Heatmap of expression levels for the inferred multi-omics signature that is indicative of a general heat stress response (component 1), shown for values scaled across rows with proteins and transcripts that are (a) up-regulated and (b) down-regulated during heat stress.

**Extended Data Fig. 3.** Heatmap of expression levels for the inferred multi-omics signature that is indicative of a heat-stress response specific to 26°C and to 30°C, shown for values scaled across rows with proteins and transcripts that are (a) up-regulated and (b) down-regulated during heat stress.

**Extended Data Fig. 4.** Detailed reconstruction of the central carbon metabolism of *P. cordatum* and the concerted responses to heat stress response at transcript (blue), protein (red) and metabolite (yellow) levels.

**Extended Data Fig. 5.** Clusters of genes with similar expression patterns across different conditions in (a) exponential (Ex) and (b) stationary (St) phase, and (c) enriched gene functions in the Molecular Function GO terms in SC2, SC3, SC5, and SC6 in the St phase.

**Extended Data Fig. 6.** Enriched functions in differentially regulated genes in 26°C *versus* 20°C in stationary phase, showing enriched Molecular Function GO terms in (a) up-regulated and (b) down-regulated genes, and the enriched KEGG pathways in (c) up-regulated and (d) down-regulated genes.

**Extended Data Fig. 7.** Sashimi plot across the HSP70 locus displaying read coverage with lines between exons indicating connections supported by >500 alignments.

**Extended Data Fig. 8.** Transcript profiles of (a) HSP70 and (b) RuBisCo gene models based on clustering in Figures 6a and 6c.

## Supplementary Information

### Supplementary Note

#### Supplementary Figures

**Supplementary Fig. 1.** Analysis of P. cordatum genome based on (a) distribution of 21-mers indicating a haploid genome, (b) genome-size estimation based on 21-mers using GenomeScope2, and (c) repeat landscape excluding unknown repeats.

**Supplementary Fig. 2.** Gene functions encoded in the genome of *P. cordatum* and the other five representative taxa based on count of Gene Ontology (GO) terms.

**Supplementary Fig. 3.** Unidirectional coding of genes in dinoflagellate genomes based on gene-orientation changes in ten-gene windows.

**Supplementary Fig. 4.** Maximum likelihood protein tree of a putative hydantoin racemase indicating bacterial origin of the encoding gene in *P. cordatum*.

**Supplementary Fig. 5.** Transcriptome analysis of *P. cordatum* showing (a) principal component analysis of samples based on FPKM values, and (b) Venn diagram detailing number of DEGs between any two temperature conditions, independently for Ex and St phases.

**Supplementary Fig. 6.** The proteome landscape of *P. cordatum* heat-stress response, showing (a) NMDS visualization of individual samples based on peptide count data (max error = 1.24 × 10^−11^), and (b) Venn diagram of number of proteins in distinct conditions.

**Supplementary Fig. 7.** Network visualization of multi-omics signature inferred from both proteins and transcripts that represent a heat-stress response specific to 26°C and to 30°C. Colors of the lines indicate the nature of the relationships between proteins and transcripts, with positive relationships in green and negative relationships in purple.

**Supplementary Fig. 8.** Functional enrichment of dispersed gene copies based on GO terms in homologous sets of size 20 or greater (total of 13,529 genes), shown for (a) Biological Process, (b) Molecular Function, and (c) Cellular Component.

**Supplementary Fig. 9.** Schematic diagram three distinct gene models identified in *P. cordatum*: (a) standard gene model, (b) gene model with polycistronic mRNA, and (c) gene model with multiple coding units.

**Supplementary Fig. 10.** Detailed information on gene models encoding multiple coding units (CUs), showing the scale model for (a) those containing only fragments of RuBisCo CU with s10_g491 as a reference that encodes 3 CUs; (b) RuBisCo gene models and the corresponding exons; and (c) those containing multiple CUs that encode protein functions related to central metabolism.

**Supplementary Fig. 11.** Comparison of RuBisCo gene models of P. cordatum and other dinoflagellate species, showing the (a) alignment of the predicted leader sequences, N-terminal sequence of the first CU, and the spacer region between two complete CUs; (b) alignment of spacer regions of chloroplast-localized proteins (RuBisCo and phosphoglycerate kinase) and non-chloroplastic proteins (HSPv70 and glyceraldehyde 3-phosphate dehydrogenase); and (c) histogram of Rubisco genes encoded in the genomes of P. cordatum and S. microadriaticum relative to protein-sequence length.

#### Supplementary Tables

**Supplementary Table 1.** *P. cordatum* genes that contain introner elements in one or more introns.

**Supplementary Table 2.** Introner elements in genomes of dinoflagellates.

**Supplementary Table 3.** Functional annotation of 85,849 *P. cordatum* proteins.

**Supplementary Table 4.** Annotated protein sequences from genome of *P. cordatum* and other dinoflagellates.

**Supplementary Table 5.** Classification of gene duplicates in *P. cordatum* and the five representative dinoflagellate taxa.

**Supplementary Table 6.** Tandemly duplicated gene blocks in *P. cordatum* genome.

**Supplementary Table 7.** The 47 *P. cordatum* proteins with top bacterial hit in GORG-Tropics.

**Supplementary Table 8.** Summary statistics of RNA-Seq transcriptome data generated and reads mapped to the genome sequences.

**Supplementary Table 9.** Expression of genes in all conditions shown in FPKM values.

**Supplementary Table 10.** Normalized peptide count per gene model of *P. cordatum*.

**Supplementary Table 11.** Relative intensity values of metabolite data measured by GC-MS.

**Supplementary Table 12.** *P. cordatum* genes and their corresponding UniProt-KB annotations for the transcriptome and proteome datasets selected by mixOmics DIABLO as indicative of the two components.

**Supplementary Table 13.** Transcript (FPKM) and protein (normalized peptide counts) expression for gene functions related to photosynthesis, oxidative phosphorylation, and central carbon metabolism.

**Supplementary Table 14.** Wilcoxon test for expression level for transcripts and proteins related to enzymes of central carbon metabolism in the exponential growth phase.

**Supplementary Table 15.** Differentially expressed genes in all possible pairwise comparisons of growth conditions.

**Supplementary Table 16.** Clusters of differentially expressed genes based on expression profile during exponential growth phase.

**Supplementary Table 17.** Clusters of differentially expressed genes based on expression profile during stationary growth phase.

**Supplementary Table 18.** Enrichment analysis of GO terms in differentially expressed genes.

**Supplementary Table 19.** Analysis of differentially expressed genes based on annotated KEGG Orthology (KO) terms.

**Supplementary Table 20.** Enrichment analysis of KEGG pathways based on differentially expressed KO genes.

**Supplementary Table 21.** Expression of 13 putative polyketide synthase I genes in *P. cordatum*.

**Supplementary Table 22.** Differential exon usage identified in *P. cordatum* genes.

**Supplementary Table 23.** Differentially edited mRNA sites identified in *P. cordatum* genes.

**Supplementary Table 24.** The 17,214 gene models associated with DinoSL-containing transcripts in *P. cordatum*.

**Supplementary Table 25.** Genome sequence data generated for this study.

**Supplementary Table 26.** Statistics of preliminary and final genome assemblies of *P. cordatum*.

**Supplementary Table 27.** Dinoflagellate proteins used to guide prediction of protein-coding genes from the *P. cordatum* genome.

**Supplementary Table 28.** Protein sequences from 31 dinoflagellate taxa used for identification of homologous sequence sets.

**Supplementary Table 29.** The protein sequences from 82 eukaryote taxa used in this study.

**Supplementary Table 30.** The 688,212 protein sequences from 543 single-cell assembled genomes in GORG-Tropics used in this study.

## References

1. Brown, A.R. et al. Assessing risks and mitigating impacts of harmful algal blooms on mariculture and marine fisheries. Reviews in Aquaculture 12, 1663–1688 (2020).

2. Wells, M.L. et al. Future HAB science: directions and challenges in a changing climate. Harmful Algae 91, 101632 (2020).

3. Karlson, B. et al. Harmful algal blooms and their effects in coastal seas of Northern Europe. Harmful Algae 102, 101989 (2021).

4. Murray, S.A. et al. A fish kill associated with a bloom of *Amphidinium carterae* in a coastal lagoon in Sydney, Australia. Harmful Algae 49, 19–28 (2015).

5. Gobler, C.J. Climate change and harmful algal blooms: insights and perspective. Harmful Algae 91, 101731 (2020).

6. Taylor, F.J.R., Hoppenrath, M. & Saldarriaga, J.F. Dinoflagellate diversity and distribution. Biodivers. Conserv. 17, 407–418 (2008).

7. LaJeunesse, T.C. et al. Systematic revision of Symbiodiniaceae highlights the antiquity and diversity of coral endosymbionts. Curr. Biol. 28, 2570–2580 (2018).

8. Stoecker, D.K., Hansen, P.J., Caron, D.A. & Mitra, A. Mixotrophy in the marine plankton. Annu. Rev. Mar. Sci. 9, 311–335 (2017).

9. Rädecker, N. et al. Heat stress destabilizes symbiotic nutrient cycling in corals. Proc. Natl. Acad. Sci. U. S. A. 118, e2022653118 (2021).

10. Johnson, J.G., Morey, J.S., Neely, M.G., Ryan, J.C. & Van Dolah, F.M. Transcriptome remodeling associated with chronological aging in the dinoflagellate, *Karenia brevis*. Mar. Genom. 5, 15–25 (2012).

11. Shi, X. et al. Transcriptomic and microRNAomic profiling reveals multi-faceted mechanisms to cope with phosphate stress in a dinoflagellate. ISME J. 11, 2209–2218 (2017).

12. Wang, X. et al. Transcriptome sequencing of a toxic dinoflagellate, *Karenia mikimotoi* subjected to stress from solar ultraviolet radiation. Harmful Algae 88, 101640 (2019).

13. LaJeunesse, T.C., Lambert, G., Andersen, R.A., Coffroth, M.A. & Galbraith, D.W. *Symbiodinium* (Pyrrhophyta) genome sizes (DNA content) are smallest among dinoflagellates. J. Phycol. 41, 880–886 (2005).

14. Saad, O.S. et al. Genome size, rDNA copy, and qPCR assays for Symbiodiniaceae. Front. Microbiol. 11, 847 (2020).

15. Lin, S. Genomic understanding of dinoflagellates. Res. Microbiol. 162, 551–569 (2011).

16. Wisecaver, J.H. & Hackett, J.D. Dinoflagellate genome evolution. Annu. Rev. Microbiol. 65, 369–387 (2011).

17. Shoguchi, E. et al. Draft assembly of the *Symbiodinium minutum* nuclear genome reveals dinoflagellate gene structure. Curr. Biol. 23, 1399–1408 (2013).

18. Lin, S. et al. The *Symbiodinium kawagutii* genome illuminates dinoflagellate gene expression and coral symbiosis. Science 350, 691–694 (2015).

19. Aranda, M. et al. Genomes of coral dinoflagellate symbionts highlight evolutionary adaptations conducive to a symbiotic lifestyle. Sci. Rep. 6, 39734 (2016).

20. Liu, H. et al. *Symbiodinium* genomes reveal adaptive evolution of functions related to coral-dinoflagellate symbiosis. Commun. Biol. 1, 95 (2018).

21. González-Pech, R.A. et al. Comparison of 15 dinoflagellate genomes reveals extensive sequence and structural divergence in family Symbiodiniaceae and genus *Symbiodinium*. BMC Biol. 19, 73 (2021).

22. Beedessee, G. et al. Integrated omics unveil the secondary metabolic landscape of a basal dinoflagellate. BMC Biol. 18, 139 (2020).

23. Camp, E.F. et al. Proteome metabolome and transcriptome data for three Symbiodiniaceae under ambient and heat stress conditions. Scientific Data 9, 153 (2022).

24. Dougan, K.E. et al. Genome-powered classification of microbial eukaryotes: focus on coral algal symbionts. Trends Microbiol. 30, 831–840 (2022).

25. Zaheri, B. & Morse, D. Assessing nucleic acid binding activity of four dinoflagellate cold shock domain proteins from *Symbiodinium kawagutii* and *Lingulodinium polyedra*. BMC Molecular and Cell Biology 22, 27 (2021).

26. Wong, J.T.Y. Architectural organization of dinoflagellate liquid crystalline chromosomes. Microorganisms 7, 27 (2019).

27. Levin, R.A. et al. Sex, scavengers, and chaperones: transcriptome secrets of divergent *Symbiodinium* thermal tolerances. Mol. Biol. Evol. 33, 2201–2215 (2016).

28. Liew, Y.J., Li, Y., Baumgarten, S., Voolstra, C.R. & Aranda, M. Condition-specific RNA editing in the coral symbiont *Symbiodinium microadriaticum*. PLoS Genet. 13, e1006619 (2017).

29. Mohamed, A.R. et al. Dual RNA-sequencing analyses of a coral and its native symbiont during the establishment of symbiosis. Mol. Ecol. 29, 3921–3937 (2020).

30. Zhang, H. et al. Spliced leader RNA trans-splicing in dinoflagellates. Proc. Natl. Acad. Sci. U. S. A. 104, 4618–4623 (2007).

31. Slamovits, C.H. & Keeling, P.J. Widespread recycling of processed cDNAs in dinoflagellates. Curr. Biol. 18, R550–552 (2008).

32. Mungpakdee, S. et al. Massive gene transfer and extensive RNA editing of a symbiotic dinoflagellate plastid genome. Genome Biol. Evol. 6, 1408–1422 (2014).

33. Velikova, V. & Larsen, J. The *Prorocentrum cordatum/Prorocentrum minimum* taxonomic problem. Grana 38, 108–112 (1999).

34. Zhang, F., Li, M., Glibert, P.M. & Ahn, S.H. A three-dimensional mechanistic model of *Prorocentrum minimum* blooms in eutrophic Chesapeake Bay. Sci. Total Environ. 769, 144528 (2021).

35. Khanaychenko, A.N., Telesh, I.V. & Skarlato, S.O. Bloom-forming potentially toxic dinoflagellates *Prorocentrum cordatum* in marine plankton food webs. Protistology 13, 95–125 (2019).

36. Seebens, H., Schwartz, N., Schupp, P.J. & Blasius, B. Predicting the spread of marine species introduced by global shipping. Proc. Natl. Acad. Sci. U. S. A. 113, 5646–5651 (2016).

37. Stephens, T.G. et al. Genomes of the dinoflagellate *Polarella glacialis* encode tandemly repeated single-exon genes with adaptive functions. BMC Biol. 18, 56 (2020).

38. Dougan, K.E. et al. Whole-genome duplication in an algal symbiont serendipitously confers thermal tolerance to corals. bioRxiv, 2022.2004.2010.487810 (2022).

39. Chen, Y. et al. Improved *Cladocopium goreaui* genome assembly reveals features of a facultative coral symbiont and the complex evolutionary history of dinoflagellate genes. Microorganisms 10, 1662 (2022).

40. John, U. et al. An aerobic eukaryotic parasite with functional mitochondria that likely lacks a mitochondrial genome. Sci. Adv. 5, eaav1110 (2019).

41. Stephens, T.G., Ragan, M.A., Bhattacharya, D. & Chan, C.X. Core genes in diverse dinoflagellate lineages include a wealth of conserved dark genes with unknown functions. Sci. Rep. 8, 17175 (2018).

42. Singh, A. et al. DIABLO: an integrative approach for identifying key molecular drivers from multi-omics assays. Bioinformatics 35, 3055–3062 (2019).

43. Salvucci, M.E. & Crafts-Brandner, S.J. Inhibition of photosynthesis by heat stress: The activation state of Rubisco as a limiting factor in photosynthesis. Physiol. Plant. 120, 179–186 (2004).

44. Schroda, M., Hemme, D. & Mühlhaus, T. The *Chlamydomonas* heat stress response. Plant J. 82, 466–480 (2015).

45. Verma, A. et al. The genetic basis of toxin biosynthesis in dinoflagellates. Microorganisms 7, 222 (2019).

46. Shi, X., Zhang, H. & Lin, S. Tandem repeats, high copy number and remarkable diel expression rhythm of form II RuBisCO in *Prorocentrum donghaiense* (Dinophyceae). PLoS ONE 8, e71232 (2013).

47. Lee, M.G. The 3’ untranslated region of the hsp 70 genes maintains the level of steady state mRNA in *Trypanosoma brucei* upon heat shock. Nucleic Acids Res. 26, 4025–4033 (1998).

48. Quijada, L., Soto, M., Alonso, C. & Requena, J.M. Identification of a putative regulatory element in the 3’-untranslated region that controls expression of HSP70 in *Leishmania infantum*. Mol. Biochem. Parasitol. 110, 79–91 (2000).

49. Zilka, A., Garlapati, S., Dahan, E., Yaolsky, V. & Shapira, M. Developmental regulation of heat shock protein 83 in *Leishmania*. 3’ processing and mRNA stability control transcript abundance, and translation id directed by a determinant in the 3’-untranslated region. J. Biol. Chem. 276, 47922–47929 (2001).

50. Zhang, H. & Lin, S. Complex gene structure of the form II RuBisCo in the dinoflagellate *Prorocentrum minimum* (Dinophyceae). J. Phycol. 39, 1160–1171 (2003).

51. Bruce, B.D. Chloroplast transit peptides: structure, function and evolution. Trends Cell Biol. 10, 440–447 (2000).

52. Lee, D.W. & Hwang, I. Evolution and design principles of the diverse chloroplast transit peptides. Mol. Cell 41, 161–167 (2018).

53. Csurös, M., Rogozin, I.B. & Koonin, E.V. Extremely intron-rich genes in the alveolate ancestors inferred with a flexible maximum-likelihood approach. Mol. Biol. Evol. 25, 903–911 (2008).

54. Rogozin, I.B., Carmel, L., Csuros, M. & Koonin, E.V. Origin and evolution of spliceosomal introns. Biol. Direct 7, 11 (2012).

55. van der Burgt, A., Severing, E., De Wit, P.J.G.M. & Collemare, J. Birth of new spliceosomal introns in fungi by multiplication of introner-like elements. Curr. Biol. 22, 1260–1265 (2012).

56. Huff, J.T., Zilberman, D. & Roy, S.W. Mechanism for DNA transposons to generate introns on genomic scales. Nature 538, 533–536 (2016).

57. Roy, S.W., Gozashti, L., Bowser, B.A., Weinstein, B.N. & Larue, G.E. Massive intron gain in the most intron-rich eukaryotes is driven by introner-like transposable elements of unprecedented diversity and flexibility. bioRxiv, 2020.2010.2014.339549 (2020).

58. Sanchez-Garcia, S., Wang, H. & Wagner-Döbler, I. The microbiome of the dinoflagellate *Prorocentrum cordatum* in laboratory culture and its changes at higher temperatures. Front. Microbiol. 13, 952238 (2022).

59. Wohlrab, S., Iversen, M.H. & John, U. A molecular and co-evolutionary context for grazer induced toxin production in *Alexandrium tamarense*. PLoS ONE 5, e15039 (2010).

60. Kang, H.C. et al. Comparative transcriptome analysis of the phototrophic dinoflagellate *Biecheleriopsis adriatica* grown under optimal temperature and cold and heat stress. Front. Mar. Sci. 8 (2021).

61. Gallaher, S.D. et al. Widespread polycistronic gene expression in green algae. Proc. Natl. Acad. Sci. U. S. A. 118, e2017714118 (2021).

62. Strassert, J.F.H., Irisarri, I., Williams, T.A. & Burki, F. A molecular timescale for eukaryote evolution with implications for the origin of red algal-derived plastids. Nat. Commun. 12, 1879 (2021).

63. Guillard, R.R.L. & Hargraves, P.E. *Stichochrysis immobilis* is a diatom, not a chrysophyte. Phycologia 32 (1993).

64. Michelle Wood, A., Everroad, R.C. & Wingard, L.M. Measuring growth rates in microalgal cultures. In Algal Culturing Techniques (ed. Andersen, R.A.) 269–286 (Elsevier Academic Press, Burlington, MA, 2005).

65. Levi-Setti, R., Gavrilov, K.L. & Rizzo, P.J. Divalent cation distribution in dinoflagellate chromosomes imaged by high-resolution ion probe mass spectrometry. Eur. J. Cell Biol. 87, 963–976 (2008).

66. Zimin, A.V. et al. Hybrid assembly of the large and highly repetitive genome of *Aegilops tauschii*, a progenitor of bread wheat, with the MaSuRCA mega-reads algorithm. Genome Res. 27, 787–792 (2017).

67. Xue, W. et al. L_RNA_scaffolder: scaffolding genomes with transcripts. BMC Genomics 14, 604 (2013).

68. Wang, M. & Kong, L. pblat: a multithread blat algorithm speeding up aligning sequences to genomes. BMC Bioinformatics 20, 28 (2019).

69. Laetsch, D.R. & Blaxter, M.L. BlobTools: interrogation of genome assemblies. F1000Res. 6, 1287 (2017).

70. Chen, S., Zhou, Y., Chen, Y. & Gu, J. fastp: an ultra-fast all-in-one FASTQ preprocessor. Bioinformatics 34, i884–i890 (2018).

71. Grabherr, M.G. et al. Full-length transcriptome assembly from RNA-Seq data without a reference genome. Nat. Biotechnol. 29, 644–652 (2011).

72. Li, W. & Godzik, A. Cd-hit: a fast program for clustering and comparing large sets of protein or nucleotide sequences. Bioinformatics 22, 1658–1659 (2006).

73. Kim, D., Paggi, J.M., Park, C., Bennett, C. & Salzberg, S.L. Graph-based genome alignment and genotyping with HISAT2 and HISAT-genotype. Nat. Biotechnol. 37, 907–915 (2019).

74. Chen, Y., González-Pech, R.A., Stephens, T.G., Bhattacharya, D. & Chan, C.X. Evidence that inconsistent gene prediction can mislead analysis of dinoflagellate genomes. J. Phycol. 56, 6–10 (2020).

75. Li, H. Minimap2: pairwise alignment for nucleotide sequences. Bioinformatics 34, 3094–3100 (2018).

76. Haas, B.J. et al. Improving the *Arabidopsis* genome annotation using maximal transcript alignment assemblies. Nucleic Acids Res. 31, 5654–5666 (2003).

77. Remmert, M., Biegert, A., Hauser, A. & Söding, J. HHblits: lightning-fast iterative protein sequence searching by HMM-HMM alignment. Nat. Methods 9, 173–175 (2012).

78. Korf, I. Gene finding in novel genomes. BMC Bioinformatics 5, 59 (2004).

79. Hoff, K.J. & Stanke, M. Predicting genes in single genomes with AUGUSTUS. Curr. Protoc. Bioinform. 65, e57 (2019).

80. Borodovsky, M. & Lomsadze, A. Eukaryotic gene prediction using GeneMark.hmm-E and GeneMark-ES. Curr. Protoc. Bioinform. 35, 4.6.1–4.6.10 (2011).

81. Campbell, M.S., Holt, C., Moore, B. & Yandell, M. Genome annotation and curation using MAKER and MAKER-P. Curr. Protoc. Bioinform. 48, 4.11.11–14.11.39 (2014).

82. Haas, B.J. et al. Automated eukaryotic gene structure annotation using EVidenceModeler and the Program to Assemble Spliced Alignments. Genome Biol. 9, R7 (2008).

83. Mŕazek, J. & Xie, S. Pattern locator: a new tool for finding local sequence patterns in genomic DNA sequences. Bioinformatics 22, 3099–3100 (2006).

84. Farhat, S. et al. Rapid protein evolution, organellar reductions, and invasive intronic elements in the marine aerobic parasite dinoflagellate *Amoebophrya* spp. BMC Biol. 19, 1 (2021).

85. Johnson, L.K., Alexander, H. & Brown, C.T. Re-assembly, quality evaluation, and annotation of 678 microbial eukaryotic reference transcriptomes. GigaScience 8, giy158 (2019).

86. Emms, D.M. & Kelly, S. OrthoFinder: phylogenetic orthology inference for comparative genomics. Genome Biol. 20, 238 (2019).

87. Pachiadaki, M.G. et al. Charting the complexity of the marine microbiome through single-cell genomics. Cell 179, 1623–1635 (2019).

88. Katoh, K. & Standley, D.M. MAFFT multiple sequence alignment software version 7: improvements in performance and usability. Mol. Biol. Evol. 30, 772–780 (2013).

89. Capella-Gutiérrez, S., Silla-Martínez, J.M. & Gabaldón, T. trimAl: a tool for automated alignment trimming in large-scale phylogenetic analyses. Bioinformatics 25, 1972–1973 (2009).

90. Minh, B.Q. et al. IQ-TREE 2: new nodels and efficient methods for phylogenetic inference in the genomic era. Mol. Biol. Evol. 37, 1530–1534 (2020).

91. Rohart, F., Gautier, B., Singh, A. & Lê Cao, K.A. mixOmics: an R package for ‘omics feature selection and multiple data integration. PLoS Comp. Biol. 13, e1005752 (2017).

92. Zhou, G. et al. NetworkAnalyst 3.0: a visual analytics platform for comprehensive gene expression profiling and meta-analysis. Nucleic Acids Res. 47, W234–W241 (2019).

93. Voolstra, C.R. et al. Contrasting heat stress response patterns of coral holobionts across the Red Sea suggest distinct mechanisms of thermal tolerance. Mol. Ecol. 30, 4466–4480 (2021).

94. Ibarbalz, F.M. et al. Global trends in marine plankton diversity across Kingdoms of life. Cell 179, 1084–1097 (2019).

95. Alexander, M.A. et al. Projected sea surface temperatures over the 21st century: changes in the mean, variability and extremes for large marine ecosystem regions of Northern Oceans. Elementa 6, 9 (2018).

96. Paul, C., Mausz, M.A. & Pohnert, G. A co-culturing/metabolomics approach to investigate chemically mediated interactions of planktonic organisms reveals influence of bacteria on diatom metabolism. Metabolomics 9, 349–359 (2013).

97. Liao, Y., Smyth, G.K. & Shi, W. FeatureCounts: an efficient general purpose program for assigning sequence reads to genomic features. Bioinformatics 30, 923–930 (2014).

98. Hamilton, N.E. & Ferry, M. ggtern: Ternary diagrams using ggplot2. J. Stat. Softw. 87, 1–17 (2018).

99. Robinson Mark, D., McCarthy Davis, J. & Smyth Gordon, K. edgeR: a Bioconductor package for differential expression analysis of digital gene expression data. Bioinformatics 26, 139–140 (2010).

100. Supek, F., Bošnjak, M., Škunca, N. & Šmuc, T. REVIGO summarizes and visualizes long lists of Gene Ontology terms. PLoS ONE 6, e21800 (2011).

101. Aramaki, T. et al. KofamKOALA: KEGG Ortholog assignment based on profile HMM and adaptive score threshold. Bioinformatics 36, 2251–2252 (2020).

102. Yu, G., Wang, L.G., Han, Y. & He, Q.Y. ClusterProfiler: an R package for comparing biological themes among gene clusters. OMICS: J. Integrative Biol. 16, 284–287 (2012).

103. Bradford, M.M. A rapid and sensitive method for the quantitation of microgram quantities of protein utilizing the principle of protein-dye binding. Anal. Biochem. 72, 248–254 (1976).

104. Wöhlbrand, L., Rabus, R., Blasius, B. & Feenders, C. Influence of NanoLC column and gradient length as well as MS/MS frequency and sample complexity on shotgun protein identification of marine bacteria. J. Mol. Microbiol. Biotechnol. 27, 199–212 (2017).

105. Wöhlbrand, L. et al. Analysis of membrane-protein complexes of the marine sulfate reducer *Desulfobacula toluolica* Tol2 by 1D blue native-PAGE complexome profiling and 2D blue native-/SDS-PAGE. Proteomics 16, 973–988 (2016).

106. Neuhoff, V., Arold, N., Taube, D. & Ehrhardt, W. Improved staining of proteins in polyacrylamide gels including isoelectric focusing gels with clear background at nanogram sensitivity using Coomassie Brilliant Blue G-250 and R-250. Electrophoresis 9, 255–262 (1988).

107. Hiller, K. et al. Metabolite detector: comprehensive analysis tool for targeted and nontargeted GC/MS based metabolome analysis. Anal. Chem. 81, 3429–3439 (2009).

