## Extended Data for "Multi-omics analysis reveals the molecular response to heat stress in a “red tide” dinoflagellate"

**Extended Data Table 1.** Statistics of assembled genomes of *P. cordatum* and other dinoflagellates.

| Species | *Prorocentrum cordatum* | **Amphidinium gibbosum* | *Polarella glacialis* | *Symbiodium natans* | *Durusdinium trenchii* | *Cladocopium goreaui* | *Amoebophrya ceratii* |
|---|---|---|---|---|---|---|---|
| Isolate | CCMP1329 | Not defined | CCMP1383 | CCMP2548 | CCMP2556 | SCF055 | AT5.2 |
| Reference | This study | Beedessee et al.[22] | Stephens et al.[37] | Gonzalez-Pech et al.[21] | Dougan et al.[38] | Chen et al.[39] | John et al.[40] |
| Assembly size (Gbp) | 4.15 | 7.55 | 2.99 | 0.76 | 1.71 | 1.17 | 0.09 |
| Estimated genome size based on *k*-mers (Gbp) | 4.75 | 6.30 | 1.48 | 0.74 | 1.05 | 1.30 | 0.12 |
| Number of scaffolds | 22,724 | 4,221,750 | 33,494 | 2,855 | 29,137 | 6,843 | 2,351 |
| Genome scaffolds N50 (Kbp) | 349.2 | 150.4 | 170.3 | 610.5 | 774.3 | 353.9 | 84.0 |
| Maximum scaffold length (Mbp) | 3.56 | 1.36 | 2.17 | 3.40 | 4.57 | 7.38 | 0.54 |
| Genome GC-content (%) | 59.7 | 47.1 | 45.9 | 51.8 | 49.8 | 44.4 | 55.9 |
| BUSCO proteins recovered (%) [genome mode; alveolata_odb10] | 57.9 | 48.5 | 60.2 | 62.0 | 61.4 | 66.7 | 82.5 |

*: Values for *Amphidinium gibbosum* (not displayed in Figure 1) were derived directly from the published assembly version 1.0 at https://marinegenomics.oist.jp/amphidinium/viewer/download?project_id=83

**Extended Data Table 2.** Statistics of predicted genes in *P. cordatum* and other dinoflagellates.

| Species | | *Prorocentrum cordatum* | **Amphidinium gibbosum* | *Polarella glacialis* | *Symbiodium natans* | *Durusdinium trenchii* | *Cladocopium goreaui* | *Amoebophrya ceratii* |
|---|---|---|---|---|---|---|---|---|
| Isolate | | CCMP1329 | Not defined | CCMP1383 | CCMP2548 | CCMP2556 | SCF055 | AT5.2 |
| Reference | | This study | Beedessee et al.[22] | Stephens et al.[37] | Gonzalez-Pech et al.[21] | Dougan et al.[38] | Chen et al.[39] | John et al.[40] |
| Number of predicted genes | | 85,849 | 85,139 | 58,232 | 35,270 | 55,799 | 45,322 | 19,925 |
| Recovery of BUSCO proteins (%) [protein mode; alveolata_odb10] | | 61.4 | 45.0 | 70.2 | 74.9 | 69.6 | 82.4 | 86.5 |
| Genes with transcript support (%) | | 84.8 | 75.9 | 94.0 | 83.0 | 75.7 | 82.5 | 24.4 |
| Average gene length (bp) | | 24,462 | 26,201 | 16,206 | 8,780 | 15,334 | 15,745 | 2,772 |
| Average CDS length (bp) | | 2,798 | 1,193 | 1,230 | 1,660 | 1,647 | 2,018 | 1,964 |
| CDS GC-content (%) | | 65.9 | 54.9 | 57.8 | 58.16 | 55.69 | 54.24 | 60.8 |
| Number of exons per gene | | 11.7 | 8.0 | 11.6 | 15.7 | 16.7 | 17.2 | 3.4 |
| Average exon length (bp) | | 239.9 | 184.8 | 105.7 | 106 | 98.7 | 120.4 | 578.7 |
| Genes with introns (%) | | 83.7 | 92.7 | 73.8 | 85.5 | 93.1 | 95.9 | 71.3 |
| Number of introns per gene | | 9.8 | 7.0 | 10.6 | 14.7 | 15.7 | 16.2 | 2.4 |
| Average intron length (bp) | | 4709 | 3,732 | 1408 | 486 | 869 | 839 | 377 |
| Splice donor motif (%) | GT | 24.7 | 74.9 | 28.8 | 23.6 | 30.3 | 36.6 | 99.9 |
| | GC | 51.4 | 25.0 | 52.7 | 58 | 52.3 | 43.6 | <0.1 |
| | GA | 23.8 | 0.1 | 18.5 | 18.4 | 17.4 | 19.8 | 0 |
| Splice acceptor with AGG motif (%) | | 79.7 | 93.2 | 96.9 | 97.1 | 96.5 | 96.1 | 56.5 |
| Number of intergenic regions | | 48,574 | 47,727 | 35,271 | 33,042 | 47,452 | 39,720 | 17,856 |
| Average length of intergenic regions (bp) | | 26,278 | 26,756 | 21,625 | 11,585 | 13,222 | 7,388 | 1,522 |

*: Values for *Amphidinium gibbosum* (not displayed in Figure 1) were derived directly from the published gene models version 1.0 at https://marinegenomics.oist.jp/amphidinium/viewer/download?project_id=83

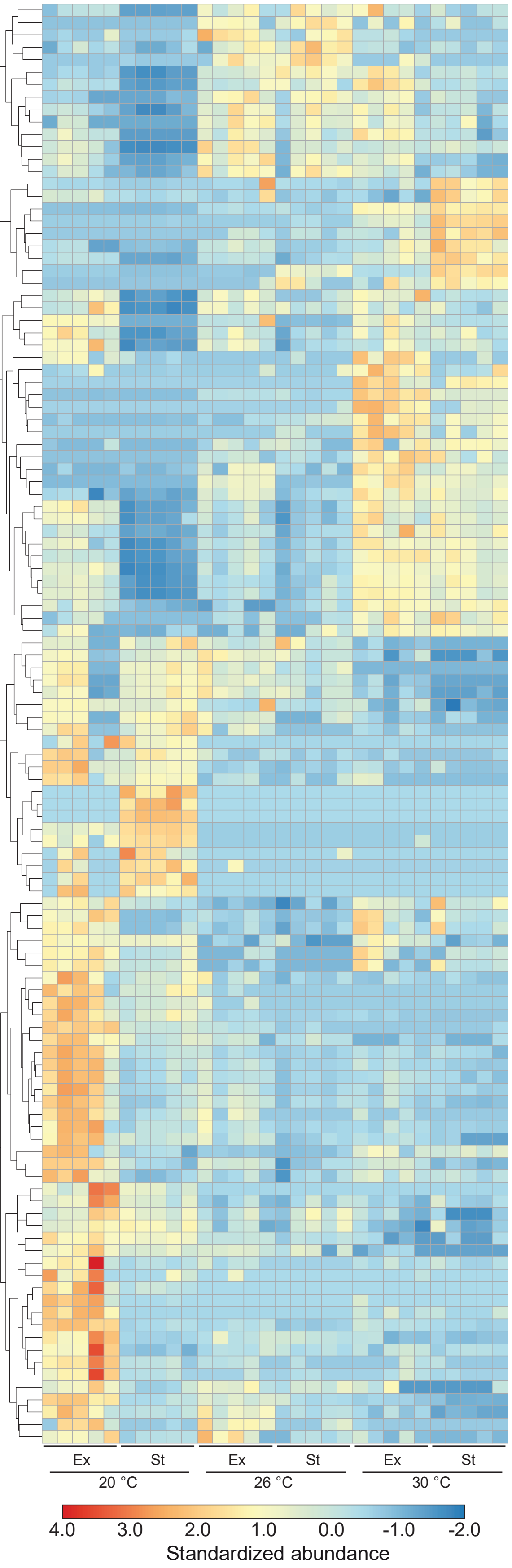


**Extended Data Fig. 1.** Heatmap of all identified metabolites ($p < 0.001$) at 20°C, 26°C and 30°C in *P. cordatum*, for Ex and St phase. Five biological replicates and three technical replicates were run per experiment. Relative intensity values are shown, and data were normalized to internal standard and cell count. Normalization for heatmap was done by using z-score, and significance was calculated using two-tailed T-test.

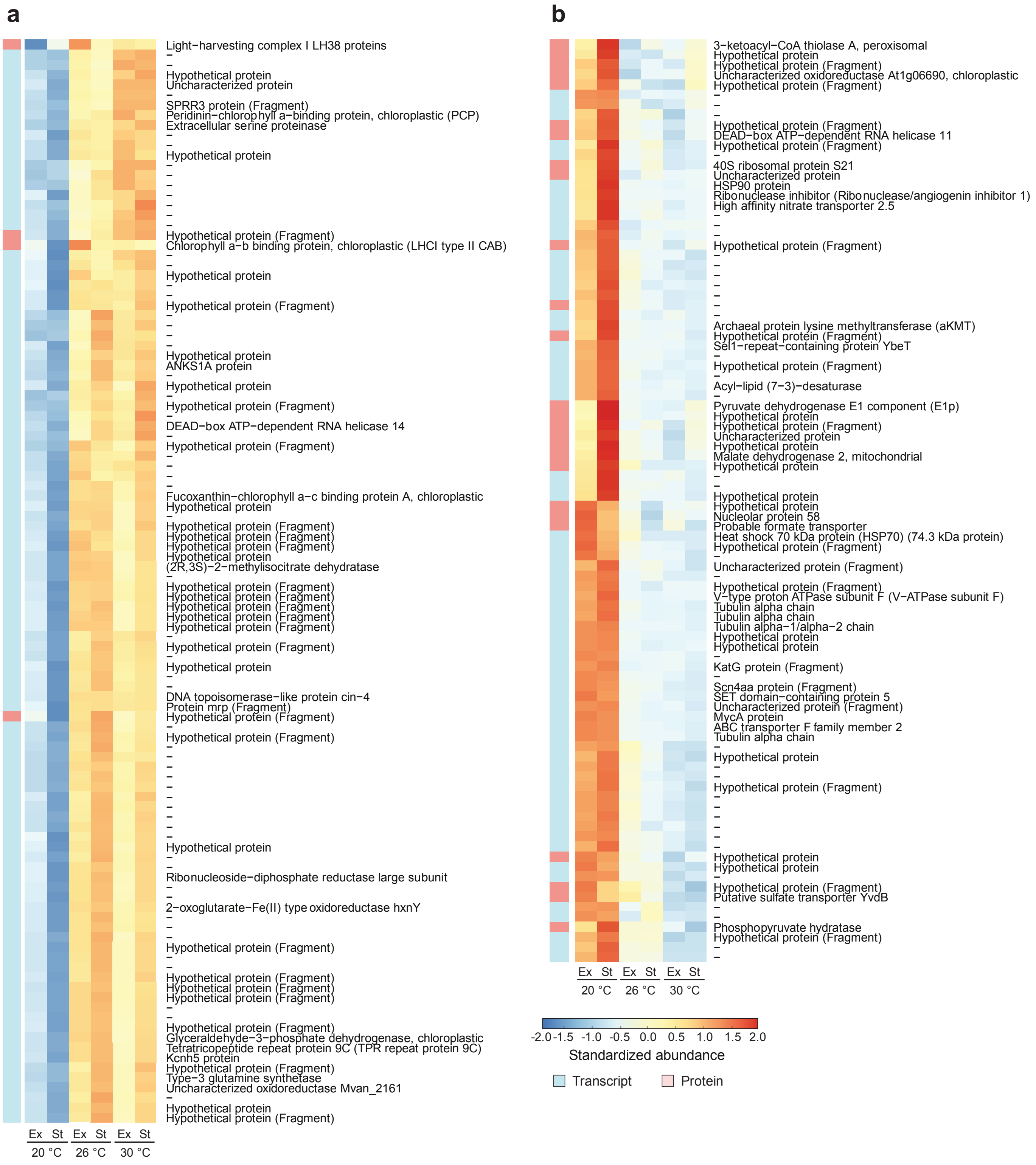


**Extended Data Fig. 2.** Heatmap of expression levels for the inferred multi-omics signature that is indicative of a general heat stress response (component 1), shown for values scaled across rows with proteins and transcripts that are (a) up-regulated and (b) down-regulated during heat stress.

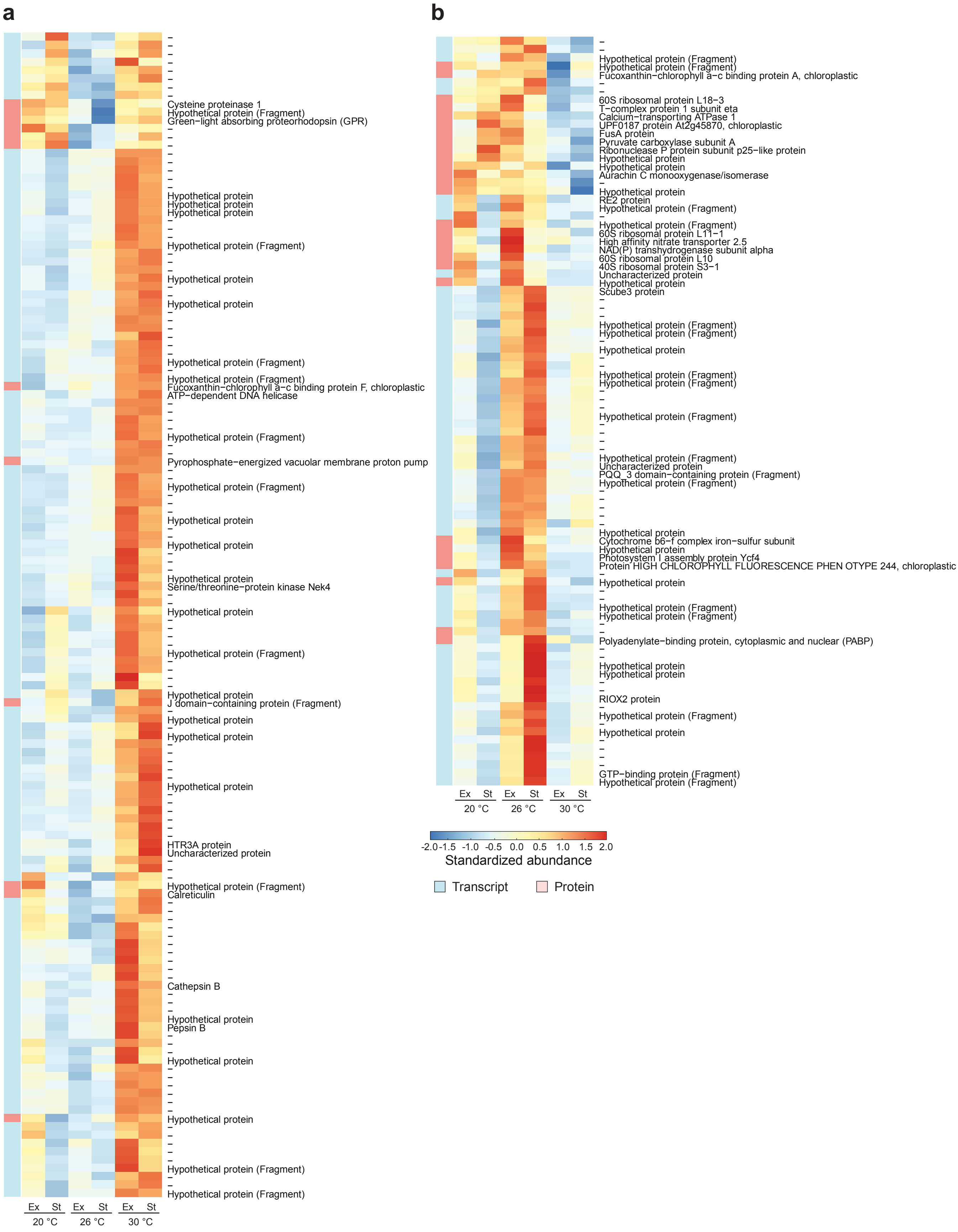


**Extended Data Fig. 3.** Heatmap of expression levels for the inferred multi-omics signature that is indicative of a heat-stress response specific to 26°C and to 30°C, shown for values scaled across rows with proteins and transcripts that are (a) up-regulated and (b) down-regulated during heat stress.

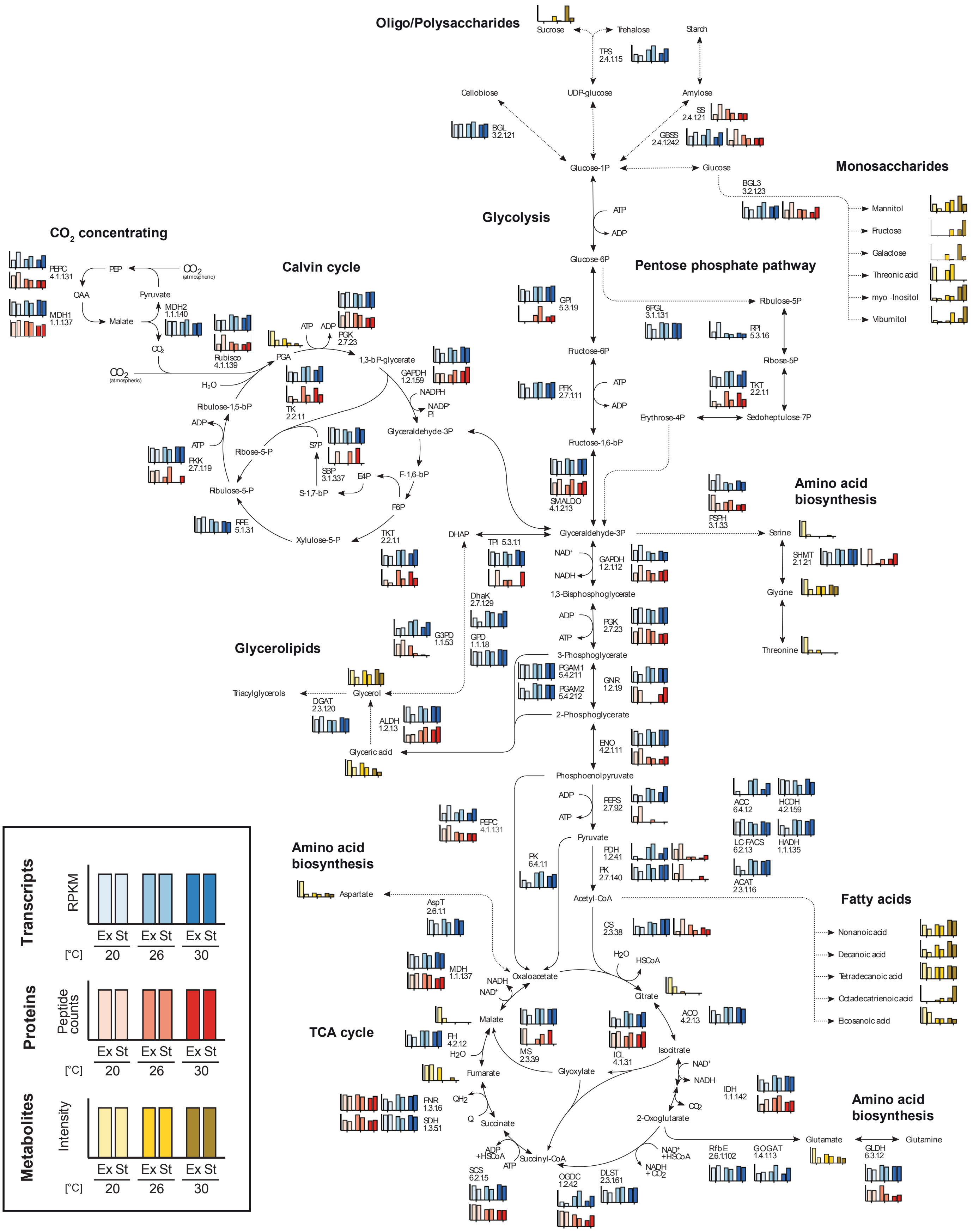


**Extended Data Fig. 4.** Detailed reconstruction of the central carbon metabolism of *P. cordatum* and the concerted responses to heat stress response at transcript (blue), protein (red) and metabolite (yellow) levels.

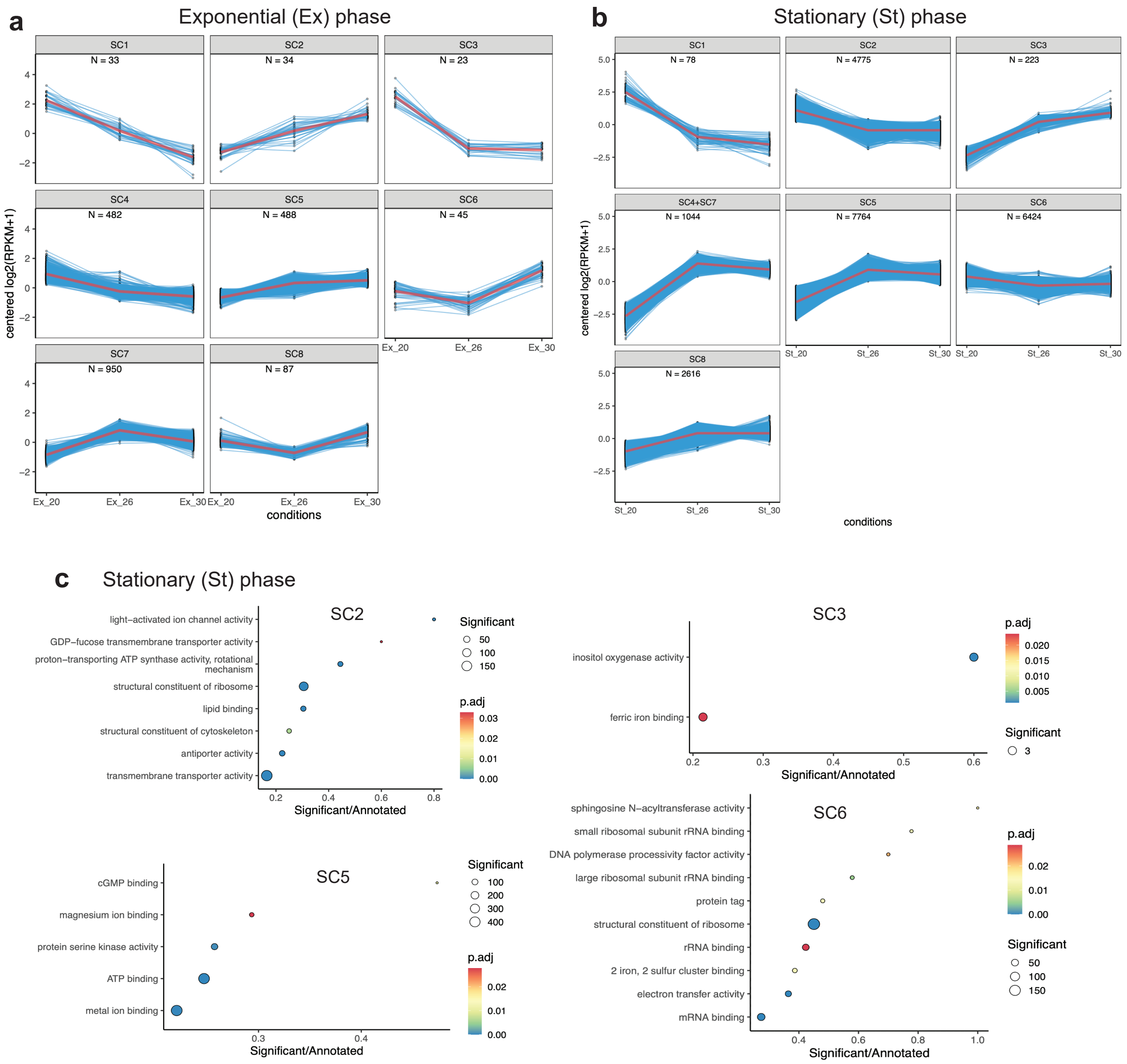


**Extended Data Fig. 5.** Clusters of genes with similar expression patterns across different conditions in (a) exponential (Ex) and (b) stationary (St) phase, and (c) enriched gene functions in the Molecular Function GO terms in SC2, SC3, SC5, and SC6 in the St phase.

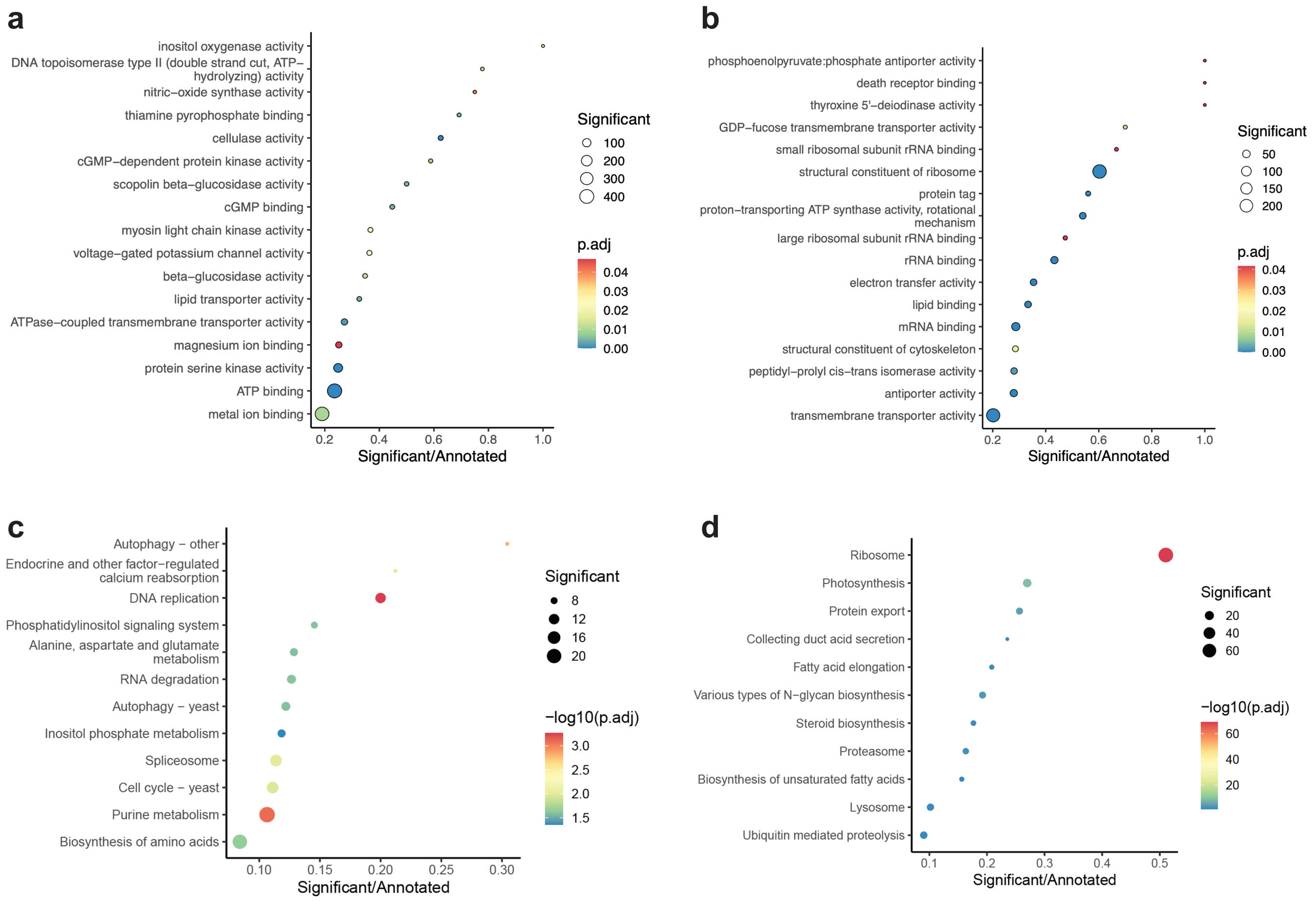


**Extended Data Fig. 6.** Enriched functions in differentially regulated genes in 26°C *versus* 20°C in stationary phase, showing enriched Molecular Function GO terms in (a) up-regulated and (b) down-regulated genes, and the enriched KEGG pathways in (c) up-regulated and (d) down-regulated genes.

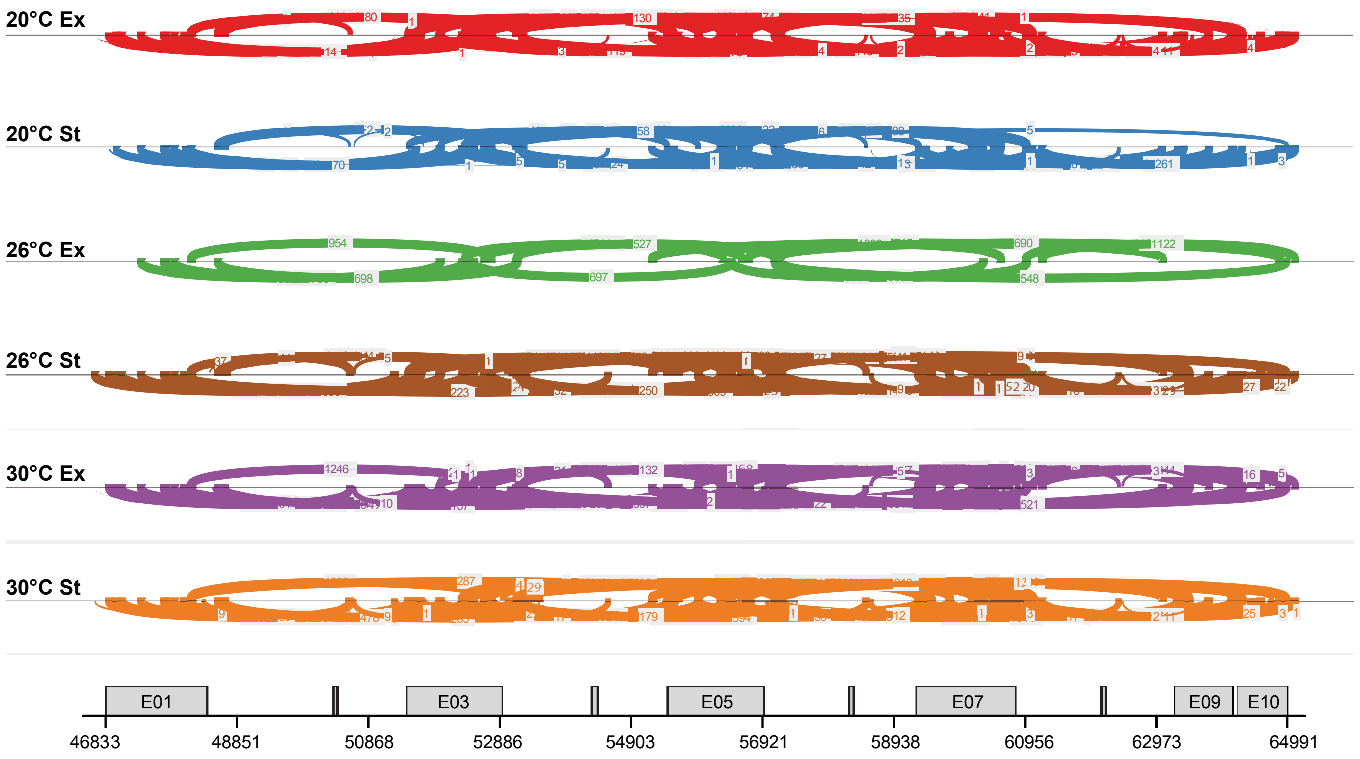


**Extended Data Fig. 7.** Sashimi plot across the HSP70 locus displaying read coverage with lines between exons indicating connections supported by >500 alignments.

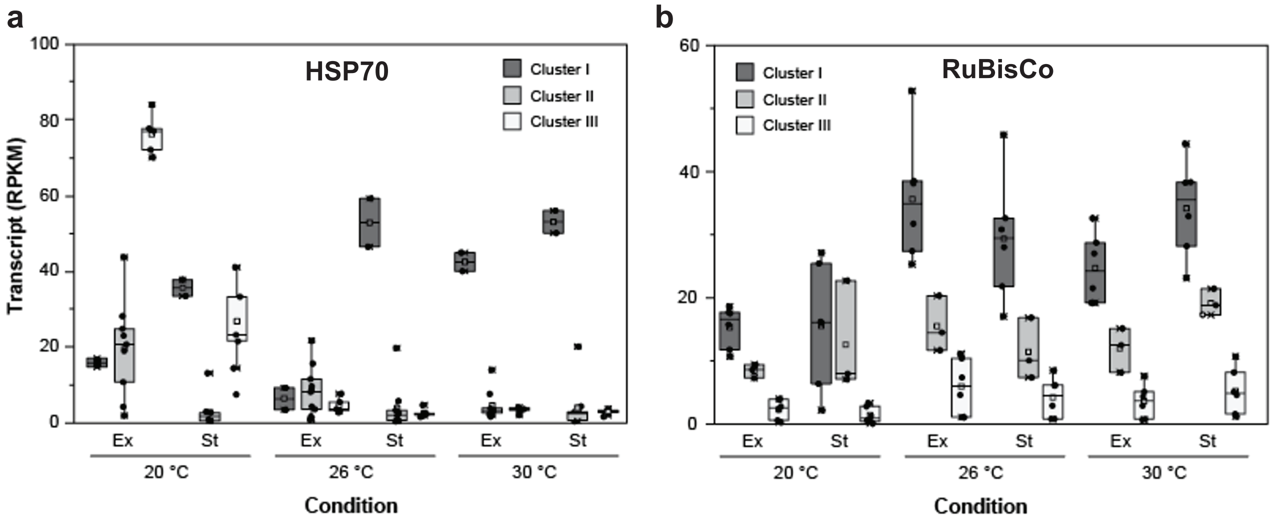


**Extended Data Fig. 8.** Transcript profiles of (a) HSP70 and (b) RuBisCo gene models based on clustering in Figures 6a and 6c.
