## Supplementary Note for "Multi-omics analysis reveals the molecular response to heat stress in a “red tide” dinoflagellate"

### *Analysis of Prorocentrum cordatum genome*

The *de novo* assembled haploid genome of *Prorocentrum cordatum* CCMP1329 (4.15 Gb) shares similar completeness (60.2% recovery of core alveolate genes) to other published dinoflagellate genomes (Extended Data Table 1). Most (61.0%) assembled bases were annotated as repetitive elements; 46.0% and 7.8% represent unknown and simple repeats, respectively (Fig. 1a).

Excluding the Order Suessiales (largely symbiotic) and the early-branching Syndiniales (parasitic), dinoflagellates are free-living and bloom-forming (Fig. 1b). To assess the implications of distinct ecological niches on genome evolution of dinoflagellates, we examined features of *P. cordatum* relative to five representative taxa chosen based on ecological niche, phylogenetic position, and quality of available genome data: the free-living *Polarella glacialis* CCMP1383[1] and *Symbiodinium natans* CCMP2548[2], the symbiotic *Durusdinium trenchii* CCMP2556[3] and *Cladocopium goreaui* SCF055[4], and the parasitic *Amoebophrya ceratii* AT5.2[5]. The draft genome of the other bloom-forming species (*Amphidinium gibbosum*)[6] is highly fragmented and less-complete (see next section below).

The haploid genome of *P. cordatum* (4.75 Gb; Supplementary Fig. 1), while smaller than that of *A. gibbosum*, is 3.2- and 39.6-fold larger than the genomes of *Po. glacialis* and the parasitic *A. ceratii*, respectively (Fig. 1e and Extended Data Table 1). *P. cordatum* has the highest G+C overall and in the protein-coding genes. The *A. gibbosum* genome has a moderate G+C content (47.1%) similar to that of *Po. glacialis* (45.9%), suggesting that high G+C is not common among free-living and/or bloom-forming taxa. G-C pairing is more thermally stable, thus the maintenance of high G+C in *P. cordatum* genome likely reflect an adaptation to changing environments in open oceans, *e.g.* up to 30°C in red tides. As comparison, the moderate G+C content in the *Po. glacialis* genome could be an adaptation to the extreme cold and relatively stable polar regions where the species is found. These conditions may counter-select against strong thermostability of its genes. More genome data from the free-living taxa are needed to determine if high G+C is a genome signature of free-living dinoflagellates.

The *P. cordatum* genome has the largest proportion of helitron repeats (1.2% of the assembled bases compared to <0.8% in the other genomes; Fig. 1e), whereas long interspersed nuclear elements and long-terminal repeats are expanded in the genomes of Suessiales including *Po. glacialis*. These elements are largely conserved in the *P. cordatum* genome (Supplementary Fig. 1 and Supplementary Note). A large proportion of the repetitive elements (46.0%) identified in the *P. cordatum* genome (Fig. 1a) are not classified into any known repeats. Long interspersed nuclear elements (LINEs) appear to be largely conserved in the genome, with estimated Kimura substitutions centered between 0 and 10 (Supplementary Fig. 1), whereas long-terminal repeats (LTRs) and the rolling circles of helitrons are slightly more diverged with Kimura substitutions centered between 10 and 20, indicating that most of these elements are likely active. In comparison, although some DNA transposons appear to be highly conserved (Kimura substitutions < 5), most of the others are highly diverged (Kimura substitutions >25) which suggests the accumulation of relic or inactive DNA transposons in the genome.

Compared to the other taxa, genomes of bloom-forming dinoflagellates encode more protein-coding genes (85,849 and 85,139 in *P. cordatum* and *A. gibbosum*, respectively) that are also longer, likely due to long introns: the mean gene length (and mean intron length) are 24.5 (4.7) Kb in *P. cordatum* and 26.2 (3.7) Kb in *A. gibbosum*, compared to 16.2 (1.4) Kb in *Po. glacialis* (Fig. 1e and Extended Data Table 2). Our results suggest a prevalence of introner elements in the genomes of free-living dinoflagellates (Supplementary Tables 1 and 2).

***Genome data from the bloom-forming Amphidinium gibbosum***

Genome data from another bloom-forming dinoflagellate species are available from *Amphidinium gibbosum*[6]; the assembled genome has a size of 7.5 Gbp that is approximately 1.2-fold larger than the estimated size at 6.3 Gbp based on sequence data or 6.4 Gbp based on flow cytometry. The published assembly (https://marinegenomics.oist.jp/amphidinium/viewer/download?project_id=83) contains 4,221,750 scaffolds with N50 scaffold length 150.4 Kb, with an overall G+C content of 47.03%; mean G+C content of the predicted coding sequences is higher at 54.88%. These metrics, derived directly from the published assembly, are slightly different from those reported[6], which were potentially based on subset of the assembled sequences filtered by an undefined length. In any case, these values indicate high fragmentation of the assembled genome sequences, and that repetitive genomics regions are not well resolved; this is a known issue with assemblies of this size generated using only Illumina short-read data. For example, the repeat content for *A. gibbosum* was reported to cover 2,257,532,404 bases[6] (29.9% of the total assembled 7,553,259,460 bases), in lower percentage than what we observed in *P. cordatum* (61% of assembled bases; Fig. 1a).

To further verify data comparability, the completeness of the assembled *A. gibbosum* genome was assessed using the same approach we did for all other genomes in the analysis, based on recovery of core conserved alveolate genes (alveolata_odb10) from the BUSCO dataset. Our analysis yielded 48.5% recovery of these genes in the genome (Extended Data Table 1), and 45.0% among the predicted proteins (Extended Data Table 2). These values contrast with our expectation that BUSCO recovery from protein data is always greater than that from genome data. These values are also lower than the other six genome datasets, *i.e.* mean recovery of 65.5% from genome data (Extended Data Table 1), and 75.6% from protein data (Extended Data Table 2). For these reasons and to ensure data comparability, we cautiously excluded this assembly from our detailed genome comparison of dinoflagellates from distinct ecological niches, and included only bioinformatic results where relevant (Extended Data Tables 1 and 2).

***Introner elements***

Introner elements (IE) are introns that contain inverted repeats with short direct repeats at both ends, often resulting in hairpin structures[7, 8]. First identified in the green algae *Micromonas pusilla*[8], IEs act as non-autonomous DNA transposons and may have contributed to the rapid, extensive intron gain in some green algae and fungi[9, 10]. Here we found evidence of IEs in 2.4% of all introns in *P. cordatum* (Supplementary Table 2 and Fig. 1e). This percentage compares to 1.3% in *Po. glacialis* and < 1% in all lineages of Symbiodiniaceae, which are predominantly symbiotic. We found the highest percentages (10-11%) of genes containing IEs in the free-living dinoflagellates *Po. glacialis* and *P. cordatum*. The equivalent percentages in Symbiodiniaceae are between 1.4–5.9%. IEs in *P. cordatum* tended to occur in introns slightly smaller (mean 3.4 Kb) than the overall intron size (mean 4.1 Kb). When analyzing the functions of the 8,934 genes carrying IE elements, we found that the majority encoded proteins with unknown functions. However, genes encoding well characterized enzymes from KEGG pathways and other proteins (*e.g.* photosystem II reaction center protein H, calmodulin, ultraviolet receptors, and pentatricopeptide repeats) were also among them (Supplementary Table 1). We hypothesize that IEs are a prominent signature in the genomes of free-living dinoflagellates, and the reduced proportion of IEs among the introns in the family Symbiodiniaceae is likely a derived feature resulting from selection for symbiosis with diverse hosts[11].

***Duplication enhanced diversity of gene functions in a bloom-forming dinoflagellate***

We assigned putative functions to predicted proteins in the *P. cordatum* genome based on shared sequence similarity covering ≥ 70% of full-length sequences in the UniProt database (Supplementary Table 3; see Methods). This stringent approach ensures high confidence in the annotated functions, given the scarcity of dinoflagellate sequences in public repositories and their expected high sequence divergence[2, 12]. Of the 85,849 proteins in *P. cordatum*, 41,078 (47.8%) were given an annotation, 20,897 (24.3%) with Gene Ontology (GO) terms; similar proportions were observed in other genomes (Supplementary Table 4), with many genes (*e.g.* 52.15% for *P. cordatum* and 62.83% for *A. ceratii*) coding for functions yet to be discovered[13] or being distantly related homologs of sequences in UniProt. This expanded gene repertoire highlights lineage-specific innovation encompassed by conserved "dark" genes in dinoflagellates.

We assessed gene functions in *P. cordatum* relative to the other five genomes based on relative abundance of GO terms annotated per genome (Fig. 2a) and the total count of GO terms (Supplementary Fig. 2; see Methods). Compared to the other genomes, biological processes such as lipid and carbohydrate metabolic processes, recombination, integration, and repair of DNA, proteolysis, as well as signal transduction were more abundant in *P. cordatum* (Fig. 2a). A similar pattern was observed in molecular functions related to transmembrane transport, methyltransferase, oxidoreductase, DNA binding, protein kinase, serine-type endopeptidase, and hydrolase (Fig. 2a). For cellular components, the dominant functions were related to extracellular region and integral component of membrane (Fig. 2a). The free-living, cold-adapted *Po. glacialis* exhibited markedly different gene functions, suggesting that the highly abundant functions in *P. cordatum* are specific to this HAB-forming dinoflagellate. The parasitic (non-photosynthetic) *A. ceratii* contains only 2,799 genes with annotated GO terms, compared to 20,897 in *P. cordatum* (Supplementary Table 4), therefore the relative abundance of its functions is not comparable to the other genomes.

Based on analysis of collinear gene blocks, we found no evidence of segmental duplication, with 55,664 (64.8% of 85,849) genes identified as dispersed duplicates (Supplementary Table

5 and Fig. 1e). To investigate the impact of gene duplication on function, we assessed the enrichment of functions among dinoflagellate genes of distinct duplication modes (*i.e.* dispersed, proximal and tandem; see Methods). In *P. cordatum*, gene functions related to transmembrane transport, homeostasis, and organelle assembly are significantly enriched ($p \leq 0.01$) among the dispersed duplicates (Fig. 2b). Proximal duplicates are enriched in functions related to immune response and photosynthesis, whereas tandem duplicates are enriched in functions implicated in metabolic processes (*e.g.* tricarboxylic acid cycle [TCA]) and binding of biomolecules/ions (Fig. 2b and Supplementary Table 6). These GO terms are not significantly enriched ($p > 0.05$) in any of the duplication modes in the other genomes, although GO terms related to transmembrane and cation transport are also prominent in the free-living *Po. glacialis* and *S. natans*, implicating >100 genes in each genome (Fig. 2b). These results demonstrate that distinct duplication modes have shaped the evolution of *P. cordatum* genes and their functions related to adaptation.

***Tandemly repeated genes***

Tandemly repeated genes have previously been reported in the genomes of the polar dinoflagellate *Polarella glacialis*, particularly for genes encoding functions essential for survival, *e.g.* chlorophyll-binding proteins for photosynthesis and proteins containing ice-binding domain[1]. Tandemly repeated genes have been postulated to have fewer introns than the other genes in the bloom-forming dinoflagellate *Amphidinium carterae*[14]. In *P. glacialis*, many of these are bacterial like single-exon genes, thus this may enhance transcription efficiency of these genes as polycistronic mRNAs.

We found 2028 genes that appear in 876 blocks of tandemly repeated genes (Supplementary Table 5), with repetitions ranging from 2 to a maximum of 7 (Supplementary Table 6). On average these genes have 2.8 introns, compared to the overall average number of 9.8 introns among all *P. cordatum* genes. Of the 2028 tandemly repeated genes, 750 (37.0%) have single exons, more than twice the percentage of single exons genes found among the total number of genes (16.3%). The majority of the single exon genes (479 genes) are recovered in 205 blocks. Therefore, in *P. cordatum* we found a much smaller percentage of tandemly repeated genes and a much smaller number of repetitions compared to other free-living dinoflagellates, but the tandemly repeated genes shared the lower density of introns and the enrichment of single-exon genes with previous studies[1, 14].

***Horizontal gene transfer***

To assess the impact of horizontal gene transfer (HGT) of bacterial genes on the evolution of *P. cordatum* genes, we assessed shared sequence similarity between the predicted protein sequences of *P. cordatum* and those of abundant but mostly uncultivated bacterial taxa from the ocean plankton. The GORG-Tropics[15] is a comprehensive database of single-cell assembled genomes (SAGs) of prokaryotes collected from seawater samples. We used 688,212 protein sequences from 543 SAGs that are ≥ 85% complete (Supplementary Table 30). We searched for homologous sequences for each *P. cordatum* protein against a customized protein-sequence database that consists of 3,547,582 sequences: the *P. cordatum* proteins, 2,773,521 proteins from 82 other eukaryotes (Supplementary Table 29), and the 688,212 proteins from 543 SAGs.

We found 47 genes to have bacterial top hits (excluding *Prorocentrum* hits) in GORG-Tropics ($E \leq 10^{-5}$) (Supplementary Table 7), of which 10 (21%) are encoded by single-exon genes. This percentage is higher than that of single-exons genes in the genome of *P. cordatum* (14,006 single-exon genes, 16% of the total 85,849 genes). Nearly half (23, 49%) of these genes encode unknown functions, with few encoding functions related to biosynthesis and transport of antibiotics (Supplementary Table 7). The most frequently found potential source of those genes was SAR11, the most abundant but largely uncultivated marine group of bacteria.

Using the same database, we inferred 3,207,539 homologous protein sets. Of these, 1711 contain sequences from GORG-Tropics and *P. cordatum*. The 47 proteins are distributed in 21 homologous sets; we consider these sets represent evidence for putative HGT history in the *P. cordatum* genes, acquired from a bacterial source. Supplementary Fig. 4 shows the phylogenetic tree for the protein sequences that encode putative hydantoin racemase, which is part of the aspartate or glutamate racemase family. The gene is implicated in biosynthesis of peptidoglycan in bacteria, and some peptide-based antibiotics *e.g.* gramicidin S. Although the function of this protein homolog in *P. cordatum* is unclear, the bacterial origin of this gene is supported by a strongly supported (bootstrap support 87%) clade implicating proteobacteria of the SAR116 lineage that are ubiquitous but mostly uncultivated in the global oceans.

#### *Transcriptome landscape of P. cordatum*

We generated ~110 million reads of RNA-Seq data for each sample using 150 bp paired end chemistry on an Illumina NovaSeq 6000 platform. After quality control 108,209,172 ± 17,076,245 reads were retained per sample. Reads were mapped against the assembled reference genome using HISAT2 v2.2.1. To compute the gene expression, the uniquely mapped reads were counted for each gene by summing up the number of reads mapped to all its exons with featureCounts[16]. In total, 62,580 genes out of 85,849 gene models were expressed at an average read count ≥1 for all samples. Based on gene expression, the principal component analysis (PCA) clearly separated the 20°C samples from the temperature stressed samples, while the 26°C and 30°C conditions were highly similar (Supplementary Fig. 5a). The three replicate samples from each condition clustered closely together, except for one sample at 20°C in the stationary (St) growth phase that was separated from its replicates on the second principal component. Since only 7.34% of total variation was explained by the second principal component, we decided to include this replicate in the analysis.

#### *Transcriptome response to heat stress*

To demonstrate the differential regulation of pathways between different temperatures, we annotated genes with KEGG ortholog (KO) genes using KOfam[17]. About 20,637 genes could be annotated, with 6,589 unique KO genes representing around half of mapped reads (Supplementary Table 19); differentially expressed KO genes between different conditions were identified. In the St phase, 1,588 KO genes were differentially regulated between 26°C and 20°C, while 1,258 KO genes between 30 and 20. For the exponential (Ex) phase, we detected only 47 differentially regulated KO genes between 26°C and 20°C, 15 between 30°C and 20°C. A KEGG pathway enrichment analysis was performed on those differentially expressed KO genes. KEGG pathways modules including biosynthesis of amino acids, spliceosome, DNA replication, RNA degradation, various vitamin B complex biosynthesis and metabolism were up-regulated in response to heat stress (Extended Data Fig. 6c, Supplementary Table 20). Ribosome, photosynthesis, and steroid biosynthesis were down-regulated at higher temperatures compared to 20°C (Extended Data Fig. 6d, Supplementary Table 20).

A clustering analysis based on the gene expression showed eight (in Ex phase) or seven (in St phase) patterns in superclusters (SCs), of up- and down-regulated genes under the various conditions (Fig. 5d, Extended Data Fig. 5, and Supplementary Tables 16 and 17). As shown in Extended Data Fig. 5c, GO term enrichment analysis revealed that lipid binding, transmembrane transporter and structural constituent of cytoskeleton were enriched in SC2 of St phase. In comparison, Inositol oxygenase activity and ferric iron binding were enriched in SC3, cGMP/ATP binding and metal ion binding were enriched in SC5, whereas ribosome, rRNA binding, mRNA binding, protein tag and electron transfer activity were enriched in cluster SC6 (Extended Data Fig. 5c).

#### *Functional enrichment of dispersed genes with high level of duplication*

Using OrthoFinder, we identify homologous protein sets that consist of paralogs likely originated from gene duplication that share similar function. Since dispersed duplicates constitute most of all *P. cordatum* genes, we examined the functions of the dispersed genes with high duplication level. Among the highly duplicated homologous sets that contain ≥20 dispersed gene copies, GO terms including response to UV-B, DNA integration, proteolysis, RNA-mediated transposition, chlorophyll binding, ribulose-bisphosphate carboxylase activity, ATP-binding, photoreceptor activity, calcium-sodium antiporter, protein serine kinase activity, photosystem II, light-harvesting complex, mitochondrion, plastid, and microtubule were significantly enriched (Supplementary Fig. 8).

#### *Spliced leader sequences in P. cordatum transcripts*

Dinoflagellate spliced leader (dinoSL) is a conserved 22-nucleotide sequence (DCCGTAGCCATTTTGGCTCAAG, where D = T, A, or G) that is known to be added to the 5′-end of transcribed pre-mRNAs to yield mature mRNAs. We follow Stephens et al.[1] to access the presence of conserved dinoSL among *P. cordatum* transcripts. To maximize recovery of dinoSL independent from genome sequences, in this analysis we used the *de novo* assembled transcriptome, for which the redundant sequences were removed based on shared sequence identity ≥ 98% (see *Transcriptome Assembly* in Methods). Of the 1,776,418 transcripts we analyzed, 84,841 (4.78%) contain the dinoSL in the 5′-region, and additional 2,994 (0.17%) contain the dinoSL with one or more trailing relic dinoSLs: 2,793 with one, 200 with two, and 1 with three. Relic DinoSL sequences are expected when dinoSL-containing transcripts that were integrated back into the genome[18] were expressed and *trans*-spliced with a new dinoSL. Therefore, the presence of these relics may indicate high transcription activity for genes. Our results suggest that dinoSL is not present in all transcripts, as previously reported in *Po. glacialis*[1]. The dinoSL-containing transcripts implicate 17,214 (20.1% of 85,849) gene models (Supplementary Table 24), with s9118_g66601 (encoding for an unknown function) has the most dinoSL-containing transcripts (204), in addition to 236 with one relic, 10 with two relics, and 1 with three relics, indicating a high transcription activity. Other gene models associated with abundant dinoSL-containing transcripts include those coding for various subunits of 40S and 60S ribosomal proteins, and homoaconitate hydratase that is involved in lysine biosynthesis (Supplementary Table 24). Of the 17,214 gene models for which dinoSL-containing transcripts were found, 2,643 (15.4%) are single-exon genes; this compares to 14,006 (16.3%) single-exon genes among the total 85,849 gene models. DinoSL-containing transcripts implicating genes coding for HSP70 lends further support to the role of *trans*-splicing in converting polycistronic transcripts to individual (monocistronic) sequences. Although the potential functional bias for dinoSL-containing transcripts remains to be investigated with more data from other taxa, these results confirm the layer of transcription complexity in addition to differential RNA editing and exon usage in *P. cordatum* and other dinoflagellates.

### References


1. Stephens, T.G. et al. Genomes of the dinoflagellate *Polarella glacialis* encode tandemly repeated single-exon genes with adaptive functions. *BMC Biol.* **18**, 56 (2020).

2. González-Pech, R.A. et al. Comparison of 15 dinoflagellate genomes reveals extensive sequence and structural divergence in family Symbiodiniaceae and genus *Symbiodinium*. *BMC Biol.* **19**, 73 (2021).

3. Dougan, K.E. et al. Whole-genome duplication in an algal symbiont serendipitously confers thermal tolerance to corals. *bioRxiv*, 2022.2004.2010.487810 (2022).

4. Chen, Y. et al. Improved *Cladocopium goreaui* genome assembly reveals features of a facultative coral symbiont and the complex evolutionary history of dinoflagellate genes. *Microorganisms* **10**, 1662 (2022).

5. John, U. et al. An aerobic eukaryotic parasite with functional mitochondria that likely lacks a mitochondrial genome. *Sci. Adv.* **5**, eaav1110 (2019).

6. Beedessee, G. et al. Integrated omics unveil the secondary metabolic landscape of a basal dinoflagellate. *BMC Biol.* **18**, 139 (2020).

7. Farhat, S. et al. Rapid protein evolution, organellar reductions, and invasive intronic elements in the marine aerobic parasite dinoflagellate *Amoebophrya* spp. *BMC Biol.* **19**, 1 (2021).

8. Worden, A.Z. et al. Green evolution and dynamic adaptations revealed by genomes of the marine picoeukaryotes *Micromonas*. *Science* **324**, 268-272 (2009).

9. Huff, J.T., Zilberman, D. & Roy, S.W. Mechanism for DNA transposons to generate introns on genomic scales. *Nature* **538**, 533-536 (2016).

10. van der Burgt, A., Severing, E., De Wit, P.J.G.M. & Collemare, J. Birth of new spliceosomal introns in fungi by multiplication of introner-like elements. *Curr. Biol.* **22**, 1260-1265 (2012).

11. LaJeunesse, T.C. et al. Systematic revision of Symbiodiniaceae highlights the antiquity and diversity of coral endosymbionts. *Curr. Biol.* **28**, 2570-2580 (2018).

12. Dougan, K.E. et al. Genome-powered classification of microbial eukaryotes: focus on coral algal symbionts. *Trends Microbiol.* **30**, 831-840 (2022).

13. Stephens, T.G., Ragan, M.A., Bhattacharya, D. & Chan, C.X. Core genes in diverse dinoflagellate lineages include a wealth of conserved dark genes with unknown functions. *Sci. Rep.* **8**, 17175 (2018).

14. Bachvaroff, T.R. & Place, A.R. From stop to start: tandem gene arrangement, copy number and trans-splicing sites in the dinoflagellate *Amphidinium carterae*. *PLoS ONE* **3**, e2929 (2008).

15. Pachiadaki, M.G. et al. Charting the complexity of the marine microbiome through single-cell genomics. *Cell* **179**, 1623-1635 (2019).

16. Liao, Y., Smyth, G.K. & Shi, W. FeatureCounts: an efficient general purpose program for assigning sequence reads to genomic features. *Bioinformatics* **30**, 923-930 (2014).

17. Aramaki, T. et al. KofamKOALA: KEGG Ortholog assignment based on profile HMM and adaptive score threshold. *Bioinformatics* **36**, 2251-2252 (2020).

18. Slamovits, C.H. & Keeling, P.J. Widespread recycling of processed cDNAs in dinoflagellates. *Curr. Biol.* **18**, R550-552 (2008).

**a**

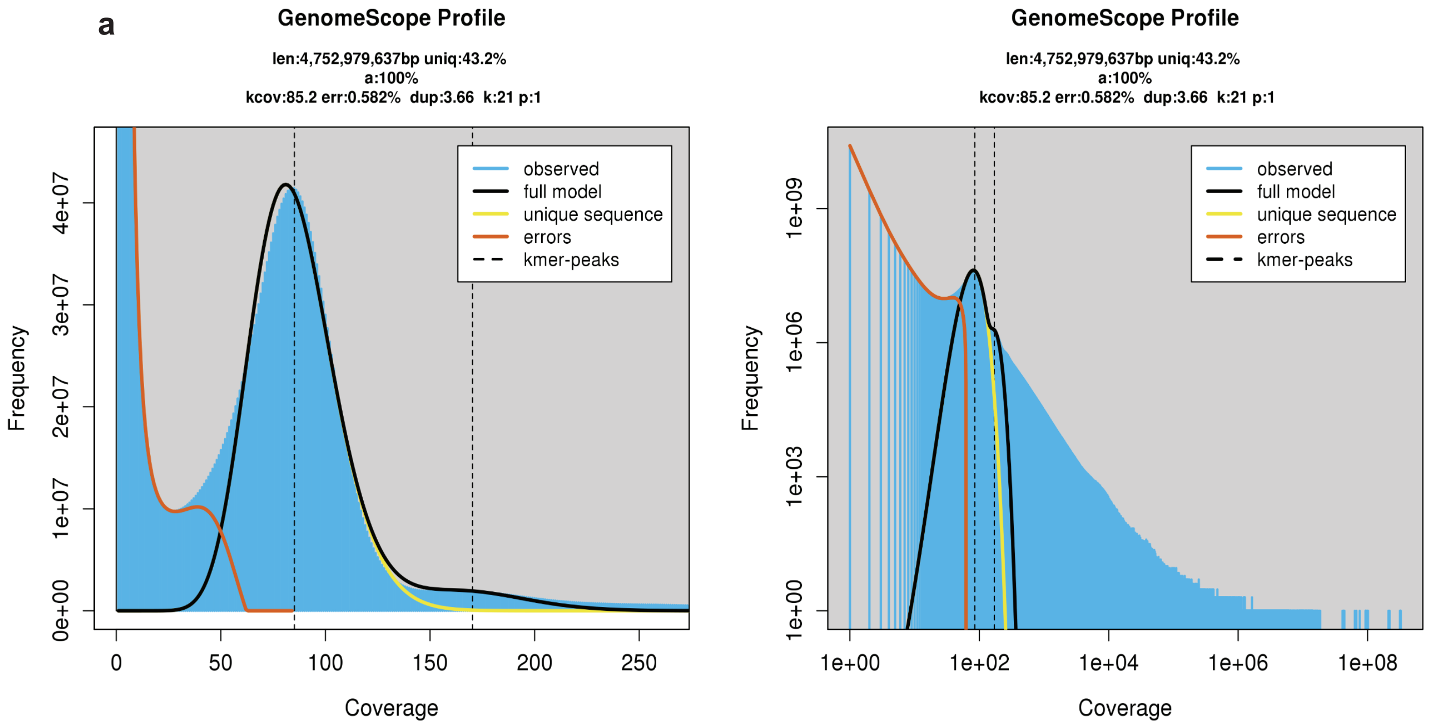


**b**

```
GenomeScope version 2.0
p = 1
k = 21

property                      min                 max
Homozygous (a)                100%                100%
Genome Haploid Length         4,730,452,501 bp    4,752,979,637 bp
Genome Repeat Length          2,685,886,986 bp    2,698,677,590 bp
Genome Unique Length          2,044,565,514 bp    2,054,302,047 bp
Model Fit                     42.9662%            90.8621%
Read Error Rate               0.582075%           0.582075%
```

**c**

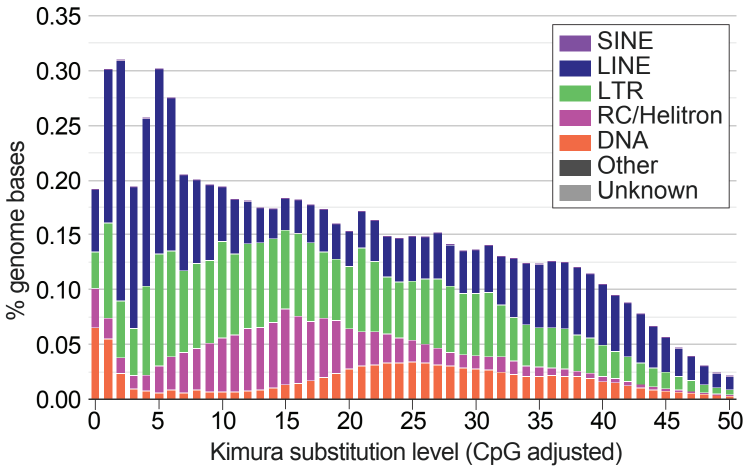


**Supplementary Fig. 1.** Analysis of *P. cordatum* genome based on (a) distribution of 21-mers indicating a haploid genome, (b) genome-size estimation based on 21-mers using GenomeScope2, and (c) repeat landscape excluding unknown repeats.

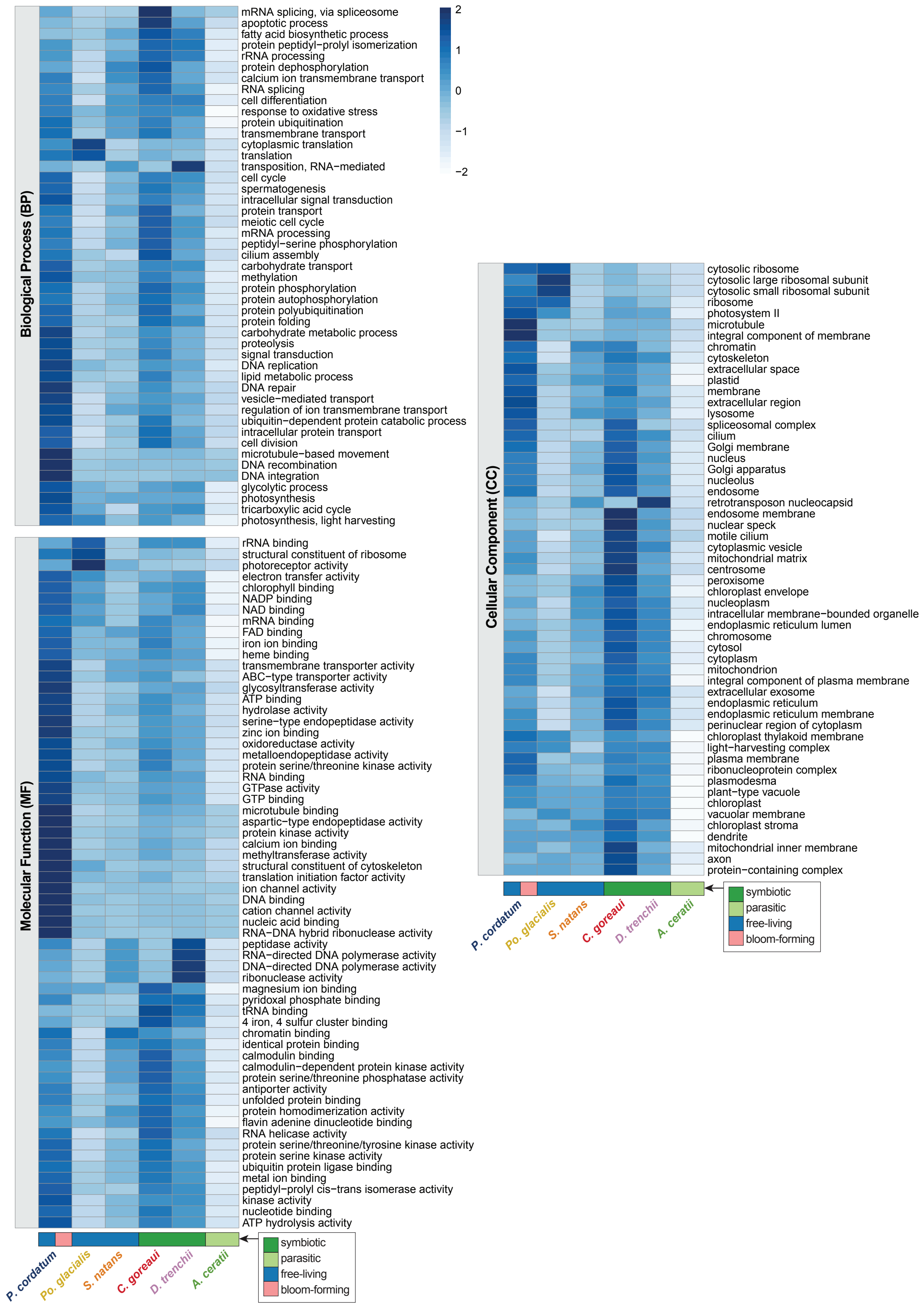


**Supplementary Fig. 2.** Gene functions encoded in the genome of *P. cordatum* and the other five representative taxa based on count of Gene Ontology (GO) terms.

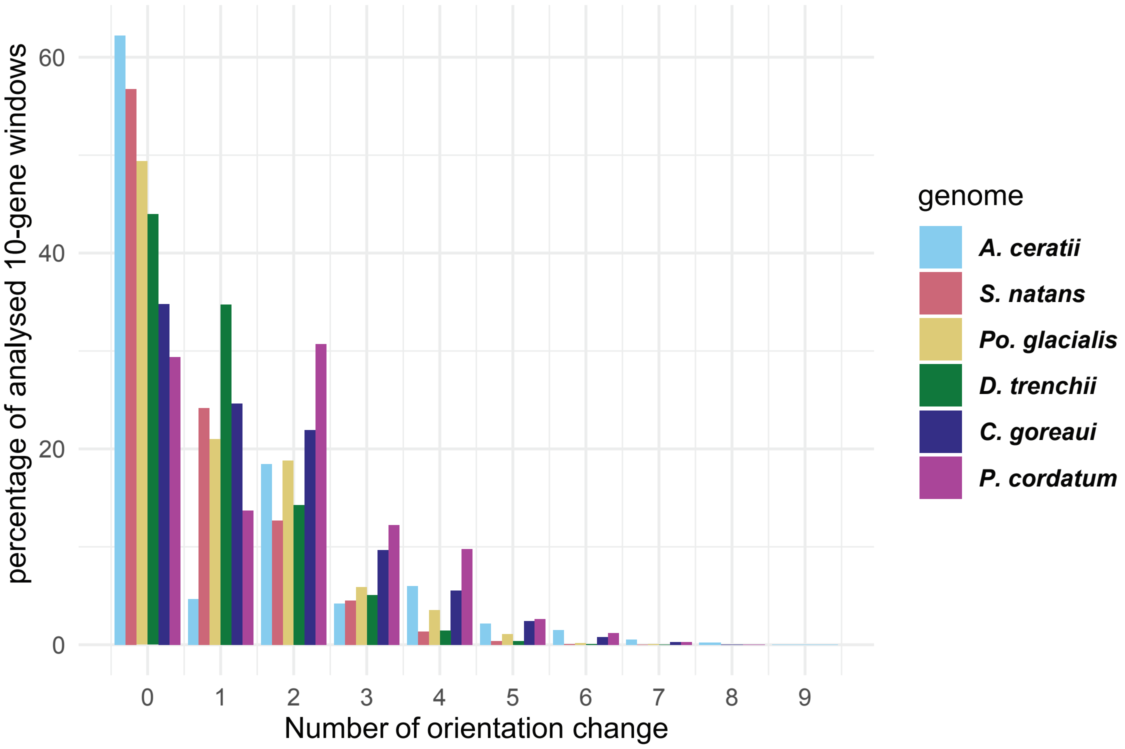


**Supplementary Fig. 3.** Unidirectional coding of genes in dinoflagellate genomes based on gene-orientation changes in ten-gene windows.

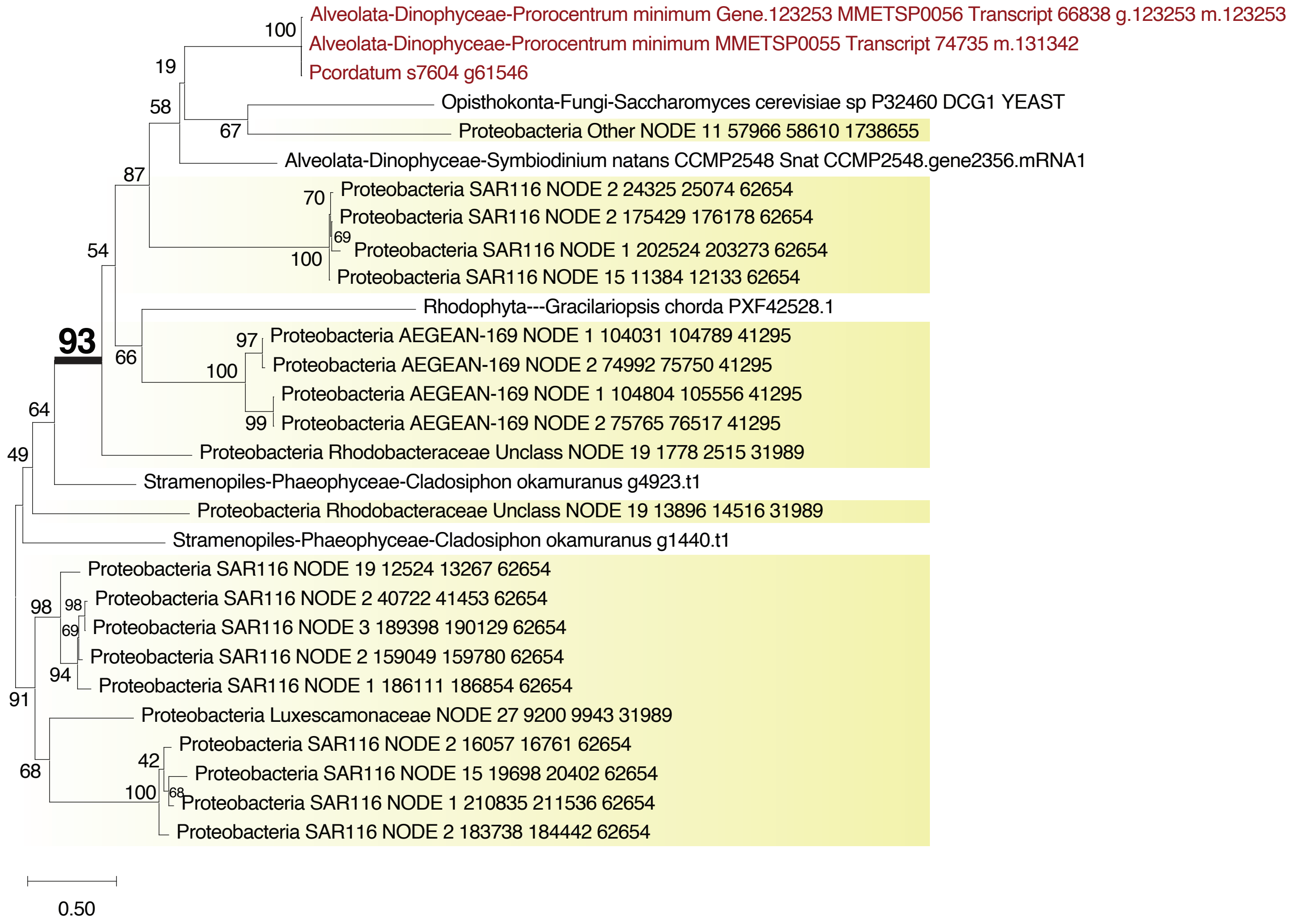


**Supplementary Fig. 4.** Maximum likelihood protein tree of a putative hydantoin racemase indicating bacterial origin of the encoding gene in *P. cordatum*.

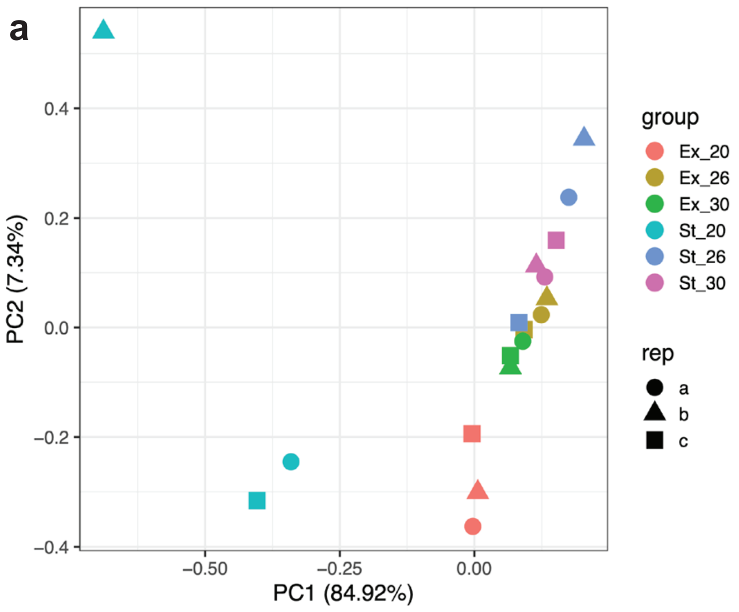


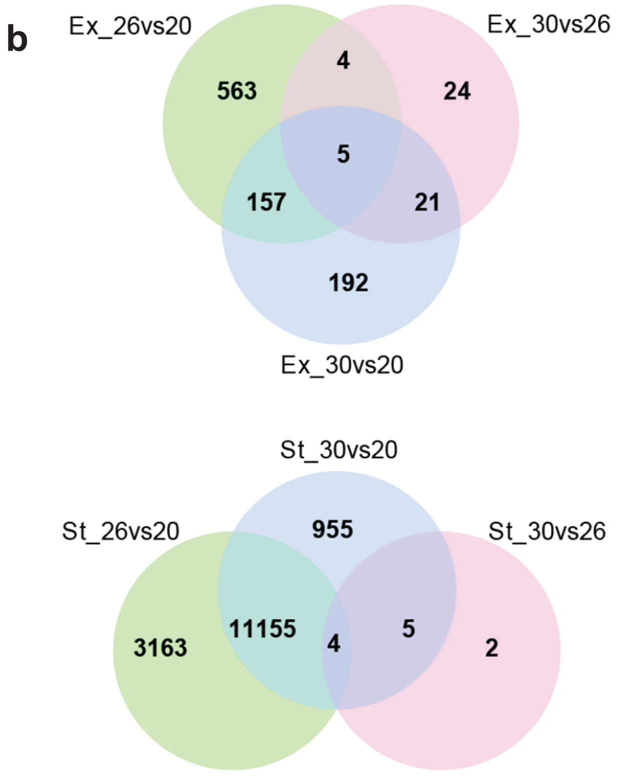


**Supplementary Fig. 5.** Transcriptome analysis of *P. cordatum* showing (a) principal component analysis of samples based on FPKM values, and (b) Venn diagram detailing number of DEGs between any two temperature conditions, independently for Ex and St phases.

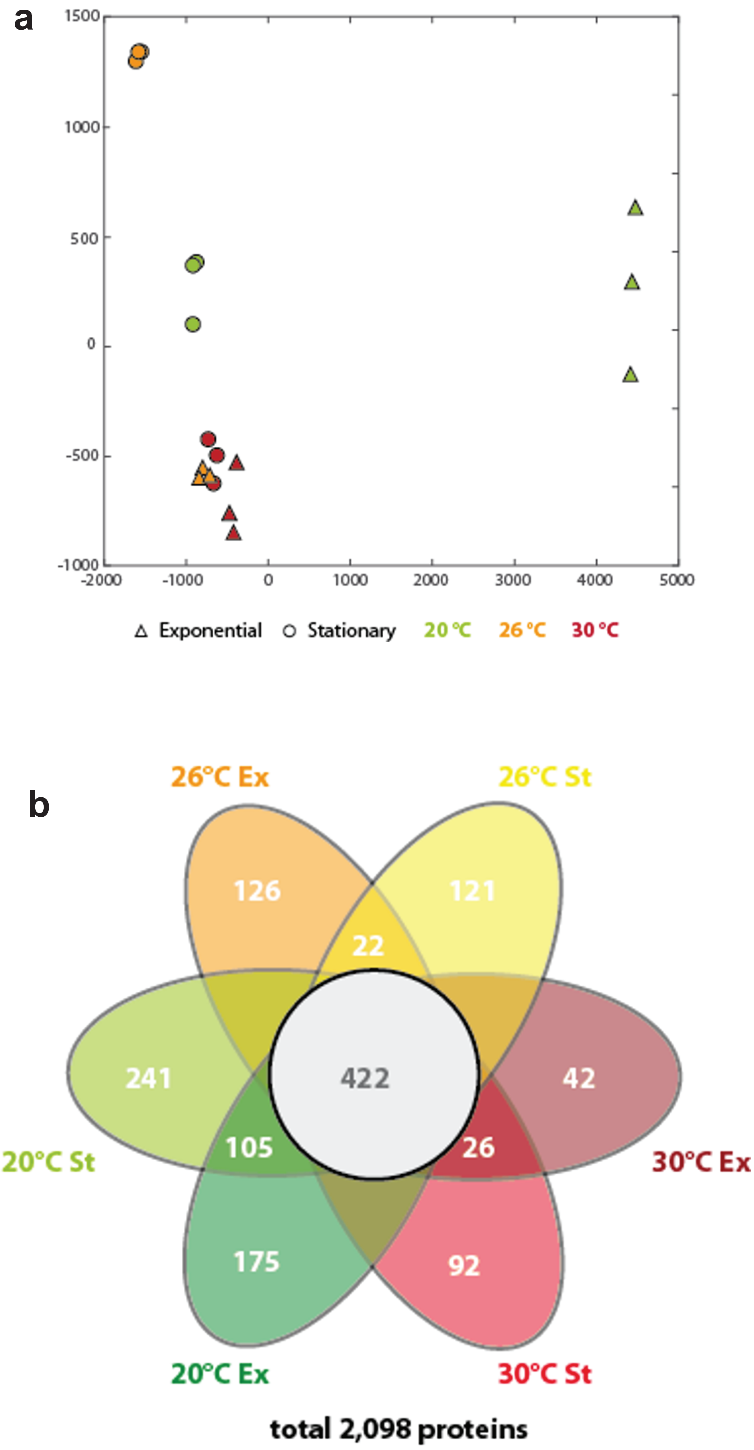


**Supplementary Fig. 6.** The proteome landscape of *P. cordatum* heat-stress response, showing (a) NMDS visualization of individual samples based on peptide count data (max error = $1.24 \times 10^{-11}$), and (b) Venn diagram of number of proteins in distinct conditions.

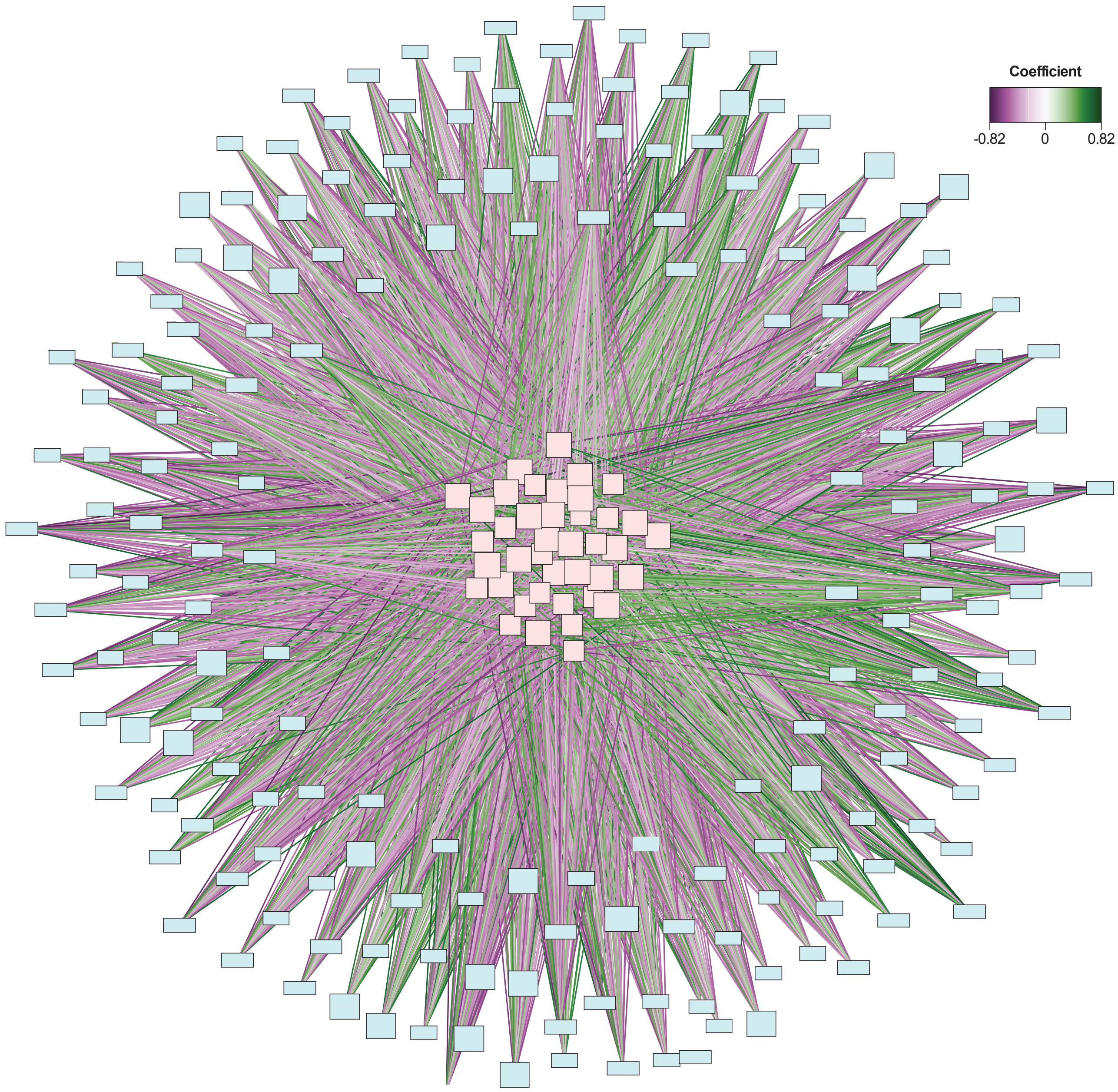


**Supplementary Fig. 7.** Network visualization of multi-omics signature inferred from both proteins and transcripts that represent a heat-stress response specific to 26°C and to 30°C. Colors of the lines indicate the nature of the relationships between proteins and transcripts, with positive relationships in green and negative relationships in purple.

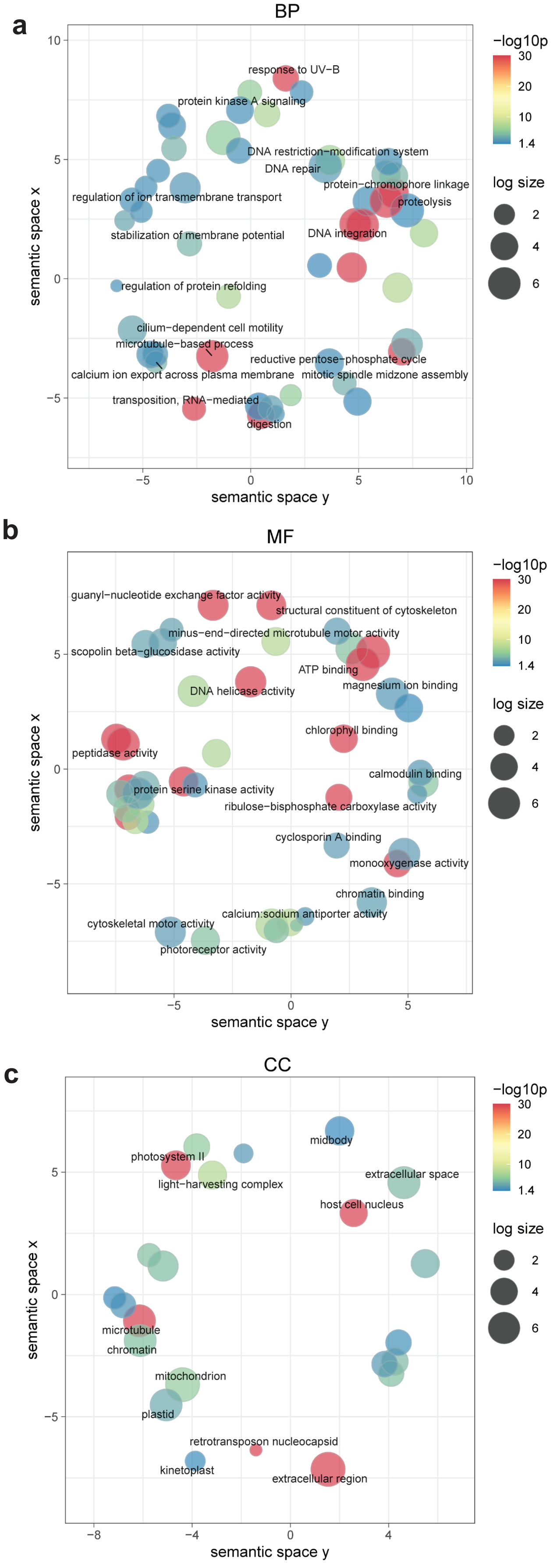


**Supplementary Fig. 8.** Functional enrichment of dispersed gene copies based on GO terms in homologous sets of size 20 or greater (total of 13,529 genes), shown for (a) Biological Process, (b) Molecular Function, and (c) Cellular Component.

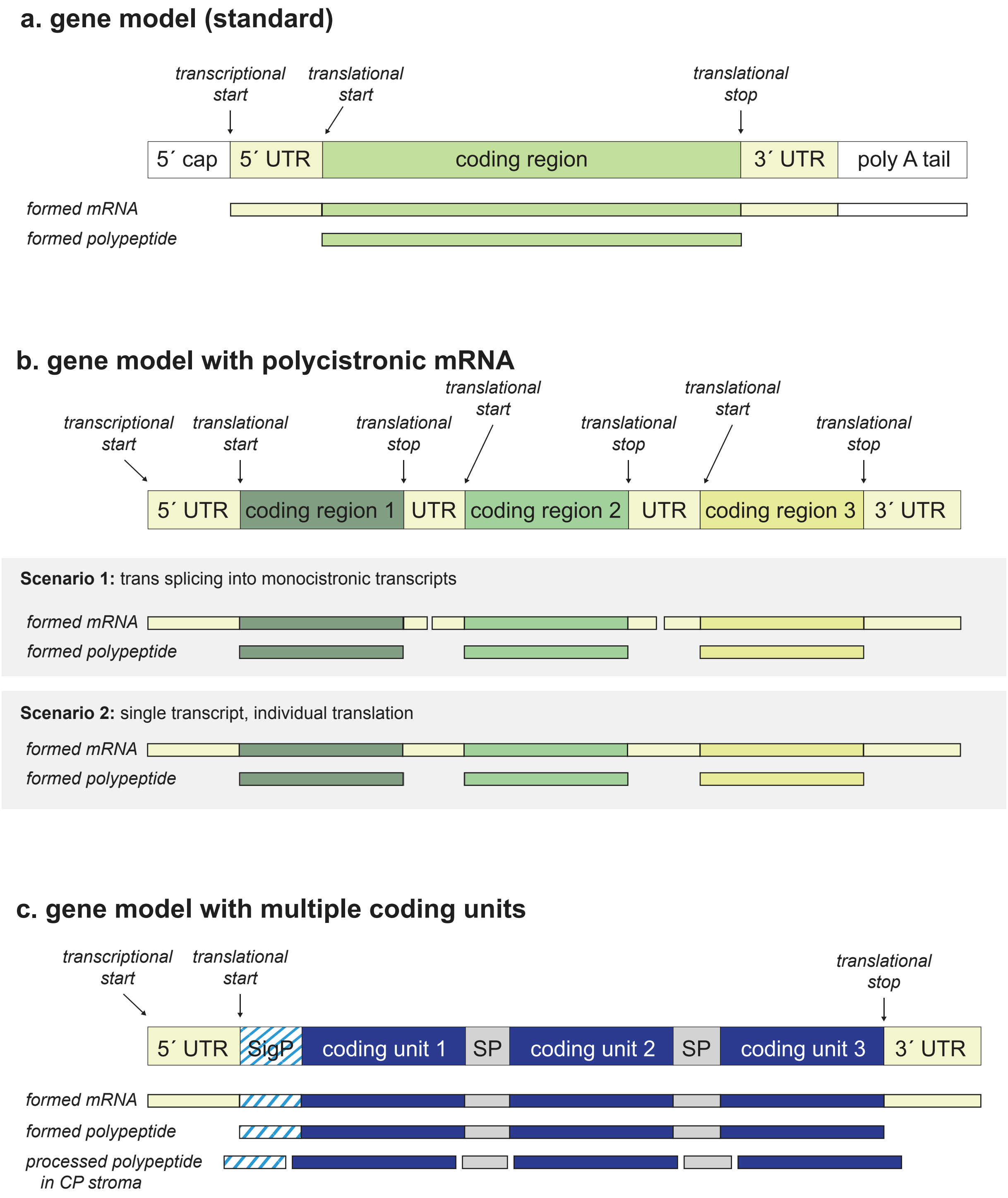


**Supplementary Fig. 9.** Schematic diagram three distinct gene models identified in *P. cordatum*: (a) standard gene model, (b) gene model with polycistronic mRNA, and (c) gene model with multiple coding units.

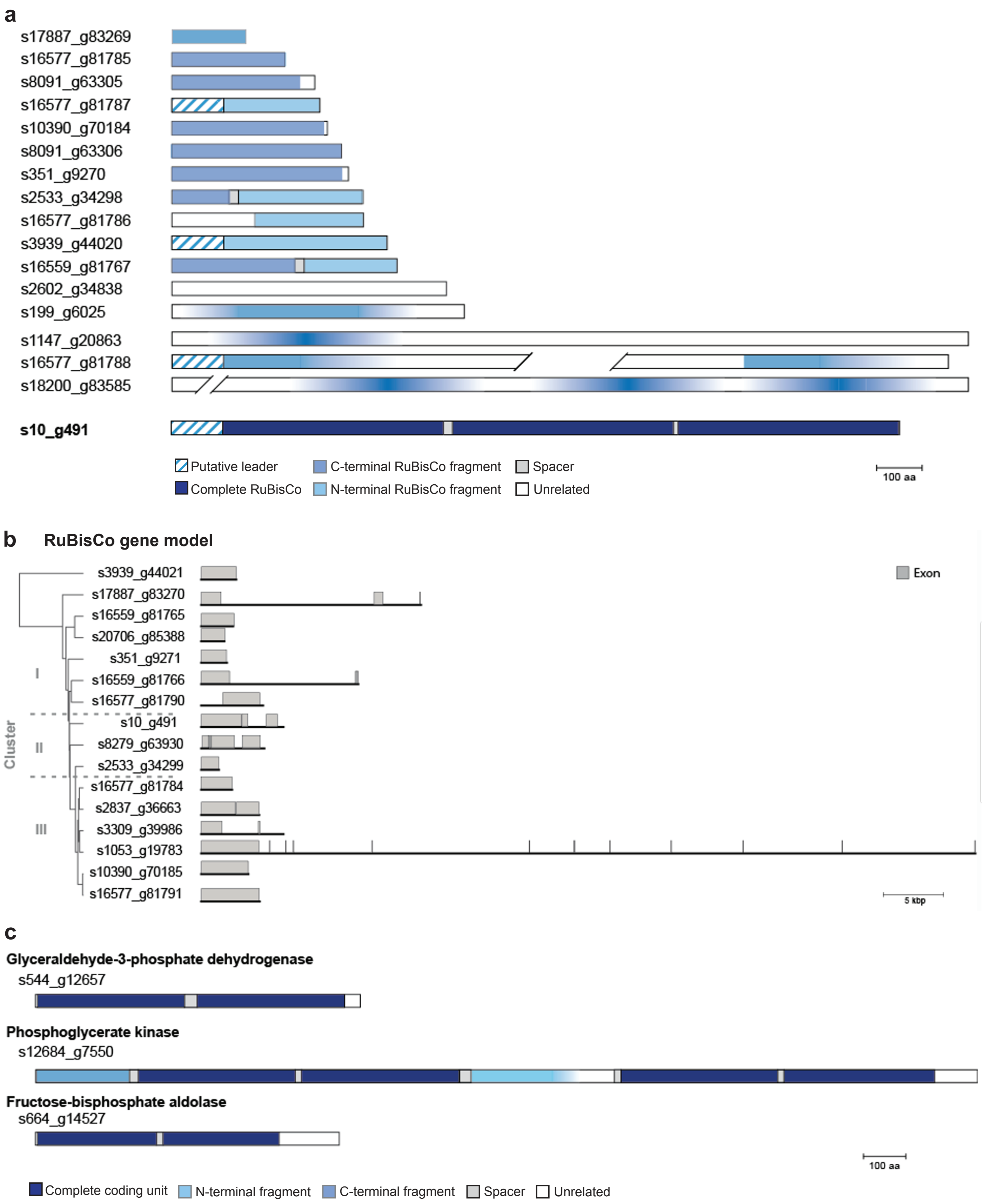


**Supplementary Fig. 10.** Detailed information on gene models encoding multiple coding units (CUs), showing the scale model for (a) those containing only fragments of RuBisCo CU with s10_g491 as a reference that encodes 3 CUs; (b) RuBisCo gene models and the corresponding exons; and (c) those containing multiple CUs that encode protein functions related to central metabolism.

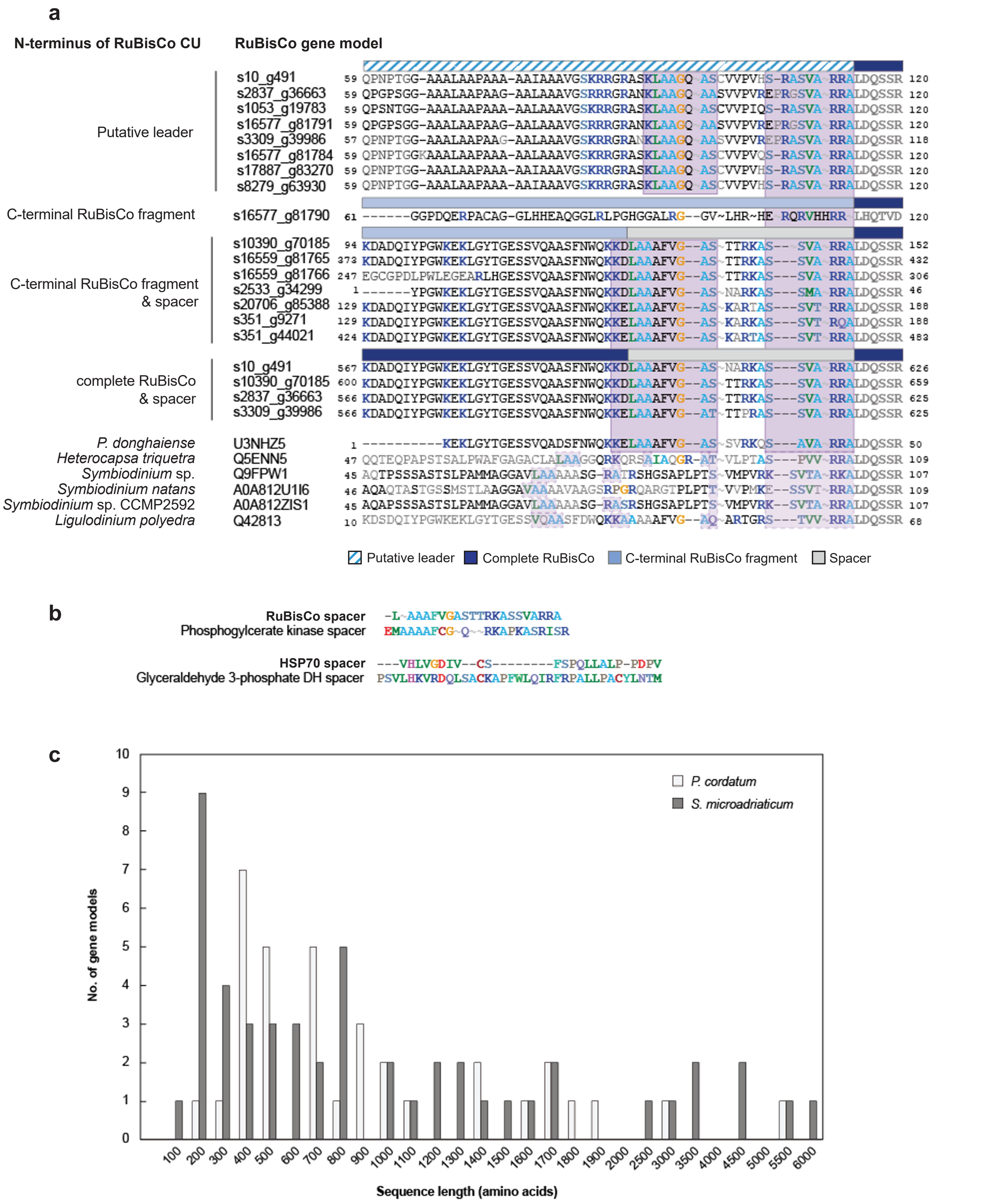


**Supplementary Fig. 11.** Comparison of RuBisCo gene models of *P. cordatum* and other dinoflagellate species, showing the (a) alignment of the predicted leader sequences, N-terminal sequence of the first CU, and the spacer region between two complete CUs; (b) alignment of spacer regions of chloroplast-localized proteins (RuBisCo and phosphoglycerate kinase) and non-chloroplastic proteins (HSPv70 and glyceraldehyde 3-phosphate dehydrogenase); and (c) histogram of Rubisco genes encoded in the genomes of *P. cordatum* and *S. microadriaticum* relative to protein-sequence length.
